# Mechano-Piezoelectric F-Peptide Hydrogels Enable *In Situ* Stem Cell and Immune Niche Modulation for Cartilage Regeneration

**DOI:** 10.64898/2026.08.18.745416

**Authors:** Xuekai Song, Zhaowen Xu, Siqi Zhang, Tongtong Zhang, Can Liu, Hongyi Huang, Yonghui Hu, Mengjie Yang, Lixiang Zhao, Yingzi Zhang, Rui Wang, Kuan Hu

## Abstract

Osteoarthritis, characterized by cartilage degradation and synovial inflammation, has spurred interest in mechano-piezoelectric bio-hydrogel therapies that can both relieve symptoms and reverse progression. However, current approaches lack sufficient piezoelectric output and dual cartilage/inflammation targeting. To address this, we demonstrated a mechano-piezoelectric peptide hydrogel composed of nanofibers integrating piezoelectric cues with mesenchymal stromal cells (MSCs) recruitment and PIEZO2 mechanosignaling. Molecularly, the hydrogel’s seed peptide incorporated four functions: COL2A1 targeting, MMP-13 responsiveness, MSCs homing, and self-assembly. Overexpressed MMP-13 in the osteoarthritis niche triggers gelation, promoting MSCs recruitment and drug retention. Fluorination modulates hierarchical nanofiber assembly, enhancing mechanical and piezoelectric properties, as confirmed by morphological, biophysical, and computational analysis. The trifluoromethyl-modified, 4-octyl itaconate (4–OI) loaded formulation reverses osteoarthritis via PI3K/AKT activation and Wnt/β–catenin suppression, as shown by improved Osteoarthritis Research Society International (OARSI) scores, bone microarchitecture, and cartilage matrix. This synergy of mechano–piezoelectric cues and 4–OI offers a clinically promising strategy for osteoarthritis.

## Main

Osteoarthritis (OA) is a prevalent degenerative joint disease driven by aging, population growth, and obesity, affecting hundreds of millions worldwide^1, 2, 3^. Current pharmacological treatments including nonsteroidal anti–inflammatory drugs (NSAIDs), glucocorticoids, and hyaluronic acid, remain palliative, failing to address underlying cartilage degeneration; and are limited by systemic adverse effects and short intra–articular retention^4, 5, 6, 7^. Electrical stimulation holds promise for tissue regeneration, yet exogenous implementation suffers from signal attenuation by tissue barriers and infection/foreign-body responses from implantable devices^8, 9^. Biocompatible, biodegradable piezoelectric materials that generate endogenous electric signals upon mechanical deformation offer an alternative. Piezoelectric biomaterials have been extensively explored for tissue repair, as exemplified by Zhou Li’s group works on injectable piezoelectric hydrogels and electrospun fiber scaffolds^10, 11^. Of particular relevance to our work, healthy cartilage itself exhibits intrinsic piezoelectricity; compressive deformation produces streaming and piezoelectric potentials^12, 13^, providing a biomimetic rationale for using such materials to recapitulate native mechano–electrical transduction. Mechanistically, local electric signals drive Ca²⁺ influx, activate p38 mitogen-activated protein kinase (MAPK) and integrin/ focal adhesion kinase (FAK) pathways to promote chondrogenic differentiation and MSCs migration^14, 15^, while piezo-type mechanosensitive ion channel component 1/2 (PIEZO1/2) and phosphatidylinositol 3-kinase/ protein kinase B (PI3K/AKT) may also contribute, though their precise roles remain unclear^16^. Inspired by these insights, mechano–piezoelectric hydrogels that produce on–demand electric signals under mechanical loading have attracted substantial interest^17^. However, existing systems face considerable limitations: inorganic nanoparticle–doped hydrogels (*e.g.*, BaTiO_3_) offer high output but are non-degradable, risking chronic inflammation^18^; synthetic polymeric materials (*e.g.*, PVDF) have good performance but suffer from hydrophobicity and lack of bioactive sites, hindering functionalization and cell–specific modulation^19^. Peptide–based piezoelectric hydrogels overcome these limitations through modular functionalization (*e.g.*, targeting ligands, enzyme–responsive sequences, cell–recruiting peptides), cost-effective solid-phase synthesis, and inherent biodegradability into non–toxic amino acids. Since the discovery of shear piezoelectricity in self–assembled Fmoc-diphenylalanine^20^, peptide materials have opened new avenues for piezoelectric biomedical applications. For instance, Stupp and co–workers recently reported a flexible, biodegradable composite of peptide–amphiphile and PVDF that can be wirelessly activated by ultrasound for neuronal stimulation^21^. Nevertheless, conventional unmodified peptides still exhibit insufficient piezoelectric output for *in vivo* therapeutic applications^22^. Fluorination effectively enhances peptide piezoelectricity. For example, monofluorination increases the piezoelectric strain coefficient of Cbz–Phe(4F)^23^, and fluorinated diphenylalanine nanotubes show over ten–fold higher piezoelectric performance than unmodified ones^24^. The high electronegativity of fluorine strengthens π-π interactions and structural stability, synergistically boosting piezoelectric properties^23, 25, 26^.

To address the above-mentioned bottlenecks, we designed a mechano–piezoelectric peptide hydrogel that integrates synergistic mechanical/piezoelectric stimulation, MSCs recruitment, and macrophage metabolic reprogramming. The innovation lies in first combining fluorination-enhanced piezoelectricity with multiple biological functions in one platform: active cartilage targeting (COL2A1–binding sequence)^27^, microenvironment–triggered self–assembly (MMP-13-responsive linker)^28, 29, 30^, and stem cell homing (SKP peptide)^31^. This transcends the “single-function” paradigm, enabling comprehensive intervention: inflammation regulation, stem cell recruitment, chondrogenesis, and matrix remodeling are all addressed in one integrated process. We synthesized varied fluorinated nanofibers and, via all-atom (AA-) and coarse-grained molecular dynamics (CG-MD) simulations, elucidated that fluorine substituents modulate π-π interactions and secondary structure, governing nanomorphology, stability, and signal outputs. Trifluoromethyl modification conferred the greatest improvements in stability and piezoelectric output. At the cellular level, the hydrogel promoted MSCs adhesion, migration, and chondrogenic differentiation; in the destabilization of the medial meniscus (DMM) rat models, it induced macrophage metabolic reprogramming and significantly recruited endogenous MSCs to cartilage defects, synergistically promoting matrix regeneration. We anticipated that the trifluoromethyl–modified, 4-OI-loaded formulation could reverse OA via PI3K/AKT activation and wingless–related integration site/beta–catenin (Wnt/β–catenin), improving Osteoarthritis Research Society International (OARSI)/Mankin scores, bone microarchitecture (bone volume/total volume *i.e.* BV/TV, osteophytes), and cartilage matrix (COL2A1, IL–1β), thereby offering a clinically promising strategy through mechano–piezoelectric/4–OI synergy. Collectively, this work presents a clinically translatable mechano-piezoelectric hydrogel that synergizes piezoelectric cues with 4–OI–mediated immunomodulation, holding promise as a next–generation, surgery–sparing strategy for OA and other degenerative joint disorders.

### Fluorination modulates self-assembled peptide hydrogels to enhance nanofiber density, BMSCs recruitment, and chondrogenic differentiation

To preliminarily investigate the effect of aromatic ring fluorination on π-π interactions and self-assembled hydrogel architecture, we synthesized four SKP-based fluorinated peptides (F-peptides): SKP-1 (unmodified), SKP-2 (monofluoro), SKP-3 (trifluoro) and SKP-4 (trifluoromethyl) that form hydrogels capable of recruiting bone marrow mesenchymal stem cells (BMSCs) and supporting chondrogenic differentiation (**Fig. 1a**). All peptides used in this study were purified by HPLC and verified by ESI-MS, with purities exceeding 90% (**Supplementary Fig. 1-19**).

**Fig. 1.**
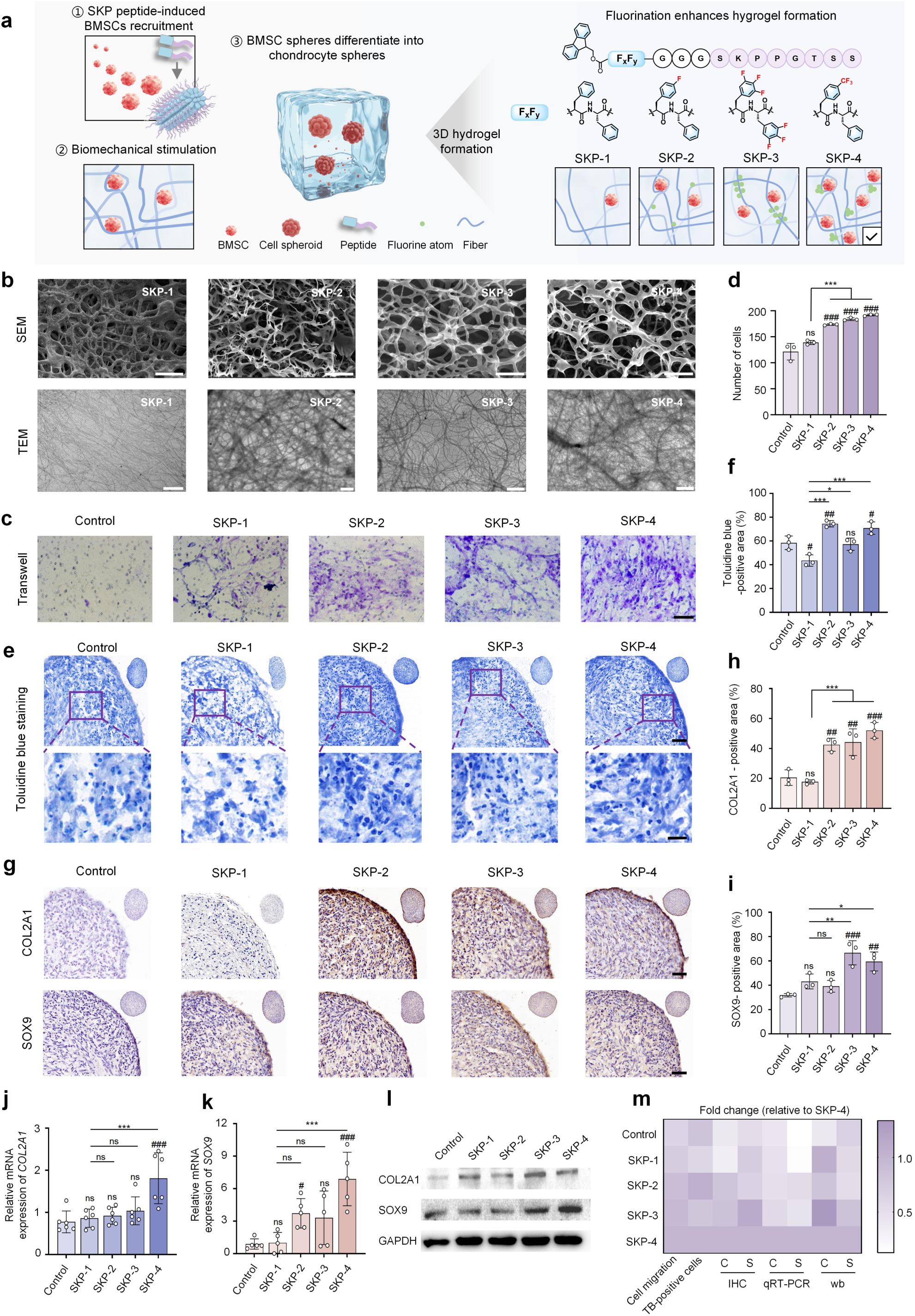
Fluorinated SKP peptide hydrogels recruit BMSCs and promote chondrogenesis. **a**, Schematic diagram of four SKP peptides’ structure, hydrogel formation, and BMSCs recruitment. **b**, SEM and TEM images of the different SKP series hydrogels. Scale bars: 200 μm (SEM, SKP-1), 20 μm (SEM, SKP-2), 1 μm (SEM, SKP-3), 2 μm (SEM, SKP-4), and 2 μm (TEM). **c**, Representative images of the lower surface of transwell membranes stained with crystal violet after 18 h of culture, scale bar: 200 μm. **d**, Quantification of crystal violet-stained cells in the transwell assay. **e**, Toluidine blue staining of BMSC spheroids after 14 days of culture in medium or with SKP-1/2/3/4 hydrogels. Scale bar: 50 μm / 20 μm (insets). **f**, Quantification of toluidine blue staining-positive area of BMSC spheroids with ImageJ. **g**, Representative COL2A1 and SOX9 immunohistochemistry staining of BMSC spheroids after 14-day culture with medium or four fluorinated hydrogels. Scale bar: 50 μm. Quantification of COL 2A1 (**h**) or SOX9 (**i**) -positive area of BMSC spheroids with ImageJ. **j**, qRT-PCR analysis of gene expression of *COL2A1* in BMSC spheroids, n = 6. **k**, qRT-PCR analysis of gene expression of *SOX9* in BMSC spheroids, n = 5. **l**, Western blot analysis of SOX9 and COL2A1 protein levels in BMSC spheroids. **m**, Heatmap summarizing the relative effects of fluorinated hydrogels on BMSC recruitment and differentiation versus the SKP-4 group. C means COL2A1; S means SOX9. Statistical significance was determined by one-way ANOVA, with multiple comparisons. Data are all presented as mean ± SD. For significance symbols: *p < 0.05, **p < 0.01, ***p < 0.001 versus SKP-1; #p < 0.05, ##p < 0.01, ###p < 0.001 versus control groups. ns, not significant.

Tetraphenylethylene (TPE)-conjugated SKP-3 exhibited aggregation-induced emission with increasing concentration, confirming molecular assembly; scanning electron microscopy (SEM) and transmission electron microscopy (TEM) at 10 mg·mL^-1^ further validated the presence of a distinct hydrogel nanofibrillar network. (**Supplementary Fig. 20a-c**). All four peptides formed stable hydrogels at 10 mg·mL^-1^ in ddH2O or PBS (**Supplementary Fig. 20d**). SEM and TEM revealed nanofibrous networks for all, with SKP-4 showing denser structure and smaller pore sizes (**Fig. 1b and Supplementary Fig. 20e**) Transwell assay revealed that SKP-2/3/4 hydrogels significantly enhanced BMSCs migration compared to SKP-1 or Control groups (**Fig. 1c, d**). Chondrogenic differentiation was assessed after 14-days culture in the presence of different fluorinated hydrogels or only medium supplemented with transforming growth factor-β (TGF-β). Toluidine blue staining, immunohistochemistry (IHC) of collagen type Ⅱ alpha 1 chain (COL2A1) and sex-determining region SRY-box transcription factor 9 (SOX9), qRT-PCR and Western blot all demonstrated that SKP-4 markedly upregulated both chondrogenic markers, while SKP-2 and SKP-3 exhibited moderate or selective effects (Fig 1e-l). Taking all these factors into consideration, a summary heatmap confirmed that SKP-4 most effectively promoted both recruitment and chondrogenic differentiation (**Fig. 1m**).

Collectively, our findings demonstrated that fluorination, particularly the trifluoromethyl group enhances fiber density, likely via electron-withdrawing and steric effects of the trifluoromethyl group. Interestingly, the divergent performance of SKP-3, which enhanced cell migration but failed to induce significant chondrogenic differentiation, suggesting that the fluorination pattern critically influences biological outcomes, possibly through alternations in hydrogel stiffness, nanostructure, or cell-signaling interactions. Overall, SKP-4 emerges as the most promising candidate for cartilage tissue repair.

### From SKP to CMP: targeting and enzyme-responsive design

The extracellular matrix (ECM) in osteoarthritis undergoes characteristic pathological alterations, including local microenvironment acidification^32, 33^ and significant upregulation of matrix metalloproteinase (MMPs), particularly of matrix metalloproteinase-13 (MMP-13, collagenase-3)^34^. These changes collectively create a destructive microenvironment: acidic conditions directly impair chondrocyte function and further activate matrix-degrading enzymes, while MMP-13 overexpression leads to abnormal degradation of key matrix components, disrupting the balance of cartilage matrix synthesis and degradation. We first evaluated the cartilage retention of fluorinated SKP peptides on knee joint sections from anterior cruciate ligament transection (ACLT) rats. Immunofluorescence staining revealed that Cy5-SKP-1/4 peptides exhibited limited cartilage uptake and retention, with negligible co-localization with COL2A1 (**Fig. 2a**). Though Cy5-SKP-4 showed greater cartilage permeability than Cy5-SKP-1, likely due to the trifluoromethyl modification enhancing lipophilicity, no apparent COL2A1 targeting was observed for either SKP peptide, suggesting that fluorination alone is insufficient for effective cartilage targeting. To address this, we designed a smart delivery system, COL2A1 targeting and MMP-13-responsive peptides (CMPs), by introducing an MMP-13-responsive sequence at the C-terminus of the SKP peptide^28, 30^, followed by a COL2A1-targeting sequence^27^. This modification enables targeted delivery to knee cartilage sites, where MMP-13-mediated cleavage triggers *in situ* SKPG self-assembly and on-demand hydrogel formation (**Fig. 2b**).

**Fig. 2.**
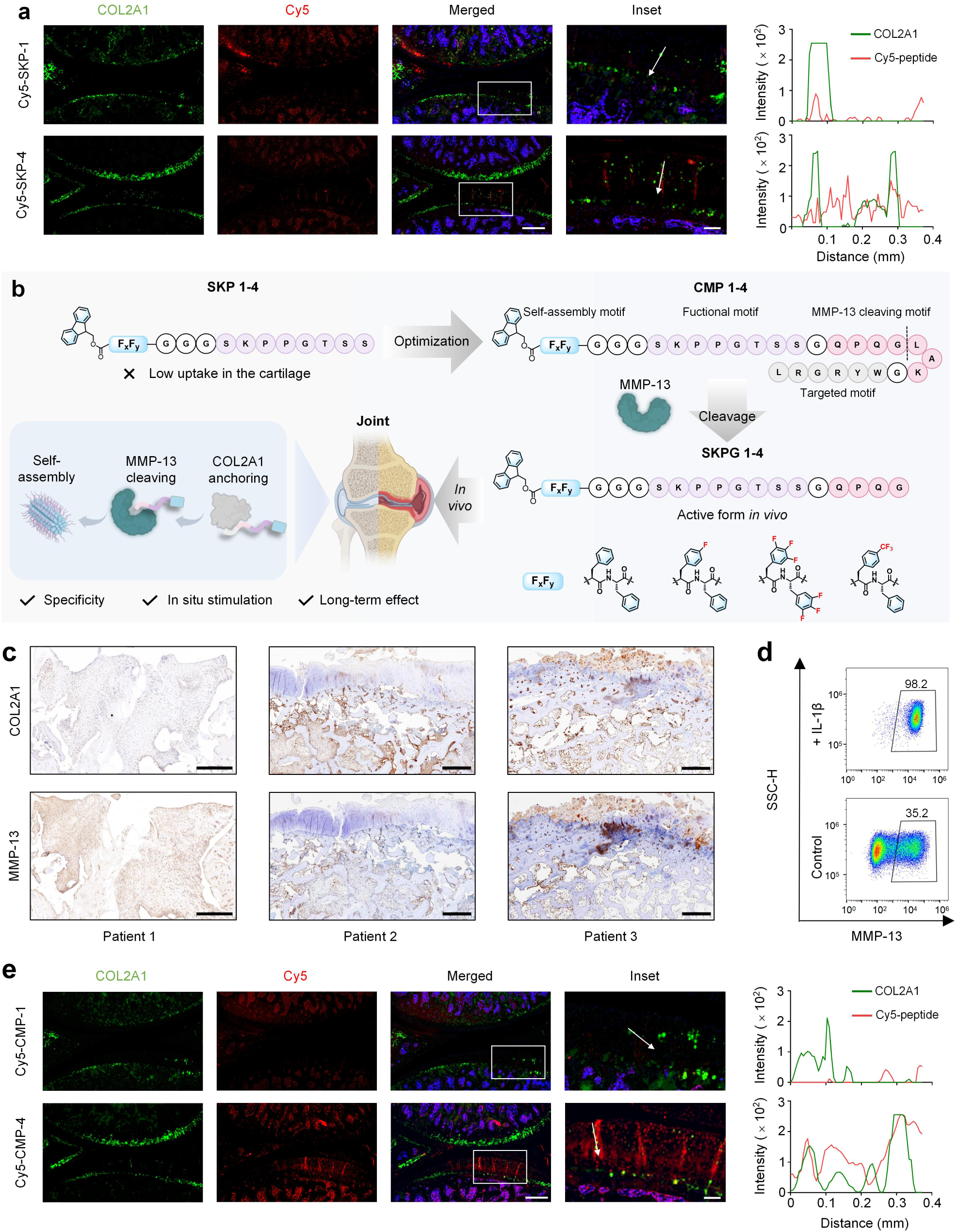
Design and validation of CMPs engineered from SKP with targeting and MMP-13-responsive cleavage. **a**, Fluorescence imaging and co-localization analysis of Cy5-SKP-1/4 (red) with COL2A1 (green)in the knee joints of rats at 8 weeks after ACLT surgery. Nuclei were stained with DAPI (blue). Scale bar: 0.5 mm, 0.1 mm (insets). **b**, Schematic of CMPs design optimized from SKP, featuring COL2A1 targeting and MMP-13-responsive cleavage, followed by *in situ* SKPG self-assembly. **c**, Immunohistochemistry of COL2A1 and MMP-13 of knee joint tissue samples from three patients with severe osteoarthritis. Scale bar: 0.5 mm (patient 1), 1 mm (patient 2 and 3). **d**, Flow cytometry analysis of MMP-13 expression in human chondrocytes before and after IL-1β stimulation. **e**, Fluorescence imaging and co-localization analysis of Cy5-CMP-1/4 (red) with COL2A1 (green) in the knee joints of rats at 8 weeks after ACLT surgery. Nuclei were stained with DAPI (blue). Scale bar: 0.5 mm, 0.1 mm (insets).

IHC analysis of cartilage tissue from three patients with severe knee OA (KOA) who underwent knee replacement surgery revealed high expression levels of both MMP-13 and COL2A1 (**Fig. 2c**). Human chondrocytes induced with IL-1β for 2 weeks to mimic the OA phenotype exhibited significantly higher MMP-13 expression than normal cells, as determined by flow cytometry (**Fig. 2d**).

Having confirmed the pathological relevance of MMP-13 and COL2A1 in OA, we next validated the cartilage-specific targeting of the CMP peptides. Notably, Cy5-CMP-4 showed superior cartilage penetration and pronounced overlapping signals with COL2A1, whereas Cy5-CMP-1 exhibited minimal co-localization (**Fig. 2e**). Quantitative analysis of Cy5 fluorescence signal in cartilage tissue further confirmed that the retention of Cy5-CMP-4 was significantly higher than that of the other groups (**Supplementary Fig. 21**). These results are consistent with our design rationale that the incorporation of both the COL2A1-targeting sequence and the fluorinated moiety within CMP-4 synergistically enhances cartilage targeting and retention. The MMP-13-responsive cleavage of the peptide sequence was further confirmed using a fluorescence resonance energy transfer (FRET) assay. FITC and 4-[[4-(dimethylamino)phenyl]diazenyl]benzoic acid (Dabcyl) were attached to the peptide termini; in the intact peptide, proximity of the donor and acceptor resulted in fluorescence quenching. Upon enzymatic cleavage, the fluorophores separated, enabling FITC fluorescence detection upon 488 nm excitation. The intact peptide showed no detectable fluorescence within 24 h, whereas fluorescence intensity at ∼525 nm increased after MMP-13 treatment, peaking before a slight decline, possibly due to reduced enzyme activity over time (**Supplementary Fig. 22a-c**). HPLC analysis after 24 h revealed near-complete digestion of the native peptide (retention time ∼15 min) and the appearance of two product peaks (∼12 min), confirming enzymatic cleavage (**Supplementary Fig. 22d, e**). Previous studies indicate that this sequence retains five amino acid residues based on the SKP peptide (*i.e.*, the SKPG peptide series) upon cleavage.

To further investigate the properties of the hydrogel formed after enzymatic cleavage, we designated the cleaved sequences that function effectively *in vivo* as SKPG-1/2/3/4 (**Fig. 3a**). As demonstrated above, fluorine-modified SKP peptide hydrogels could recruit MSCs and promoted chondrogenic differentiation (cartilage regeneration), addressing the source of chondrocytes. However, they did not attenuate the inflammatory phenotype in arthritis. To address inflammation, we encapsulated the anti-inflammatory agent 4-OI in four self–assembled peptide hydrogels: SKPG–1@4–OI (G1), SKPG–2@4–OI (G2), SKPG–3@4–OI (G3), and SKPG–4@4–OI (G4). 4–OI has been extensively reported to alleviate osteoarthritis by achieving metabolic reprogramming in macrophages^35, 36^ via activation of nuclear factor erythroid 2-related factor 2 (NRF2) through alkylation of kelch-like ECH-associated protein 1 (KEAP1) cysteine residues, thereby reducing IL–1β transcription^37^ (**Fig. 3b** and **Supplementary Fig. 23a**).

**Fig. 3.**
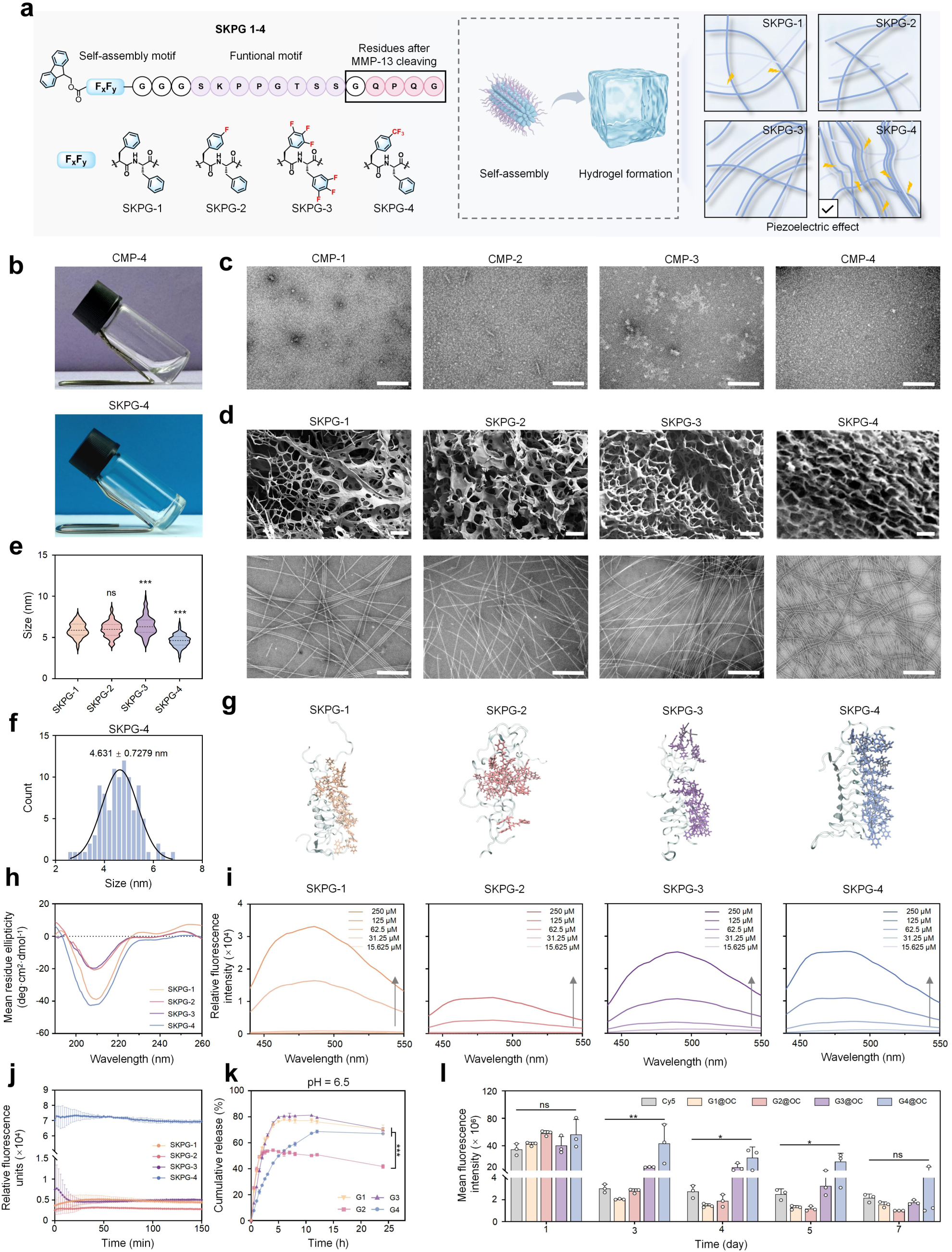
Fluorination modulates the self–assembly and drug release of SKPG hydrogels. **a**, Schematic illustration of the self–assembly of SKPG peptides into nanofibers with piezoelectric properties and subsequent hydrogel formation. **b**, Photographs of CMP-4 and SKPG-4 at 10 mg/mL in ddH_2_O. **c**, TEM images of the four CMP-series peptides. Scale bar: 200 nm. **d**, SEM and TEM images of the four SKPG-series self-assembling hydrogels. Scale bars: 50 μm (SEM) and 200 nm (TEM). **e**, Quantification of nanofiber diameters for the four SKPG peptide variants. Diameters were measured from TEM images using ImageJ (n = 100 per group). **f**, Diameter distribution histogram of SKPG–4 nanofibers. The solid line represents a Gaussian fit, indicating a normal distribution with a mean diameter of 4.631 ± 0.7279 nm (n = 100). **g**, Structure of the SKPG peptide decamers after 50 ns of AA-MD simulation. **h**, CD spectra of the SKPG-1/2/3/4 peptides. **i**, Self-assembly propensity of SKPG-1/2/3/4 probed by the 1,8-ANS assay. **j**, Comparison of β-sheet content in SKPG-1/2/3/4 assemblies using the ThT binding assay. **k**, Sustained release profiles of 4–OI from G1, G2, G3 and G4 in PBS buffer (pH 6.5), determined by UPLC–MS (n = 3). **l**, Quantification of mean fluorescence intensity in joint cavity after injection of Cy5, G1@OC, G2@OC, G3@OC and G4@OC. Data are all presented as mean ± SD. Statistical significance was determined by one–way ANOVA. *p < 0.05, **p < 0.01, ***p < 0.001; ns, not significant.

All four enzyme–responsive peptides retained self–assembly capacity and formed hydrogels with robust mechanical stability, with SKPG–4 showing the highest structural integrity. The CMPs did not assemble into appreciable nanofibers prior to enzymatic cleavage; however, upon cleavage, the SKPG series formed distinct supramolecular nanofiber structures, as visualized by SEM and TEM (**Fig. 3c, d**). Notably, SKPG–4 nanofibers were longer, denser and more flexible than those of SKPG–1 and SKPG–2, and diameter measurements further revealed that SKPG–4 fibers were significantly thinner, with a normal distribution and a median diameter of 4.631 ± 0.728 nm (**Fig. 3e, f** and **Supplementary Fig. 23b**). Chemical composition analysis by fourier transform infrared spectroscopy (FTIR) confirmed successful 4-OI loading in G1/2/3/4, as evidenced by the characteristic peaks of 4-OI superimposed on the peptide backbone (**Supplementary Fig. S23c**); energy dispersive spectroscopy (EDS) further verified the presence of C, N, O and F elements in G4(**Supplementary Fig. S23d**). Complementing these experimental findings, AA-MD simulations of decamers over 50 ns revealed that SKPG–4 adopted a more ordered and compact structure compared with the other analogs, consistent with the experimental observations (**Fig. 3g**). This was corroborated by circular dichroism spectroscopy, which revealed that all hydrogels adopted a β–sheet conformation, with SKPG–1 and SKPG–4 exhibiting more pronounced negative peaks, indicating higher β–sheet content and enhanced spatial regularity (**Fig. 3h**). Consistently, 8-anilino-1-naphthalenesulfonic acid (1,8–ANS) and Thioflavin T (ThT) assays further demonstrated that SKPG–4 possessed the strongest self–assembly propensity, with a pronounced blue shift and high fluorescence intensity even at 31.25 μM (**Fig. 3i, j**). Leveraging the superior structural features of G4, sustained–release experiments in PBS at pH 7.4 and 6.5 (mimicking the OA microenvironment) demonstrated that G4 exhibited the best performance, reaching peak concentration at approximately 12 h, about 7 h longer than the other groups, which might be attributed to the distinct fluorinated hydrogel structure (**Fig. 3k** and **Supplementary Fig. S23e**). Finally, *in vivo* retention studies using Cy5–encapsulated hydrogels in C57BL/6 mice revealed that G4@OC group (SKPG–4 encapsulating 4–OI and Cy5; G1@OC, G2@OC, and G3@OC denote SKPG–1, SKPG–2, and SKPG–3 encapsulating the same payload, respectively) group maintained the highest average fluorescence intensity in the knee joint from day 1 to day 7 (**Fig. 3l** and **Supplementary Fig. 24a, b**).

### Fluorination tunes self-assembly and mechano-piezoelectricity of SKPG

To elucidate the structural and functional basis of this piezoelectric enhancement, we selected SKPG-1 (non–fluorinated) and SKPG-4 (optimal) for comparative analysis (**Fig. 4a-o**). Atomic Force Microscopy (AFM) height sensor images confirmed that both peptides formed well-defined nanofiber morphologies, indicating that fluorination did not compromise the intrinsic self-assembly capability of the peptide (**Fig. 4b, c**). Piezoresponse force microscopy (PFM) revealed that, in stark contrast to SKPG-1, SKPG-4 displayed a classic butterfly-shaped amplitude loop and a distinct piezoelectric hysteresis loop, both indicative of superior piezoelectric performance (**Fig. 4d-g**). Quantitatively, the effective piezoelectric coefficient (d_33_) of SKPG-4 nanofibers reached 404.92 pm/V, approximately 15-fold higher than that of SKPG-1 (26.13 pm/V) (**Fig. 4h, i**). This striking enhancement suggests that trifluoromethyl modification dramatically amplifies the piezoelectric output of peptide nanofibers, positioning SKPG-4 as a potent mechano-active biomaterial.

**Fig. 4.**
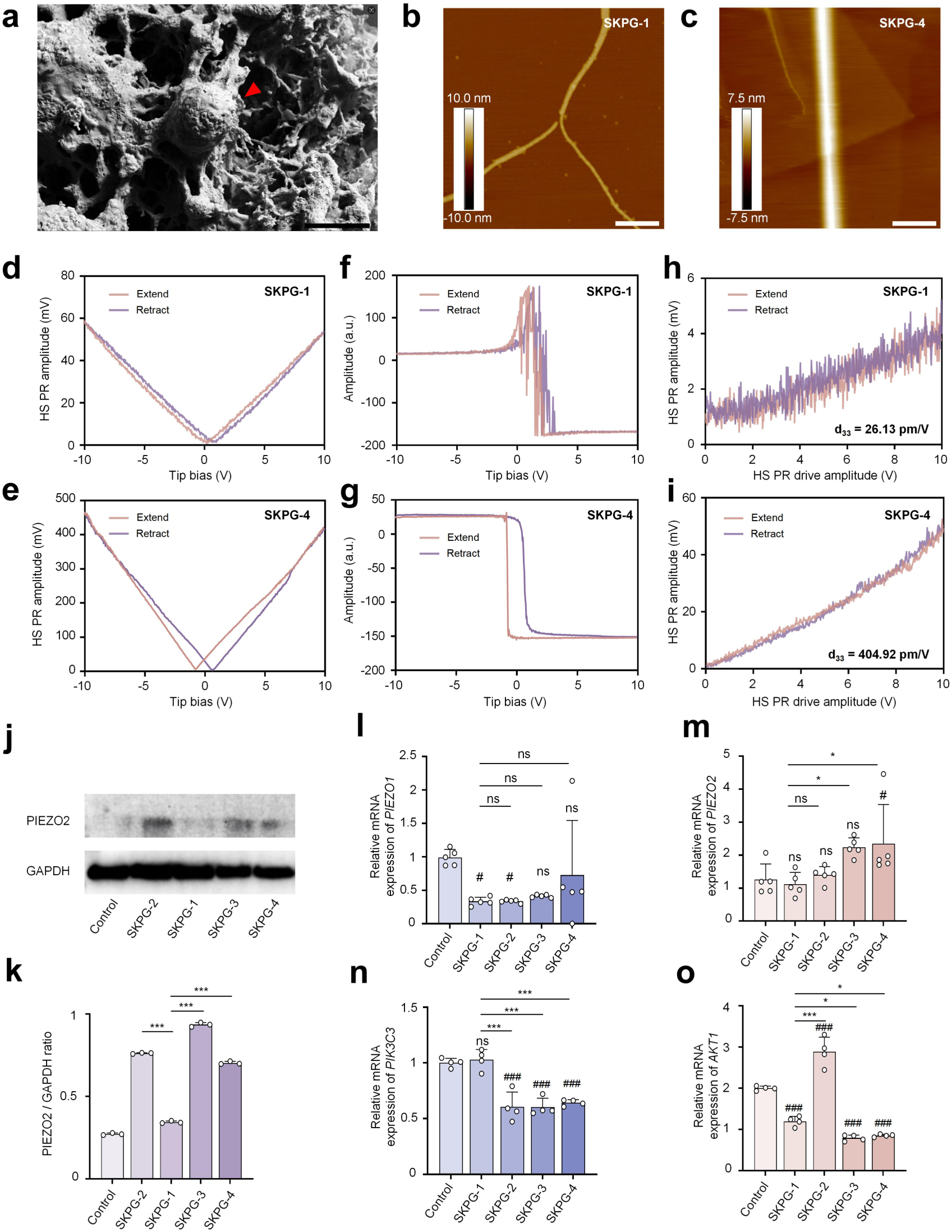
Fluorinated SKPG–4 hydrogels exhibit mechano–piezoelectric properties and activate PIEZO2–mediated mechanosignaling in BMSCs. **a**, SEM image of BMSCs adhered to the surface of SKPG-4 hydrogel. Scale bar: 40 μm. AFM height sensor images of SKPG–1 (**b**) and SKPG–4 (**c**) nanofibers. Scale bars: 200 nm. PFM amplitude loops of SKPG–1 (**d**) and SKPG–4 (**e**) nanofibers. PFM phase hysteresis loops of SKPG–1 (**f**) and SKPG–4 (**g**) nanofibers. Linear fits of piezoresponse amplitude as a function of drive voltage, from which the effective piezoelectric coefficient (d_33_) of SKPG–1 (**h**) and SKPG–4 (**i**) was derived. Western blot (**j**) and corresponding quantification (**k**) of PIEZO2 protein expression in BMSCs treated with different hydrogels. **l**-**o**, qRT–PCR analysis of *PIEZO1* (**l**), *PIEZO2* (**m**), *PIK3C3* (**n**) and *AKT1* (**o**) mRNA expression in BMSCs treated with different hydrogels. Data are all presented as mean ± SD. Statistical significance was determined by one–way ANOVA. *p < 0.05, **p < 0.01, ***p < 0.001; ns, not significant. #p<0.05, ###p<0.001; ns, not significant versus control group in **m**-**o**).

We further investigated whether this piezoelectric nanofiber network could provide mechanical stimulation to BMSCs. SEM imaging revealed robust BMSC adhesion to the SKPG-4 nanofiber surface, indicating favorable cell-material interactions (**Fig. 4a**). Quantitative analysis of mechanosensitive ion channels showed that *PIEZO1* levels remained unchanged across groups, whereas *PIEZO2* was significantly upregulated in the SKPG-4 group compared with the control or SKPG-1 group at both the mRNA and protein levels (**Fig. 4j-m**). Concurrently, phosphatidylinositol 3-kinase catalytic subunit type 3 (*PIK3C3*) and AKT serine/threonine kinase 1 (*AKT1*) expression in the SKPG-4 group was significantly downregulated (**Fig. 4n, o**). Given that PIEZO2 activation typically triggers Ca^2+^ influx^38^ and downstream signaling cascades, the observed reduction in PI3K/AKT may reflect engagement of negative feedback regulatory mechanisms, such as PTEN-mediated modulation^39, 40^, in response to sustained mechanotransduction. These findings collectively indicate that the superior mechano-piezoelectric properties of SKPG-4 nanofibers actively transduce physical cues to BMSCs, upregulating PIEZO2 and modulating the PI3K/AKT pathway.

In summary, among the four enzyme–responsive hydrogels, the trifluoromethyl–modified SKPG–4 exhibited outstanding mechano–piezoelectric performance, with a d_33_ of ∼405 pm/V, approximately 15–fold higher than non–fluorinated SKPG–1. This enhanced piezoelectricity enabled SKPG–4 nanofibers to transduce mechanical cues to BMSCs, specifically upregulating PIEZO2 while downregulating PI3K/AKT, with PIEZO1 unchanged. This mechano–piezoelectric effect may underlie the significantly enhanced BMSCs recruitment and chondrogenic differentiation previously observed with fluorinated peptide hydrogels. Thus, SKPG–4 functions as a mechano–piezoelectric biomaterial that couples enzyme–triggered gelation with mechanomodulation of stem cell signaling.

### Simulations Link Fluorination–Enhanced Assembly to SKPG–4 Piezoelectricity

To elucidate the structural basis of fluorination–modulated self–assembly and its contribution to mechano–piezoelectric performance, we performed AA–MD simulations on CMPs (before cleavage) and SKPGs (after cleavage) using GROMACS with the CHARMM36m force field. For CMPs before cleavage, the decamers maintained relatively ordered structures throughout the 50 ns simulation, with RMSD values around 1 nm and minimal fluctuations (**Supplementary Fig. 25a, b**). RMSF analysis further revealed good overall stability for CMP–1/3/4, whereas CMP–2 exhibited substantial local fluctuations, with some residues exceeding 2 nm (**Supplementary Fig. 25c**). After enzymatic cleavage, SKPG–1/3/4 adopted ordered assembly patterns (**Supplementary Fig. 25d**). RMSD and RMSF analyses confirmed that SKPG–4 possessed superior stability, with RMSD values of approximately 0.5 nm and negligible fluctuations over 50 ns, whereas SKPG–2 exhibited poor stability, characterized by a continuous increase in RMSD. (**Fig. 5a** and **Supplementary Fig. 25e, f**).

**Fig. 5.**
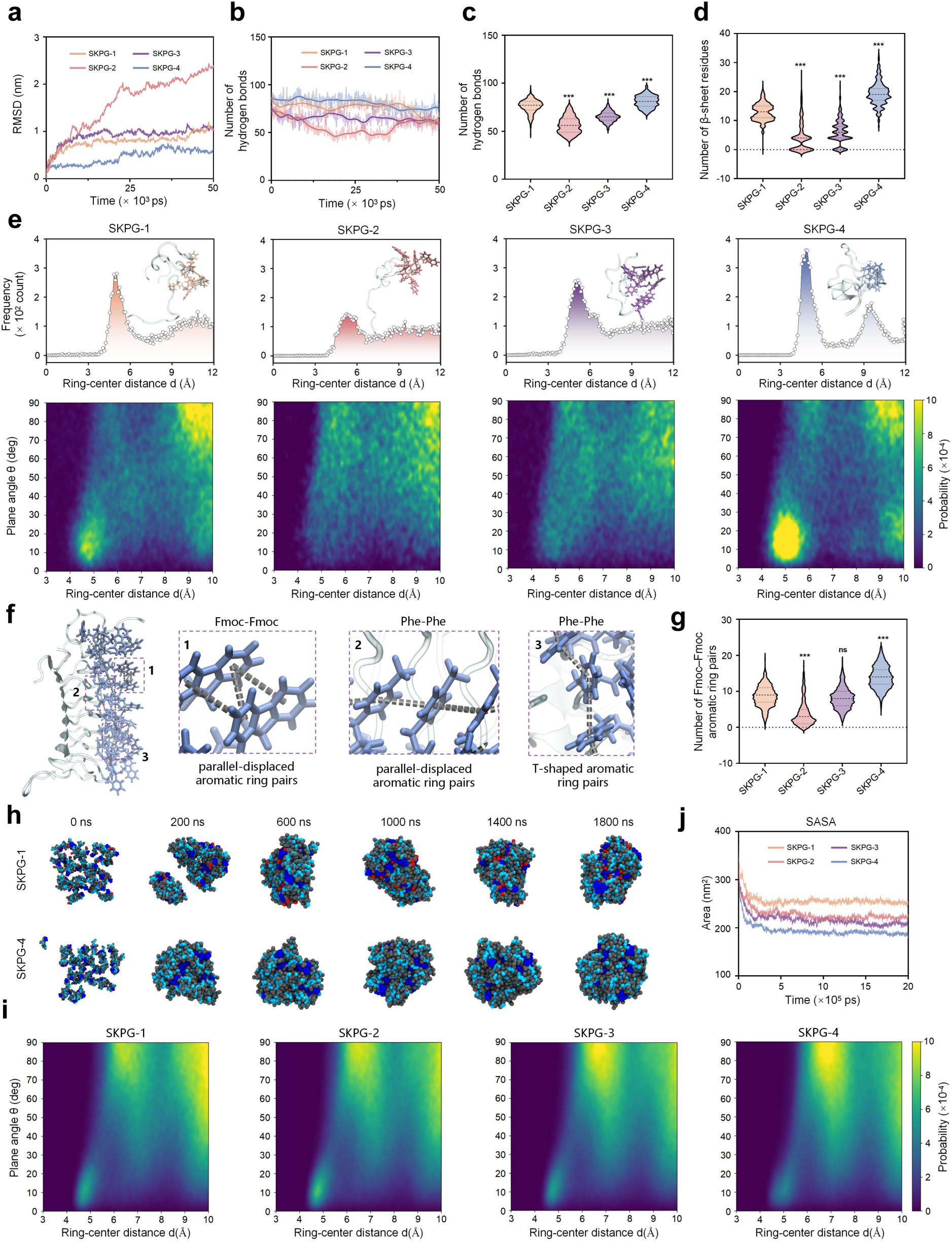
Molecular dynamics simulations reveal fluorination–enhanced assembly of SKPG–4 peptides. **a**, RMSD trajectories of SKPG–1/2/3/4 peptides over 50 ns of AA–MD simulations. **b**, Changes of hydrogen bond numbers within SKPG–1/2/3/4 assemblies over 50 ns. **c**, Statistical comparison of hydrogen bond counts for SKPG–1/2/3/4 peptides over 50 ns. **d**, Quantitative analysis of β-Sheet secondary structures content in SKPG-1/2/3/4 peptides over 50 ns. **e**, Probability density function (PDF) of ring–center distances and free–energy landscapes as a function of centroid distance and dihedral angle between aromatic ring pairs for SKPG–1/2/3/4 oligomers from AA–MD simulations. **f**, Representative configurations showing the spatial arrangements of aromatic rings between Fmoc-Fmoc and Phe-Phe pairs within the assembled SKPG–4 peptide. **g**, Statistical comparison of Fmoc-Fmoc aromatic ring pair interactions from AA–MD simulations. **h**, Representative CG–MD snapshots showing the self–assembly process of SKPG–1 and SKPG–4 peptides into spherical aggregates in aqueous solution. **i**, Free–energy landscapes as a function of ring–center distance and dihedral angle between aromatic ring pairs for SKPG–1/2/3/4 oligomers from CG–MD simulations. **j**, Solvent–accessible surface area (SASA) of SKPG–1/2/3/4 peptide assemblies over 2000 ns of CG–MD simulations. Data are all presented as mean ± SD. Statistical significance was determined by one–way ANOVA. *p < 0.05, **p < 0.01, ***p < 0.001; ns, not significant.

Next, we examined the hydrogen–bonding networks. Hydrogen–bond analysis showed little variation among CMP variants (**Supplementary Fig. 26a**), whereas SKPG–4 formed significantly more hydrogen bonds than the other groups (**Fig. 5b, c** and **Supplementary Fig. 26b**), suggesting a more stable hydrogen–bond network. Finally, secondary structure analysis revealed little variation among CMP variants (**Supplementary Fig. 27a**), but further showed that SKPG–4 contained markedly more β–sheet residues (**Fig. 5d** and **Supplementary Fig. 27b**), in line with the CD and ThT results.

To gain deeper insight into the intermolecular forces governing assembly, we constructed free–energy landscapes as a function of centroid distance and dihedral angle between aromatic ring pairs, categorized into Fmoc-Fmoc, Fmoc-Phe, Phe-Phe, and total contributions (**Fig. 5e** and **Supplementary Fig. 28a-c**). For SKPG–4, the heatmap revealed a higher probability of aromatic ring pairs at a centroid distance of ∼5 Å and dihedral angles of 0-30°, predominantly due to Fmoc-Fmoc and Phe-Phe pairs. This preferential spatial arrangement of aromatic rings-characterized by close proximity and near–parallel orientation-facilitates extensive inter–ring interactions. Notably, the aromatic rings of SKPG–4 exhibit a strong tendency to adopt a parallel, slightly offset orientation between rings (**Fig. 5f** and **Supplementary Fig. 28a-c**). Quantitative analysis of the number of aromatic ring pairs of Fmoc-Fmoc over 50 ns showed that SKPG–4 exhibited higher counts than SKPG–1 (**Fig. 5g**). In contrast, the distribution of inter–ring orientations in SKPG–2 was more heterogeneous, with a broader range of centroid distances and dihedral angles, suggesting less favorable interaction geometries among its aromatic residues.

To explore larger–scale assembly, we performed microsecond–scale CG-MD simulations in aqueous environments. SKPG–4 rapidly organized into a stable, ordered structure within 2000 ns (**Fig. 5h** and **Supplementary Fig. 29a-d**), whereas other peptides formed spherical or cylindrical clusters. Free–energy surfaces for different SKPG systems as a function of ring–center distance and angle revealed similar patterns around 6-7 Å and 70-90° across all four peptides, with distinct interaction geometries observed between Fmoc-Phe and Phe-Phe pairs (**Fig. 5i** and **Supplementary Fig. 30a-f**). Normalized solvent–accessible surface area (SASA) analysis showed stabilization around 250 ns, with the order SKPG–1 > SKPG–2 > SKPG–3 > SKPG–4, indicating that SKPG–4 forms the tightest assembly with the smallest exposed surface area (**Fig. 5j**). Consistent with the above findings, statistical analysis of aromatic ring pair interactions over 2000 ns CG-MD simulations revealed that SKPG–4 possessed more total and Fmoc–Fmoc aromatic contacts than SKPG–1, further supporting its superior assembly propensity (**Supplementary Fig. 31**).

Collectively, our simulations demonstrate that the trifluoromethyl modification in SKPG–4 enhances assembly stability through three synergistic mechanisms: a stronger hydrogen–bond network, higher β–sheet content, and more favorable aromatic ring interactions. The latter is characterized by a high frequency of close–range, near–parallel aromatic ring pairs with narrow dihedral angle distributions, which collectively promote compact and ordered packing. These features drive rapid formation of tightly packed structures with reduced solvent exposure. Critically, this fluorination–enhanced hierarchical assembly directly contributes to the superior mechanical and piezoelectric properties observed experimentally (**Fig. 4**), establishing a molecular–level foundation for the mechano–piezoelectric functionality of SKPG–4. In contrast, other fluorination patterns yield less stable aggregates with inferior assembly performance. These findings provide a mechanistic blueprint for rational biomaterial design, demonstrating that precise fluorination can tune intermolecular interactions to control assembly outcomes and piezoelectric activity.

### G4 promotes macrophage M2 polarization, alleviates oxidative stress and protects chondrocytes

Macrophages dominate the inflammatory microenvironment of osteoarthritis (OA), with M1 macrophages playing a pivotal role in disease progression. Given the superior sustained–release performance of G4 (SKPG–4@4–OI, with G1/2/3 representing SKPG–1/2/3@4–OI, respectively), we investigated whether it could regulate macrophage polarization toward the M2 phenotype and modulate inflammatory cytokine release (**Fig. 6a**). We first assessed the effects of different formulations on macrophage polarization and intracellular ROS levels using flow cytometry and confocal microscopy (**Fig. 6b-i**). Flow cytometry and confocal microscopy revealed that LPS+IFN–γ–induced macrophages (control) exhibited high CD86 and low CD206 expression. Free 4–OI treatment reduced CD86 levels, whereas G1/2/3/4 further increased CD206 expression, with G4 showing the most pronounced effect, consistent with its sustained–release profile (**Fig. 6b-e, h** and **Supplementary Fig. 32a**). We next evaluated ROS scavenging using the 2’,7’-Dichlorodihydrofluorescein diacetate (DCFH–DA) assay. Untreated M1 macrophages exhibited strong green fluorescence, indicating elevated ROS levels. G4 reduced intracellular ROS by approximately 10–fold compared with controls, whereas free 4–OI and G3 achieved only 3–fold and 5–fold reductions, respectively. G1 showed negligible effect, and G2 paradoxically increased oxidative stress, likely due to poor release performance. These findings were corroborated by CLSM imaging (**Fig. 6f, g, i**).

**Fig. 6.**
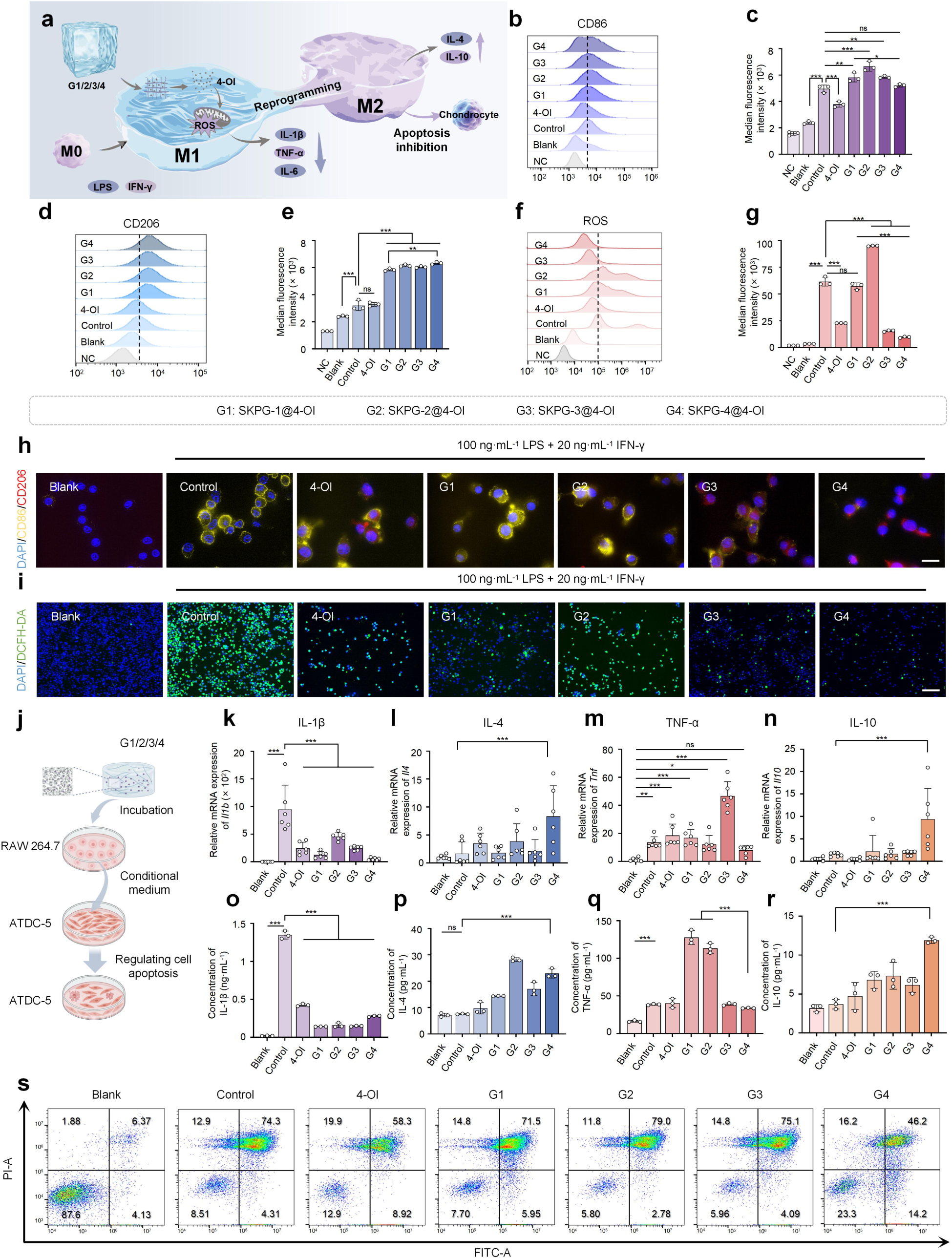
G4 promotes macrophage M2 polarization, alleviates oxidative stress and protects chondrocytes. **a**, Schematic illustration of the mechanisms by which G1/2/3/4 hydrogels regulate macrophage polarization and protect chondrocytes from apoptosis. Created with BioRender. **b**-**e**, Flow cytometry histograms of CD86 (**b**) and CD206 (**d**) expression in macrophages treated with different formulations, with corresponding quantification in (**c**) and (**e**) (n = 3). Flow cytometry histograms (**f**) and quantification (**g**) of intracellular ROS levels in macrophages treated with different formulations (n = 3). NC means negative control, no treatment with LPS+IFN-γ or antibody or probes. **h**, CLSM images of CD86 (green) and CD206 (red) immunostaining in macrophages treated with different formulations. Nuclei were stained with DAPI (blue). Scale bar: 20 μm. **i**, CLSM images of intracellular ROS levels in macrophages treated with different formulations detected by DCFH–DA staining. Scale bar: 20 μm. **j**, Schematic illustration of the experimental procedure for culturing ATDC-5 chondrocytes with CM collected from drug–treated macrophages. Created with BioRender. **k**-**n**, qRT-PCR analysis of *Il1b*, *Il4*, *Tnf* and *Il10* mRNA expression in macrophages treated with different formulations (n = 3). **o**-**r,** ELISA quantification of IL-1β, IL-4, TNF-α and IL-10 secretion from macrophages treated with different formulations (n = 3). **s**, Flow cytometry analysis of apoptosis in ATDC-5 chondrocytes after culture with CM from different treatment groups, assessed by Annexin V–FITC/PI staining, n = 3. Data are all presented as mean ± SD. Statistical significance was determined by one–way ANOVA. *p < 0.05, **p < 0.01, ***p < 0.001; ns, not significant.

Macrophages are also a major source of inflammatory cytokines, including IL–1β, IL–6 and TNF–α, which disrupt cartilage homeostasis by inducing matrix metalloproteinases and chondrocyte apoptosis. qRT-PCR and ELISA analyses of conditioned media (CM) from polarized RAW264.7 macrophages revealed that G4 significantly reduced pro–inflammatory cytokines (IL–1β, IL–6, TNF–α) while elevating anti–inflammatory cytokines (IL–4, IL–10), with effects superior to free 4–OI (**Fig. 6j-r** and **Supplementary Fig. 32b**). To assess the functional impact on chondrocytes, ATDC-5 cells were exposed to CM from treated macrophages. CCK–8 assay revealed that 40% CM from untreated M1 macrophages reduced ATDC-5 viability to 54.59%. In contrast, CM from G1/2/3/4–treated macrophages increased viability to 54.98%, 86.84%, 88.44% and 93.38%, respectively (**Supplementary Fig. 32c**). Annexin V–FITC/PI staining further demonstrated that G4 reduced the late apoptosis rate of ATDC-5 cells from 74.3% (untreated CM) to 46.2% (**Fig. 6s** and **Supplementary Fig. 32d**).

Collectively, our findings establish that G4, by enabling sustained 4–OI release, drives macrophage M2 polarization, relieves oxidative stress, curtails pro–inflammatory cytokine production and shields chondrocytes from apoptosis. These combined immunomodulatory and cytoprotective effects position G4 as a promising therapeutic strategy for OA.

### CMP-4/4-OI exerts synergistic therapeutic effects *in vivo*

To systematically evaluate the therapeutic efficacy of fluorinated CMPs/4–OI in osteoarthritis (OA), we utilized a destabilization of the medial meniscus (DMM) rat model. Intra–articular injections of CMPs (20 mg·mL^-1^) with 4–OI (1 mg·mL^-1^, 0.5% DMSO) were administered at weeks 6 and 8 post–surgery (**Fig. 7a, b**). Micro–CT imaging at 8 weeks post–surgery revealed that CMP–4/4–OI most effectively attenuated bone erosion and osteophyte formation among all treatment groups (Fig. 7c, d). Synovial inflammation was also markedly suppressed by CMP–4/4–OI, as evidenced by reduced inflammatory cell infiltration in Hematoxylin and Eosin (H&E)-stained synovial sections (**Fig. 7e**).

**Fig. 7.**
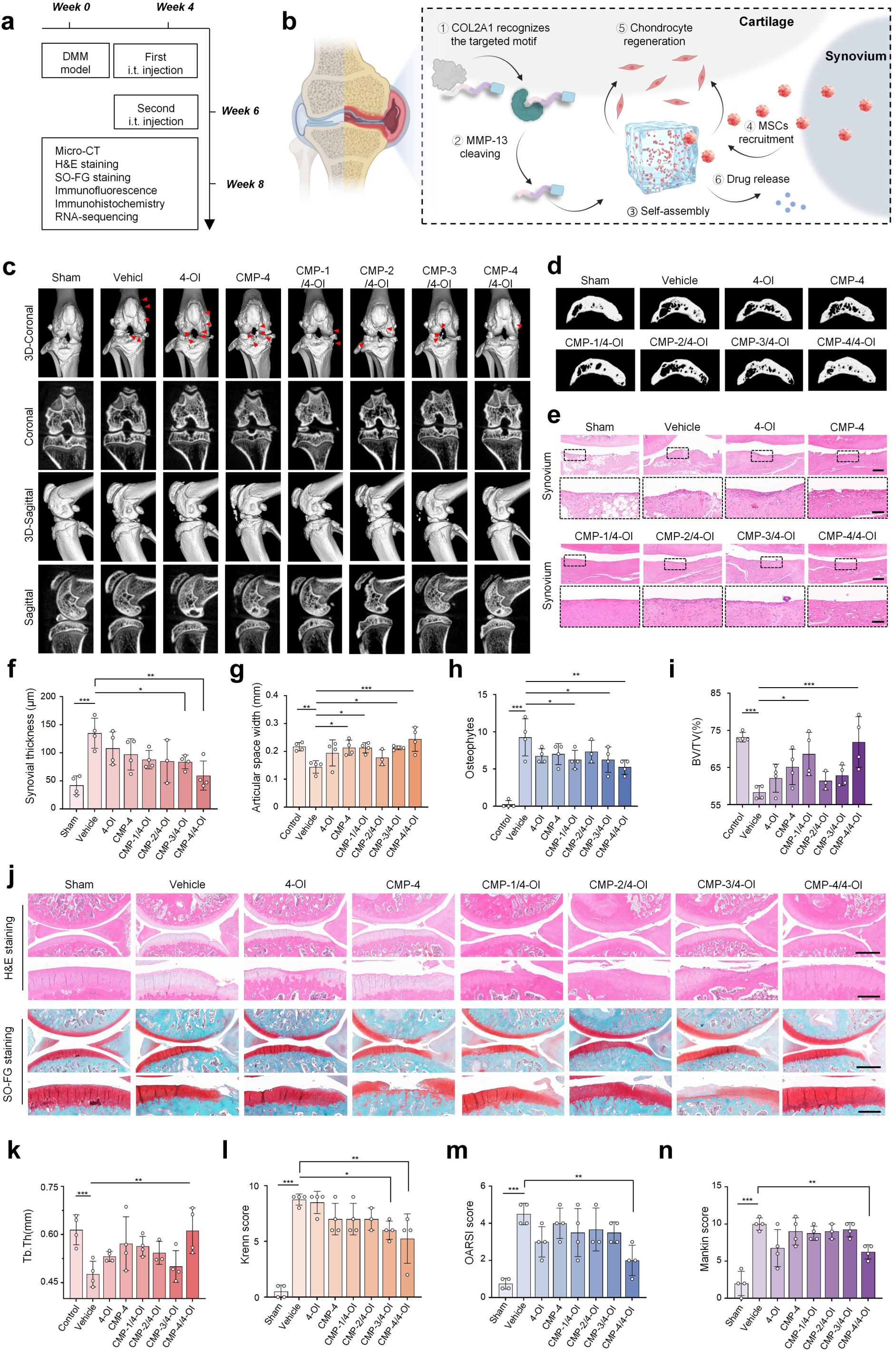
Fluorination–engineered CMP–4/4–OI hydrogel alleviates osteoarthritis in rat DMM models. **a**, Schematic diagram of the time course of *in vivo* therapeutic experiments in DMM rat models. **b**, Schematic diagram of the therapeutic mechanism of CMPs/4–OI in the rat knee joint. **c**, Three–dimensional and planar reconstructions of rat knee joints showing osteophyte formation (arrows) in Sham, Vehicle, 4–OI, CMP–4, CMP–1/4–OI, CMP–2/4–OI, CMP–3/4–OI and CMP–4/4–OI groups at 8 weeks post–surgery. **d**, Three-dimensional reconstructions of the medial tibial subchondral bone at 8 weeks after DMM surgery. **e**, H&E-stained synovial tissue of joint sections. Scale bars: 1 mm (overview) and 0.4 mm (insets). **f**, Synovial thickness quantification. **g**, Articular space width quantification. **h**, Osteophyte number quantification. **i**, BV/TV quantification. **j**, H&E and SO–FG staining of knee joint sections. Scale bars: 1 mm (overview) and 0.4 mm (insets). **k**, Tb.Th quantification. **l**, Krenn score. **m**, OARSI score. **n**, Mankin score. Data are mean ± SD (n = 4 per group, except n = 3 for CMP–2/4–OI). Data are all presented as mean ± SD. One–way ANOVA. *p < 0.05, **p < 0.01, ***p < 0.001; ns, not significant.

Quantitative analyses of synovial inflammation, bone microstructure and cartilage histology collectively confirmed the superior therapeutic efficacy of CMP-4/4-OI (**Fig. 7f-n**). Synovial thickness was significantly decreased from 134.9 μm (Vehicle) to 59.5 μm, representing a reduction of approximately 56% (**Fig. 7f**). Bone microstructural parameters were also markedly improved. Specifically, joint space width increased from 0.14 to 0.24 mm, osteophyte numbers decreased from 9.3 to 5.3, bone volume fraction (BV/TV) increased from 58.3% to 71.9%, and trabecular thickness (Tb.Th) increased from 0.48 to 0.61 mm (**Fig. 7g-i, k**). These quantitative improvements indicate that CMP-4/4-OI effectively restored subchondral bone architecture and mitigated aberrant osteophyte formation. Notably, the other fluorinated CMP variants (CMP-1/4-OI, CMP-2/4-OI and CMP-3/4-OI) exhibited less pronounced effects, underscoring the superior osteoprotective efficacy of CMP-4/4-OI in improving OA-related bone microstructural parameters.

Histological evaluation by H&E and Safranin O-Fast Green (SO-FG) staining further revealed that CMP-4/4-OI treatment significantly reduced cartilage erosion, restored proteoglycan distribution and preserved collagen fibers compared with the Vehicle group (**Fig. 7j**). In good agreement, the Krenn, OARSI and Mankin scores were all markedly decreased in the CMP-4/4-OI group (Krenn: from 8.8 to 5.3; OARSI: from 4.5 to 2.0; Mankin: from 10.0 to 6.3) (**Fig. 7l-n**), confirming its capacity to alleviate cartilage matrix degradation and preserve overall joint integrity. Together, these *in vivo* data demonstrate that CMP-4/4-OI not only suppresses synovial inflammation and improves bone microarchitecture, but also effectively protects cartilage from OA-induced degeneration.

IHC and immunofluorescence (IF) staining analysis further corroborated the *in vivo* immunomodulatory and regenerative effects of CMP-4/4-OI (**Fig. 8a-f**). Specifically, IHC analysis revealed that CMP–4/4–OI significantly upregulated COL2A1 expression and downregulated IL–1β levels in the knee joints compared with the Vehicle group (**Fig. 8a, c, d**). Immunofluorescence staining for CD206 revealed a marked increase in M2–type macrophages on the synovial surface in the CMP–4/4–OI group (**Fig. 8b, e**), consistent with our *in vitro* findings of macrophage M2 polarization. Additionally, CD73/CD90 co–staining confirmed the presence of dual–positive cells (MSCs) on the cartilage and synovial surfaces, with the CMP–3/4–OI and CMP–4/4–OI groups exhibiting significantly higher cell numbers, indicating enhanced stem cell recruitment (**Fig. 8f**).

**Fig. 8.**
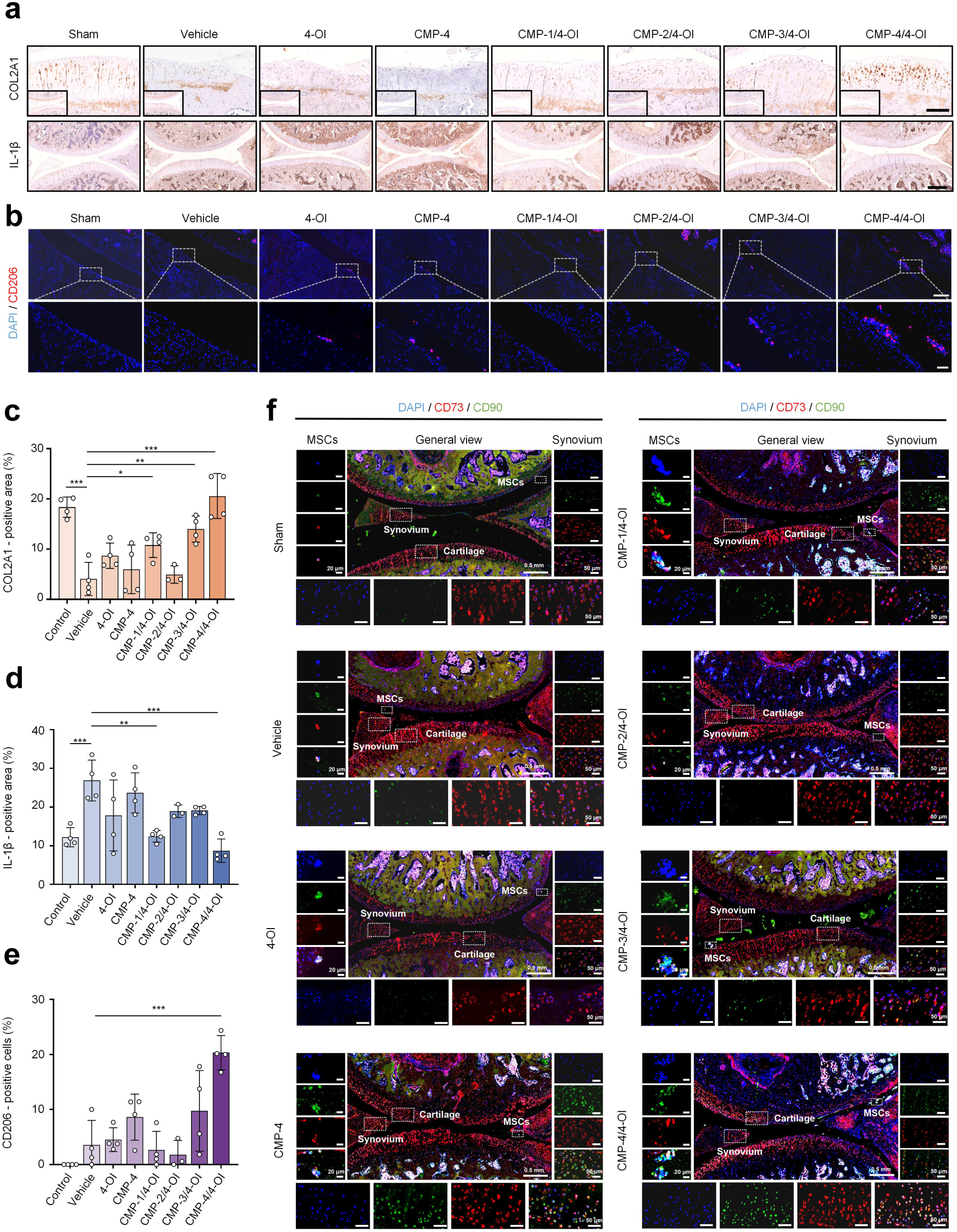
CMP–4/4–OI promotes cartilage repair and immunomodulation *in vivo*. **a**, Immunohistochemical staining of COL2A1 and IL–1β in rat knee joint sections. Scale bars: 0.2 mm (COL2A1) and 0.5 mm (IL–1β). **b**, Immunofluorescence staining of CD206 (green) in knee joint sections. Nuclei were stained with DAPI (blue). Scale bars: 0.2 mm (overview) and 50 μm (insets). Quantitative analysis of COL2A1 (**c**) and IL-1β (**d**) expression in knee joint cartilage after treatment with the indicated formulations (n = 4 per group, except n = 3 for the CMP–2/4–OI group). **e**, Quantitative analysis of CD206–positive area in knee joint sections after treatment with the indicated formulations (n = 4 per group, except n = 3 for the CMP–2/4–OI group). **f**, Immunofluorescence co–staining of CD73 (green) and CD90 (red) for detecting endogenous MSCs recruitment to the knee joint. Scale bars: 0.5 mm (overview), 50 μm (synovium) and 20 μm (cartilage). Data are presented as mean ± SD. Statistical significance was determined by one–way ANOVA. *p < 0.05, **p < 0.01, ***p < 0.001; ns, not significant.

To explore the versatility of this delivery platform, we substituted 4–OI with dexamethasone (Dex) in an ACLT–induced OA rat model. CMP–4/Dex treatment similarly attenuated osteophyte formation and preserved joint integrity, as shown by histological staining and transcriptomic analysis (**Supplementary Fig. 33-35**).

Collectively, these findings establish that the CMP–4/4–OI hydrogel integrates mechano–piezoelectric cues with biochemical therapy to address the complex pathophysiology of OA. The fluorination–enhanced nanofiber assembly not only improves bone microarchitecture and reduces osteophyte formation, but also provides a piezoelectric scaffold that actively transduces physical signals to endogenous stem cells. Concurrently, the sustained release of 4–OI reprograms macrophages toward the M2 phenotype, suppresses IL–1β and promotes a pro–regenerative immune microenvironment, while upregulating COL2A1 expression to restore cartilage matrix integrity. This dual mechano–biochemical synergy, combining piezoelectric stimulation with immunomodulatory drug delivery, enables simultaneous cartilage protection, inflammation resolution and endogenous stem cell recruitment, positioning CMP–4/4–OI as a clinically promising strategy for OA therapy.

### Transcriptomic analysis reveals dual mechanisms of CMP 4/4 OI: ECM anabolism and metabolic-immune reprogramming

To elucidate the molecular mechanisms underlying the superior efficacy of CMP 4/4-OI, we performed mRNA sequencing on rat knee joints after treatment. We focused on two key comparisons: CMP-4/4-OI versus Vehicle (overall treatment effect) and CMP-4/4-OI versus CMP-1/4-OI (trifluoromethyl modification advantage). A total heatmap of differentially expressed genes (DEGs) indicated that CMP-4/4-OI exhibited the most pronounced transcriptomic alterations among all treatment groups (**Supplementary Fig. 36a**).

Kyoto Encyclopedia of Genes and Genomes (KEGG) pathway analysis of DEGs from both comparisons revealed common enrichment in oxidative phosphorylation, gluconeogenesis, cell cycle, Fc gamma R–mediated phagocytosis, cellular senescence and apoptosis (**Fig. 9a, b**). These pathways are closely linked to the metabolic reprogramming and immunomodulatory functions we observed in previous experiments (**Fig. 6**). Notably, the CMP–4/4–OI versus CMP–1/4–OI comparison showed specific enrichment in ECM organization, cell adhesion and tissue remodeling, which are processes directly relevant to cartilage repair. Gene Set Enrichment Analysis (GSEA) further confirmed that CMP–4/4–OI treatment robustly enriched cartilage development gene sets while negatively enriching inflammatory response and senescence sets, with effect magnitudes substantially exceeding those of CMP–1/4–OI (**Fig. 9c, d**). Heatmap analysis revealed that CMP–4/4–OI coordinately upregulated genes associated with cell migration and ECM organization (**Fig. 9e, f**), which are processes directly relevant to MSCs recruitment and cartilage matrix repair, while concurrently enhancing oxidative phosphorylation and M2–associated macrophage polarization gene signatures (**Fig. 9g, h**), consistent with our *in vitro* findings on metabolic reprogramming and immunomodulation (**Fig. 6**).

**Fig. 9.**
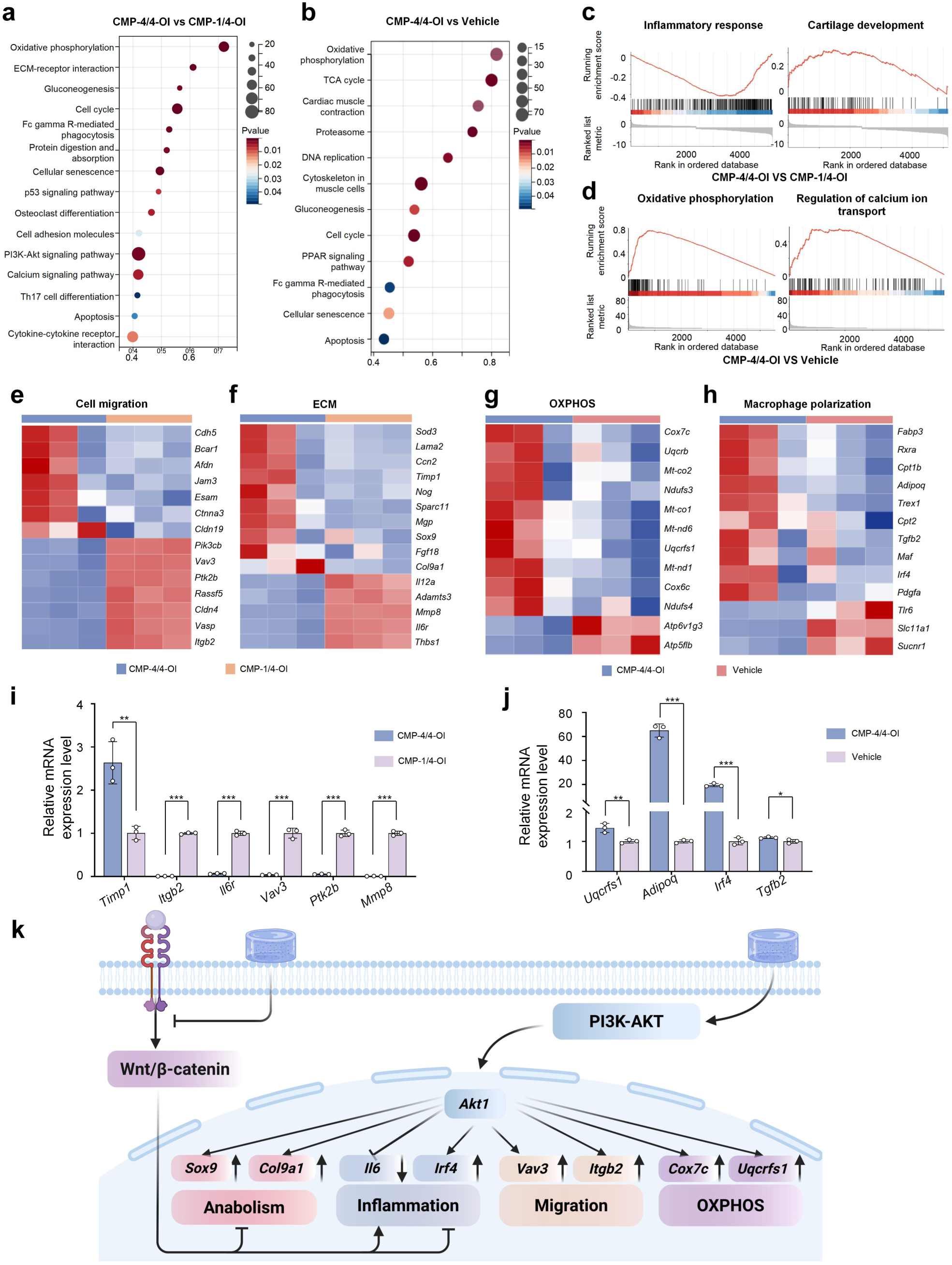
Transcriptomics reveals metabolic and ECM–remodeling pathways activated by CMP–4/4–OI in OA. **a**, KEGG pathway enrichment analysis of DEGs in CMP-4/4-OI versus CMP-1/4-OI groups. **b**, KEGG pathway enrichment analysis of DEGs in CMP-4/4-OI versus Vehicle groups. DEGs were defined with cutoffs of FDR < 0.05 and |log₂ fold change| > 1. GSEA of inflammatory response and cartilage development (**c**) between CMP–4/4–OI and CMP–1/4–OI, and oxidative phosphorylation and regulation of calcium ion transport (**d**) between CMP–4/4–OI and Vehicle groups. **e**, **f**, Heatmaps of genes related to cell migration and ECM organization in CMP–4/4–OI and CMP–1/4–OI groups (n = 3). **g**, **h**, Heatmaps of genes related to oxidative phosphorylation (OXPHOS) and macrophage polarization in CMP–4/4–OI and Vehicle groups (n = 3). **i**, qRT–PCR validation of genes related to cell migration and ECM organization in CMP–4/4–OI versus CMP–1/4–OI groups (n = 3). **j**, qRT–PCR validation of genes related to OXPHOS and macrophage polarization in CMP–4/4–OI versus Vehicle groups (n = 3). **k**, Schematic diagram of the proposed mechanism by which CMP–4/4–OI hydrogel protects chondrocytes. Created with BioRender. Data are presented as mean ± SD. Statistical significance was determined by unpaired t–test. *p < 0.05, **p < 0.01, ***p < 0.001 versus control group.

These transcriptomic findings align with and extend our previous observations: the mechano–piezoelectric properties of SKPG–4 nanofibers (**Fig. 4**) and the sustained 4–OI release driving macrophage M2 polarization (**Fig. 6**) converge at the molecular level to activate the PI3K/AKT pathway while suppressing Wnt/β–catenin signaling. This dual regulation was further validated by qRT-PCR analysis of key DEGs (**Fig. 9i, j**). The activation of PI3K/AKT promotes chondrocyte survival and anabolic gene expression, whereas suppression of Wnt/β–catenin reduces catabolic and inflammatory signals, collectively relieving IL–1β–induced chondrocyte damage (**Fig. 9k**). Thus, CMP–4/4–OI orchestrates a coordinated transcriptional program that not only mitigates pathological processes like inflammation and senescence but also actively promotes ECM anabolism and cartilage regeneration, distinguishing it from its non–fluorinated counterpart, which lacks the synergistic benefits of trifluoromethyl modification and sustained 4–OI release.

## Conclusion

In this study, we have developed a mechano–piezoelectric peptide hydrogel system that integrates fluorination–enhanced piezoelectricity with targeted biochemical therapy to address the complex pathophysiology of OA. Our findings establish that the trifluoromethyl–modified SKPG–4 peptide, through hierarchical self–assembly into ordered nanofibers, exhibits a substantially enhanced piezoelectric coefficient (d_33_ ≈ 405 pm/V, ∼15–fold higher than its non–fluorinated counterpart). This piezoelectric output, together with the sustained release of the immunomodulatory agent 4–OI, creates a synergistic mechano–biochemical platform that simultaneously promotes MSCs recruitment and chondrogenic differentiation, reprograms macrophages toward the M2 phenotype, and suppresses the inflammatory microenvironment.

Mechanistically, the system operates through coordinated molecular interaction: mechano-piezoelectric cues activate PIEZO2-mediated mechanotransduction and engage the PI3K/AKT pathway, while 4–OI drives metabolic reprogramming and M2 polarization; these signals converge to suppress Wnt/β–catenin signaling, possibly upregulate anabolic ECM genes (*Col9a1*, *Sox9*), and downregulate catabolic and inflammatory mediators (*Mmp8*, *Il1b*). In a rat DMM model, CMP-4/4-OI significantly improved bone microarchitecture (increased BV/TV and joint space width, reduced osteophytes), cartilage integrity (OARSI and Mankin scores), and synovial inflammation, while promoting endogenous MSCs homing and M2–type macrophage accumulation at the joint surface. The platform also demonstrated versatility in delivering alternative therapeutics, such as dexamethasone, suggesting broad applicability.

Despite these promising results, several limitations warrant acknowledgment. The translational relevance to human OA, which develops over decades with more complex mechanical and inflammatory milieu, remains to be established. The long-term safety and degradation profile of fluorinated peptides require systematic evaluation, and optimal dosing regimens need further optimization. Beyond OA, this platform holds promise for other musculoskeletal disorders characterized by mechanical dysfunction and chronic inflammation, including tendinopathy, meniscus injury, and intervertebral disc degeneration. The modular peptide design also allows facile substitution of targeting ligands, enzyme-responsive sequences, and therapeutic payloads for broader disease applications. With continued optimization, this mechano-piezoelectric hydrogel could offer a minimally invasive, disease-modifying therapeutic option for OA and related conditions.

## Methods

### Ethics approval

All experimental procedures involving animals were reviewed and approved by the Institutional Animal Ethics Committee of Animal Center of Institute of Materia Medica, CAMS & PUMC (ID: IMM-S-25-0312, IMM-S-25-0240) and were carried out in accordance with the Care and Use of Laboratory Animals (Ministry of Health, China). This study was approved by the Ethics Committee Review Board of The Ethics Committee of The Second Affiliated Hospital of Soochow University (Approval No. JD-LK2023123-I01). All procedures performed in studies involving human participants were in accordance with the ethical standards of the institutional and national research committee and with the 1964 Helsinki Declaration and its later amendments or comparable ethical standards. Informed consent was obtained from all individual participants included in the study.

### Peptide synthesis

Peptide SKP-1, SKP-2, SKP-3, SKP-4, TPE-SKP-3, CMP-1, CMP-2, CMP-3, CMP-4 were prepared by Shanghai Yaxian Chemical Co.,Ltd (Shanghai, China). SKPG-1, SKPG-2, SKPG-3, SKPG-4, Cy5-SKP-1, Cy5-SKP-4, Cy5-CMP-1, Cy5-CMP-4 were prepared by Qingdao KL Biotechnology Co., LTD (Qingdao, China).

### Preparation of self-assembled hydrogels and hydrogels@4-OI

The lyophilized peptide (SKP-1/2/3/4, TPE-SKP-3, SKPG-1/2/3/4) powder was dissolved in ddH_2_O or PBS to achieve a final concentration of 10 mg·mL^-1^. The solution was allowed to stand at 4℃ or room temperature for approximately 15 minutes, forming an almost transparent self-assembling peptide hydrogel. Subsequently, a solution of 4-OI (Cat No. 3133-16-2, Shanghai Xin Heng Yan Technology) or Dex dissolved in 10% DMSO was added to the hydrogel achieving a final concentration of 0.1 mg/mL for 4-OI/Dex.

### SEM and EDS

The peptide hydrogel sample was diluted to 0.1% (w/v) with deionized water. A 5 μL droplet of the diluted sample was placed onto a carbon–coated copper grid (300 mesh) using fine tweezers. After 1 min of adsorption, the excess liquid was gently blotted away with filter paper. The grid was then immediately plunged into liquid nitrogen for 30 s and transferred to a freeze dryer for overnight lyophilization. The dried samples were subsequently observed under an *in situ* high–resolution field–emission scanning electron microscope (FE–SEM, Apreo, Thermo Fisher Scientific) operating at an accelerating voltage of 5 kV. EDS analysis was performed on the same instrument at the China Academy of Information and Communications Technology (CAICT) to confirm the presence of carbon (C), nitrogen (N), oxygen (O) and fluorine (F) elements within the hydrogels.

### TEM

Hydrogel specimens were diluted to 0.1 mg·mL^-1^ in deionized water. A 5 μL droplet was deposited onto glow–discharged, carbon–coated copper grids (300 mesh). After 1 min of adsorption, the grid was blotted with filter paper to remove excess solvent. Negative staining was performed with 1% (w/v) phosphotungstic acid (pH 7.0) for 45 s, followed by rinsing with deionized water and air–drying at room temperature. Bright–field TEM imaging was carried out under low–dose conditions at an accelerating voltage of 200 kV to minimize beam–induced deformation. TEM characterization was performed at Hangzhou Yanqu Information Technology Co., Ltd.

### hBMSC migration assay

The SKP-1/2/3/4 hydrogels were placed in the lower compartment of 24-well Transwell plates. hBMSCs (CP-H166Y, Procell) were seeded into the upper inserts at a density of 1 × 10^5^ cells per well in DMEM supplemented with 10% FBS. After incubation at 37 ℃ for 18 h, the non–migrated cells on the upper surface of the inserts were gently removed with a cotton swab. The migrated cells on the lower surface were fixed with ice–cold methanol for 30 min, washed twice with PBS, and stained with 2% (w/v) crystal violet for 15 min. After washing twice with PBS, the stained cells were imaged under a light microscope and quantified.

### hBMSCs differentiation assay

To generate cell spheroids, hBMSCs were first detached using 0.25% trypsin containing EDTA, collected by centrifugation, and reconstituted in culture medium to achieve a final concentration of 1 × 10^5^ cells per 100 μL. The cell suspension was then incubated in chondrogenic differentiation medium (Human Bone Marrow Mesenchymal Stem Cell Chondrogenic Differentiation Medium) for 24 h at 37 ℃ under a 5% CO₂ atmosphere. After incubation, the formed spheroids were embedded into SKP–1/2/3/4 hydrogels. Cultures were continued for 14 days, during which the medium was replaced every other day.

### Histological staining

All histological staining procedures, including hematoxylin and eosin, safranin O-fast green, immunohistochemistry and immunofluorescence were performed in collaboration with Servicebio Co., Ltd. (Wuhan, China).

### FRET assay

Collagenase-3 (MMP-13) was dissolved in 25 mL Tris-HCl buffer (pH 7.5) to prepare 5 CDU/mL MMP-13 stock solution. FRET-1 peptide (1 mg) was dissolved in 200 μL of DMSO. Take 20 μL of the above peptides and add them to 300 μL of Tris-HCl buffer and collagenase Tris-HCl buffer, respectively, and incubate for 1, 3, 12, and 24 hours. Use a fluorescence ELISA reader to quantify fluorescence intensity, and deduct solvent background fluorescence before use.

### CD

For CD spectroscopy, SKPG-1/2/3/4 peptides were dissolved in ddH_2_O at a concentration of 0.2 mg·mL^-1^. CD spectra were recorded on a JASCO J-815 spectropolarimeter (JASCO, Tokyo, Japan) using a 0.1 cm path length quartz cuvette at 25 ℃. Spectra were collected over a wavelength range of 190-260 nm with a scanning speed of 50 nm min^-1^, a bandwidth of 1.0 nm, and a response time of 1 s. Three accumulations were averaged for each sample, and the spectrum of ddH_2_O was subtracted as blank. The results were expressed as mean residue ellipticity (deg·cm^2^·dmol^-1^).

### 1, 8-ANS binding assay

For the 1, 8–ANS binding assay, peptide stock solutions (1 mM in PBS) were diluted to 250 μM in PBS and incubated with 10 μM 1,8–ANS for 15 min at room temperature in the dark. Fluorescence emission spectra were recorded on a fluorescence microplate reader over a wavelength range of 430-550 nm with excitation at 370 nm. The blue shift of the maximum emission wavelength and the increase in fluorescence intensity were used to evaluate the self–assembly propensity of the peptides, with greater shifts indicating stronger hydrophobic core exposure and assembly capability. All measurements were performed in triplicate.

### ThT binding assay

For the ThT binding assay, SKPG-1/2/3/4 hydrogels were incubated with 80 μM ThT in PBS (pH 7.4) for 120 min at room temperature in the dark. Fluorescence intensity was measured using a microplate reader with excitation at 445 nm and emission at 485 nm. PBS buffer with ThT was used as blank, and all measurements were performed in triplicate. Higher fluorescence intensity indicated greater β-sheet content within the hydrogel assemblies.

### Drug release of SKPG peptides

For the *in vitro* release study, SKPG-1/2/3/4 peptides (4 mg each) were dissolved in 200 μL PBS (pH 7.4) to form hydrogels, into which 4-OI (0.5 mg in 20% DMSO) was loaded. Each hydrogel was transferred into a pre-swelled dialysis bag (MWCO: 1 kDa) and immersed in 20 mL of release medium (PBS at pH 7.4 or 6.5) at 37 ℃ with orbital shaking at 100 rpm. At predetermined time intervals, 200 μL of dialysate was collected and replaced with an equal volume of fresh medium. The concentration of 4-OI in the collected samples was measured by UPLC-MS, and the cumulative release was calculated as a percentage of the total loaded 4-OI.

### IVIS imaging

For the *in vivo* retention study, Cy5, G1@OC, G2@OC, G3@OC and G4@OC were injected into the knee joints of 6-to 8-week-old C57BL/6 mice purchased from GemPharmatech Co., Ltd. (Beijing, China). At predetermined time intervals (1, 3, 4, 5 and 7 days post–injection), mice were anesthetized and the fluorescence signals in the knee joints were captured using an IVIS imaging system (n = 3 per group). Fluorescence intensity was quantified from the region of interest (ROI) over the knee joint to evaluate the retention kinetics of different hydrogel formulations.

### PFM

PFM measurements were conducted using a Bruker Dimension Icon atomic force microscope system. Conductive probes (SCM-PIT-V2) were employed. Prior to the PFM measurements, the deflection sensitivity of the cantilever was calibrated on a rigid substrate, yielding a value of 97.0 nm/V. The electrical excitation parameters were set as follows: an AC drive voltage (Drive Voltage) with an amplitude of 10 V was applied between the conductive tip and the bottom electrode of the sample. Concurrently, a sweeping DC bias voltage (Excitation Voltage) was superimposed, with the potential linearly scanned from -10 V to +10 V.

### Macrophage polarization assessment

RAW264.7 cells (CL-0190, Procell), 5 × 10^4^ per well in 24-well plates, were stimulated with LPS (100 ng·mL^-1^) and IFN–γ (20 ng·mL^-1^) for 24 h, then treated with G1/2/3/4 for another 24 h. After washing with PBS, nonspecific binding was blocked with CD16/32 in 1% BSA. Cells were stained with fluorescent secondary antibodies and DAPI, then imaged by confocal laser scanning microscopy (CLSM; BZ-X810, Keyence, Japan). Semi–quantitative analysis was performed using ImageJ. For flow cytometry, cells per group were incubated with PE anti–mouse CD86 (1:100; Proteintech, USA) and APC anti–mouse CD206 (3:200; Abclonal, USA). Mean fluorescence intensity was measured on an Agilent NovoCyte–D2060R and analyzed with FlowJo_v10.8.1.

### Intracellular ROS detection

RAW264.7 cells were stimulated by 100 ng·mL^-1^ LPS plus 20 ng·mL^-1^ IFN-γ and then incubated with G1/2/3/4 for 24 hours. Then the media was removed, incubated with 10 μM DCFH-DA (1:1000; S0033S; Beyotime Biotechnology, China) in serum-free culture medium for 30 min at 37℃. Next, the fluorescent images were observed in CLSM and the flow cytometry was used for quantitative detection of ROS mean fluorescence intensity.

### Detection of inflammatory cytokines in CM

RAW264.7 cells were plated on 6-well plates at a density of 1 × 10^6^ cells per well and incubated with LPS plus IFN-γ and various treatment groups. After 24 hours, the supernatants of polarized macrophages were collected. The obtained supernatants were collected as mentioned above. To detect macrophage M1/M2 polarization and inflammatory cytokines in culture medium, the CM using a mouse IL-1β ELISA kit (CSB-E08054m-IS, Cusabio), a mouse IL-4 ELISA kit (CSB-E04634m-IS, Cusabio), a mouse IL-10 Elisa kit (CSB-E04594m-IS, Cusabio) and a mouse TNF-α ELISA kit (CSB-E04741m-IS, Cusabio) according to the manufactures protocol.

### CCK-8

ATDC-5 cells were seeded in 96–well plates at a density of 5 × 10³ cells/well and cultured overnight. After incubated with different CM at 0%, 5%, 10%, 20%, 40% and 80% for 48 h, 10 μL of CCK–8 solution (AQ308-10000T, Aoqing Biotechnology Ltd) was added to each well and incubated at 37 ℃ for 2 h. Absorbance was recorded at 450 nm using a microplate reader. Cell viability was calculated as:

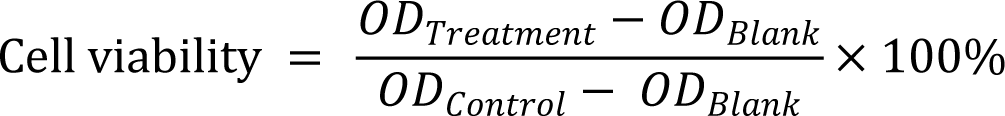

All assays were performed in triplicate, and data are presented as mean ± SD (n = 3)

### Apoptotic cell death mechanism

The apoptotic cell death mechanism was detected by annexin V-FITC apoptosis detection Kit (C1062M, Beyotime Biotechnology). Specifically, ATDC-5 cells (CL-0856, Procell) were seeded on 6-well plates at a density of 1 × 10^6^ cells per well overnight. After that, the media was removed, and the cells were incubated with CM of various treatment group for 48 hours. Then the dead and live cells were collected and further incubated with annexin V/propidium iodide (1:100 dye solution) for 15 min. The cells were washed with phosphate-buffered saline and the cell death was assessed by flow cytometry.

### Destabilization of the DMM-induced OA rat model and treatment

All male SD rats (7 weeks old, from Beijing Vital River Laboratory Animal Technology Co., Ltd) were housed under specific–pathogen–free (SPF) conditions with a 12–h light/dark cycle and free access to food and water. The destabilization of the DMM model was surgically induced as previously described. Briefly, after anesthesia and skin preparation, the right knee joint was exposed via a medial parapatellar approach. With the knee fully flexed, the anterior medial meniscotibial ligament was transected using a microsurgical knife, and complete transection was confirmed by manual displacement of the medial meniscus. The joint cavity was rinsed with sterile saline, and the capsule and skin were closed with 6-0 and 5-0 sutures, respectively. Sham operations underwent the same incision and capsulotomy without ligament transection. Thirty–two 8–week–old male SD rats were randomly divided into eight groups: Sham, Vehicle (DMM model), and DMM treated with 4–OI, CMP–4, CMP–1/4–OI, CMP–2/4–OI, CMP–3/4–OI, or CMP–4/4–OI (n = 4 per group). All treatments were administered via intra–articular injection (50 μL per joint) at weeks 6 and 8 post–surgery. The CMP peptides were injected at 20 mg·mL^-1^ with 4-OI at 1 mg·mL^-1^ in PBS, while the Sham and Vehicle groups received an equal volume of PBS. All animal procedures were performed in accordance with the institutional guidelines for animal care and use.

### Micro-CT

Rat knee joints were collected 8 weeks after DMM surgery. Reconstructed imaging of the rat knee joint was performed using Micro PET/CT (Inviscan imaging systems, France). Imaging was performed at a resolution of 60 μm using X-ray energy of 80kV. After dataset reconstruction and orientation, 3D analysis of osteophyte development, tissue calcification, and subchondral bone remodeling was conducted using Imalytics.

### RNA Sequencing and bioinformatics analysis

RNA sequencing was conducted by Shanghai Majorbio Bio-pharm Technology Co., Ltd. Raw reads were aligned to the Rattus norvegicus genome (Rnor_6.0) using HISAT2 v2.2.1, and gene expression quantification was performed using featureCounts v2.0.1. Differential gene expression analysis was performed using DESeq2 (v1.30.1) in R (v4.1.0). Genes with an adjusted P value of <0.05 and an absolute |log_2_ FC| ≥ 1 were considered differentially expressed. Low-abundance genes (mean counts < 10) were filtered out before analysis. Gene Ontology (GO) and Kyoto Encyclopedia of Genes and Genomes (KEGG) pathway enrichment analyses were carried out using Metascape (v3.5) with the Rattus norvegicusdatabase. P value of <0.05, a minimum count of 3, and an enrichment factor of >1.5 were considered statistically significant. Redundant terms were clustered based on semantic similarity with a cutoff of 0.7. Gene Set Enrichment Analysis (GSEA) was implemented via the clusterProfiler package (v4.0.5). All genes, ranked in descending order by log_2_FC, were evaluated against the MSigDB GO Biological Processes gene set collection. Gene sets with a false discovery rate (FDR) below 0.25 and Pvalue of <0.05 were considered significantly enriched.

### Statistical analysis

Data were expressed as the mean ± SD of biological replicates, unless otherwise indicated. Statistical analysis was performed using GraphPad Prism 10.2.3 software. Detailed statistical methods are described in the figure legends. All statistical tests were two sided. A P value of less than 0.05 was considered statistically significant.

## Data availability

The data that support the findings of this study are available from the corresponding author upon reasonable request. Source data are provided with this paper.

## Acknowledgements

This study was supported by the Nonprofit Central Research Institute Fund of the Chinese Academy of Medical Sciences (No. 2022-RC350-04), the National Natural Science Foundation of China (Nos. 82372002 and 82502399), the CAMS Innovation Fund for Medical Sciences (Nos. 2025-I2M-XHJC-026 and 2025-I2M-XHXX-098), and the Beijing Nova Program Interdisciplinary Cooperation Project to K. H.

## Author contributions

X. Song performed hydrogel characterization, cell experiments, partial animal experiments, data curation, and manuscript writing. Z. Xu conducted cell and animal experiments. S. Zhang conducted some cell experiments and wrote the manuscript. T. Zhang analyzed transcriptome sequencing data and generated plots. C. Liu performed partial animal experiments. H. Huang performed partial computational simulations. Y. Hu conducted rat CT imaging. M. Yang organized and curated the raw data. L. Zhao revised and polished the manuscript.

