## Supporting Information for "Mechano-Piezoelectric F-Peptide Hydrogels Enable *In Situ* Stem Cell and Immune Niche Modulation for Cartilage Regeneration"

### Supplementary methods

#### HPLC method

The solvent gradient was as follows: solvent A, aqueous solution with 0.1% formic acid; solvent B, acetonitrile with 0.1% formic acid. The data were acquired over 25 min, with acetonitrile concentration increasing from 10% to 100%, and the flow rate was set to 0.3 $mL\cdot min^{-1}$.

#### Synthesis of FRET-1 peptide

Peptide synthesis is carried out using the solid-phase synthesis method with CTC resin and Fmoc-Lys (Dabcyl)-OH, and then purified by RP-HPLC.

#### qRT-PCR

Total RNA was extracted from cells or tissues using the RNAprep Pure Micro Kit (TIANGEN, Beijing, China) following the manufacturer's protocol. cDNA was synthesized using the NovoScriptPlus All-in-one 1st Strand cDNA Synthesis SuperMix (Novoprotein, Suzhou, China). Quantitative real-time PCR (qRT-PCR) was performed on a QuantStudio$^{TM}$ 1 System (Thermo Fisher Scientific, USA) using NovoStart SYBR qPCR SuperMix Plus (Novoprotein). Gene expression was normalized to GAPDH and calculated using the $2^{\wedge-\Delta\Delta Ct}$ method. All reactions were performed in at least triplicate. Primer sequences are listed in **Supplementary table S1.**

#### Western blot

Cells were lysed in RIPA buffer containing PMSF (R0010, Solarbio Science & Technology) and phosphatase inhibitors, and centrifuged at 13,000 rpm for 15 min. Protein concentration in the supernatant was determined by BCA assay. Samples were mixed with loading buffer, separated by 8% or 10% SDS-PAGE, and transferred to PVDF membranes. After blocking with 5% non-fat milk in TBST for 2 h at room temperature, membranes were incubated with primary antibodies overnight at 4°C, followed by secondary antibodies for 2 h at room temperature. Blots were visualized using ECL and a gel imaging system. The following primary antibodies were used: anti-GAPDH antibody (A19056, Abcolonal), anti-COL2A1 antibody (EPR12268, Abcam), anti-SOX9 antibody (EPR14335-78, Abcam), anti-PIEZO2 antibody (26205-1-AP, Proteintech). The following secondary antibodies were used: HRP-conjugated goat anti-mouse or goat anti-rabbit IgG (H+L)

(AS003, AS014, Abcolonal).

**Flow cytometry**

Cells were digested and centrifuged at 1,500 rpm for 5 min and then stored in PBS solution containing 1 % PFA. The cells were resuspended with 0.2 % Tween 20 and allowed to stand for 10 min. Subsequently, the cells were washed twice with PBS, the supernatant was removed, and the cells were blocked with 5 % BSA for 30 min. Then, the cells were incubated with primary antibodies for 30 min at room temperature, washed three times by PBS and then resuspended in 300 μL PBS. Detection was performed using Agilent NovoCyte-D2060R (Agilent Technologies Inc, CA, USA).

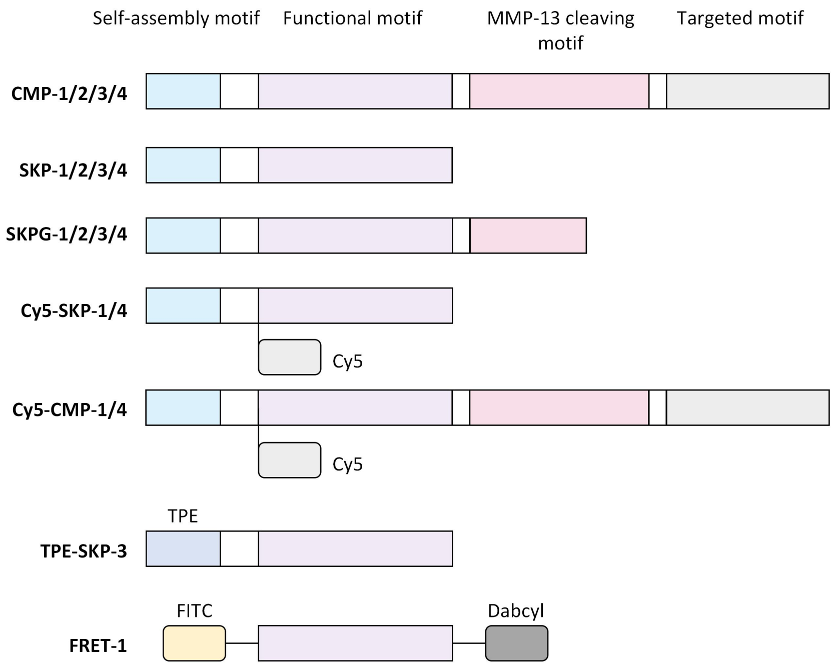

**Supplementary Fig. 1. Schematic representation of the molecular structures of all peptides used in this**

**work.**

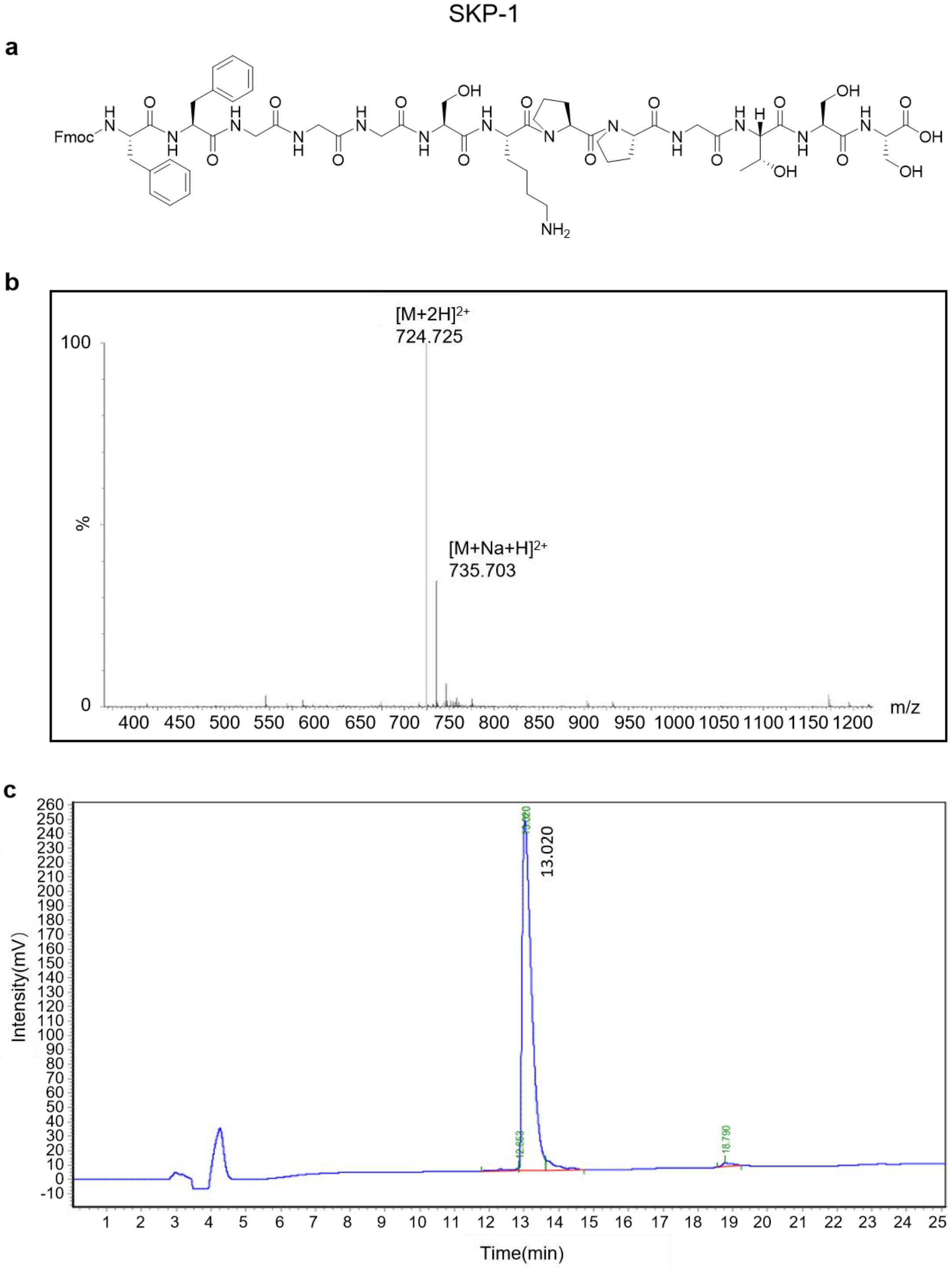

**Supplementary Fig. 2. HPLC and MS analysis of the SKP-1**.

**a**, Structural formula of SKP-1 peptide. **b**, LC/MS of SKP-1 peptide. **c**, HPLC of SKP-1 peptide. The purity of SKP-1 was greater than 90%.

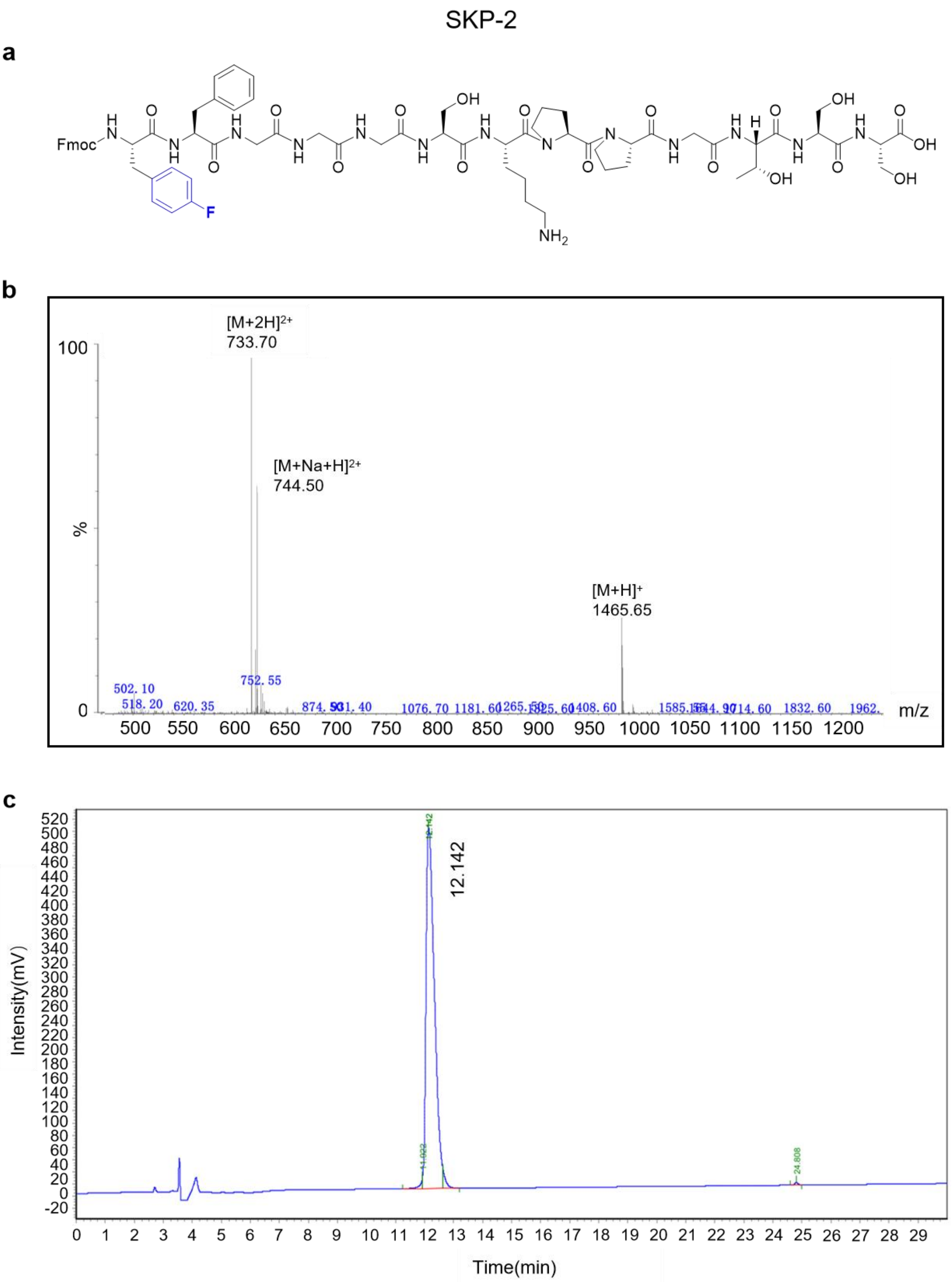

**Supplementary Fig. 3. HPLC and MS analysis of the SKP-2**.

**a**, Structural formula of SKP-2 peptide. **b**, LC/MS of SKP-2 peptide. **c**, HPLC of SKP-2 peptide. The purity of SKP-2 was greater than 90%.

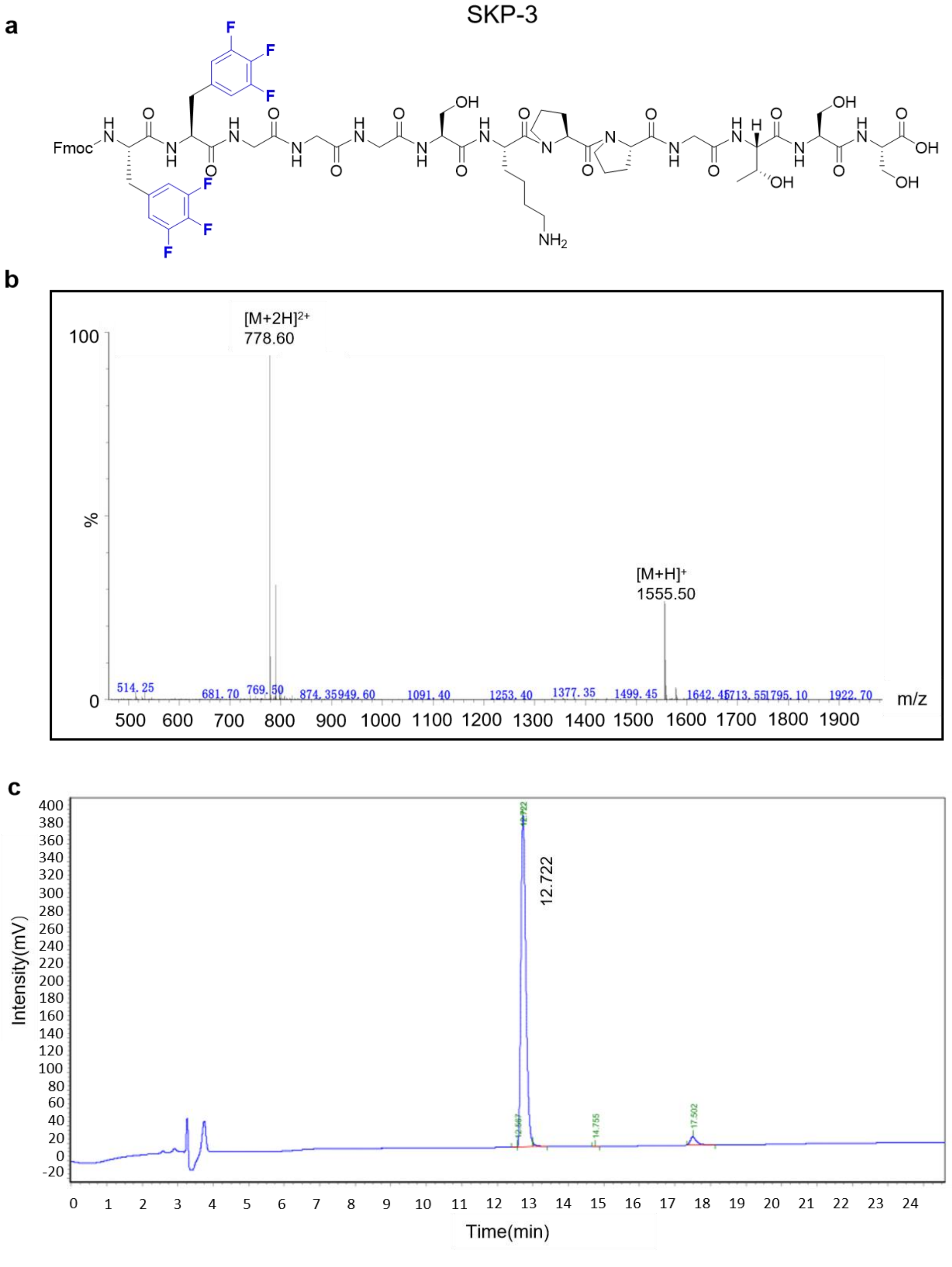

**Supplementary Fig. 4. HPLC and MS analysis of the SKP-3**.

**a**, Structural formula of SKP-3 peptide. **b**, LC/MS of SKP-3 peptide. **c**, HPLC of SKP-3 peptide. The purity of SKP-3 was greater than 90%.

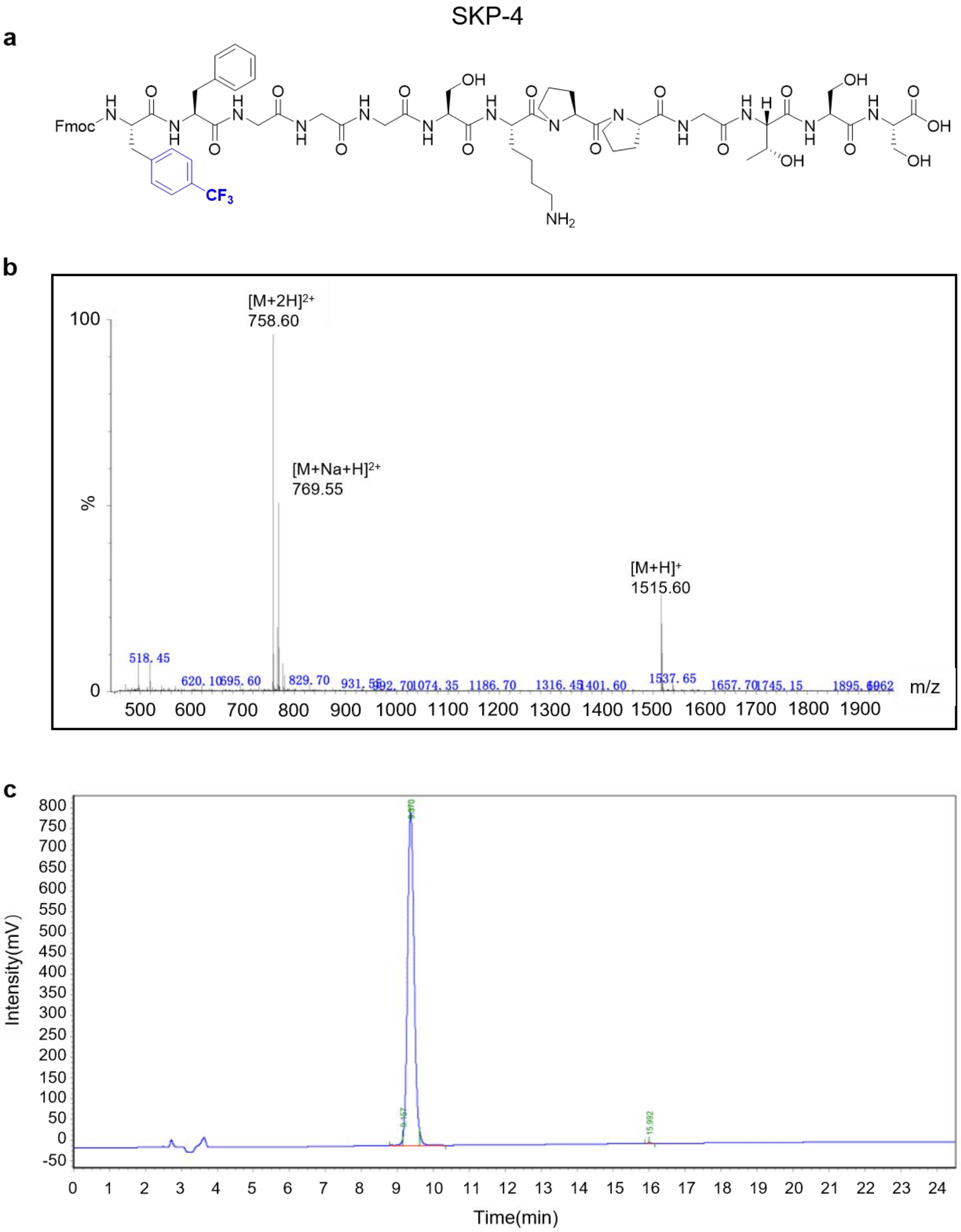

**Supplementary Fig. 5. HPLC and MS analysis of the SKP-4**.

**a**, Structural formula of SKP-4 peptide. **b**, LC/MS of SKP-4 peptide. **c**, HPLC of SKP-4 peptide. The purity of SKP-4 was greater than 90%.

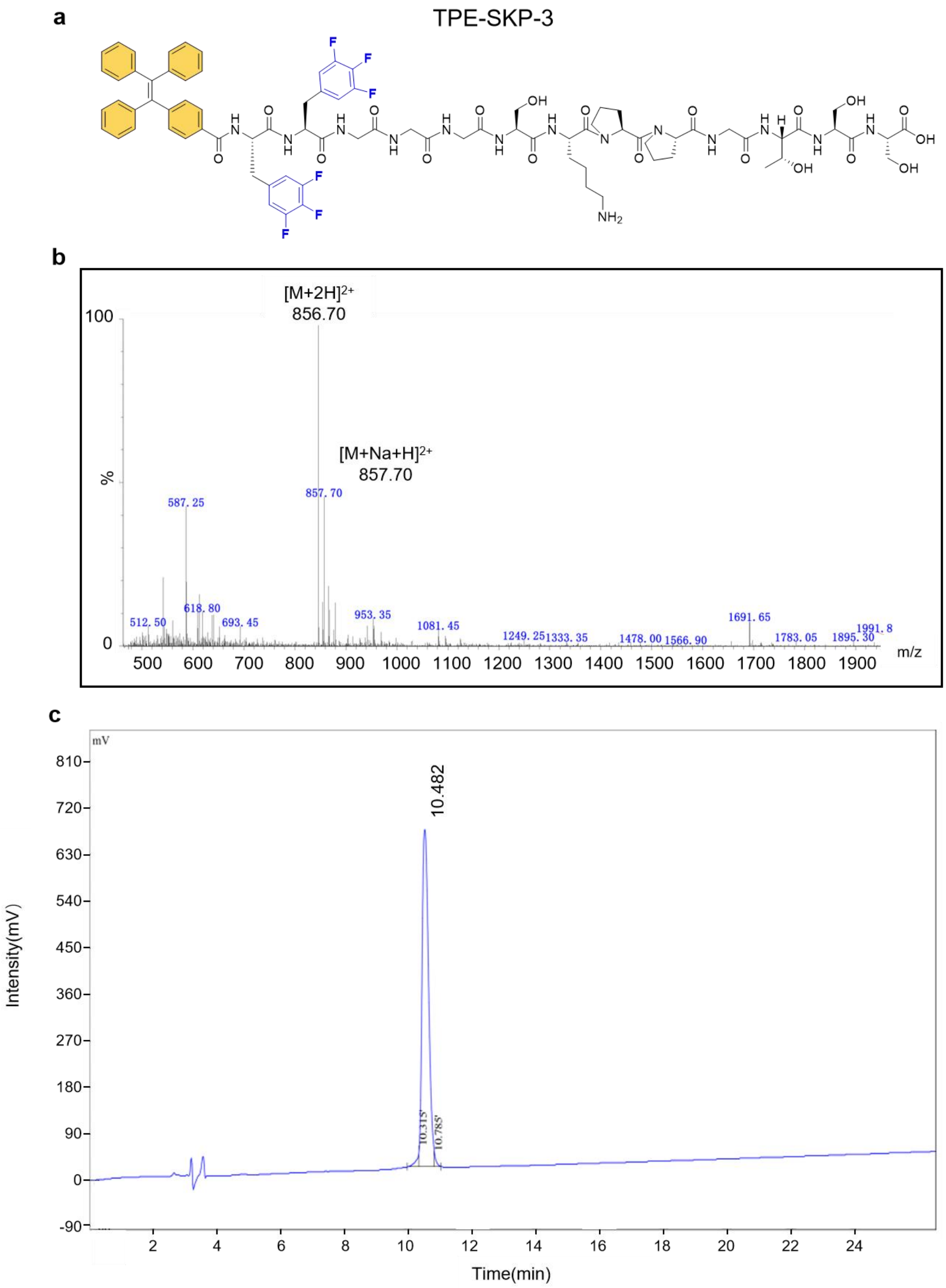

**Supplementary Fig. 6. HPLC and MS analysis of the TPE-SKP-3.**

**a**, Structural formula of TPE-SKP-3 peptide. **b**, LC/MS of TPE-SKP-3 peptide. **c**, HPLC of TPE-SKP-3 peptide. The purity of TPE-SKP-3 was greater than 90%.

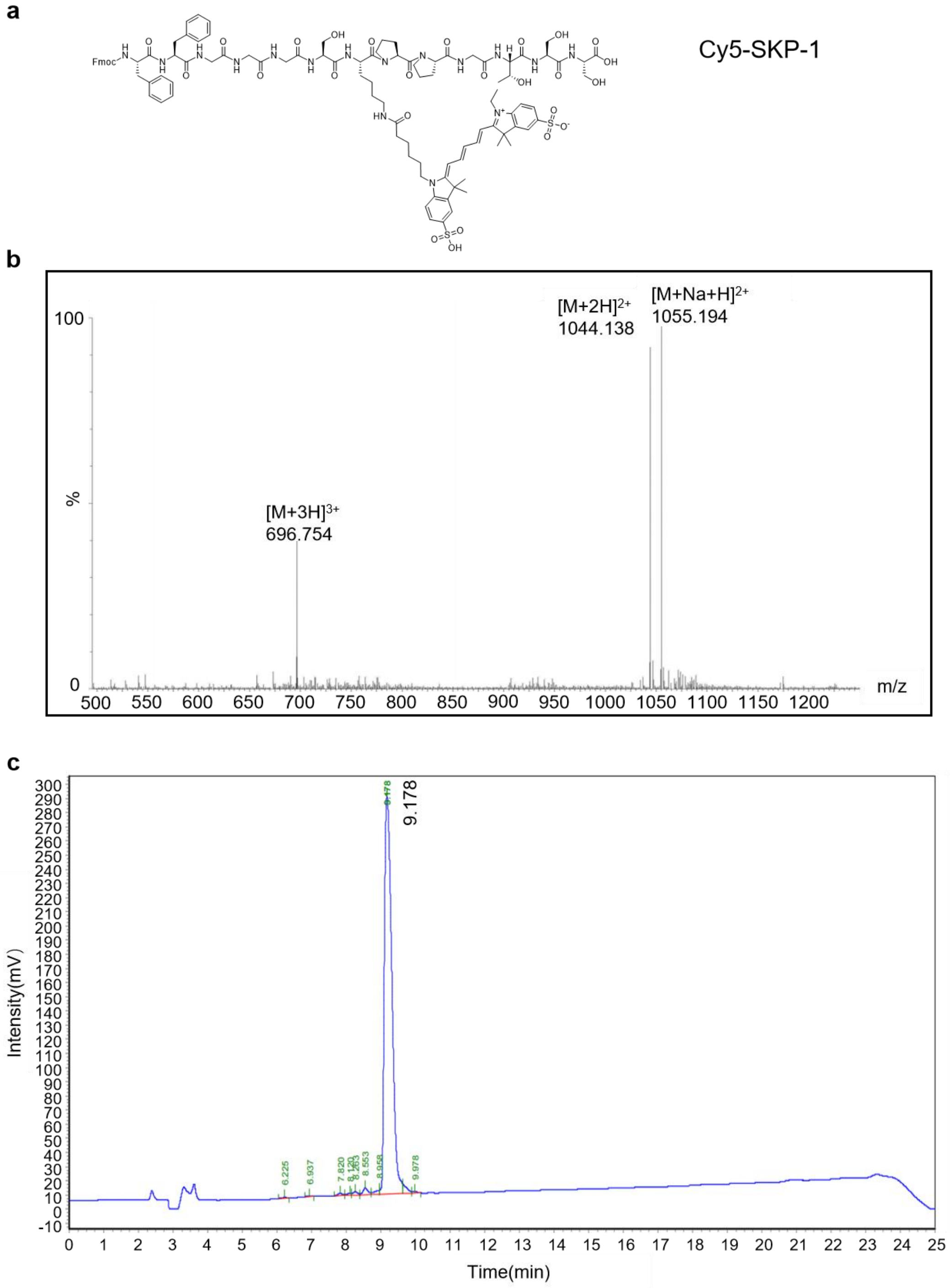

**Supplementary Fig. 7. HPLC and MS analysis of the Cy5-SKP-1.**

**a**, Structural formula of Cy5-SKP-1 peptide. **b**, LC/MS of Cy5-SKP-1 peptide. **c**, HPLC of Cy5-SKP-1 peptide. The purity of Cy5-SKP-1 was greater than 90%.

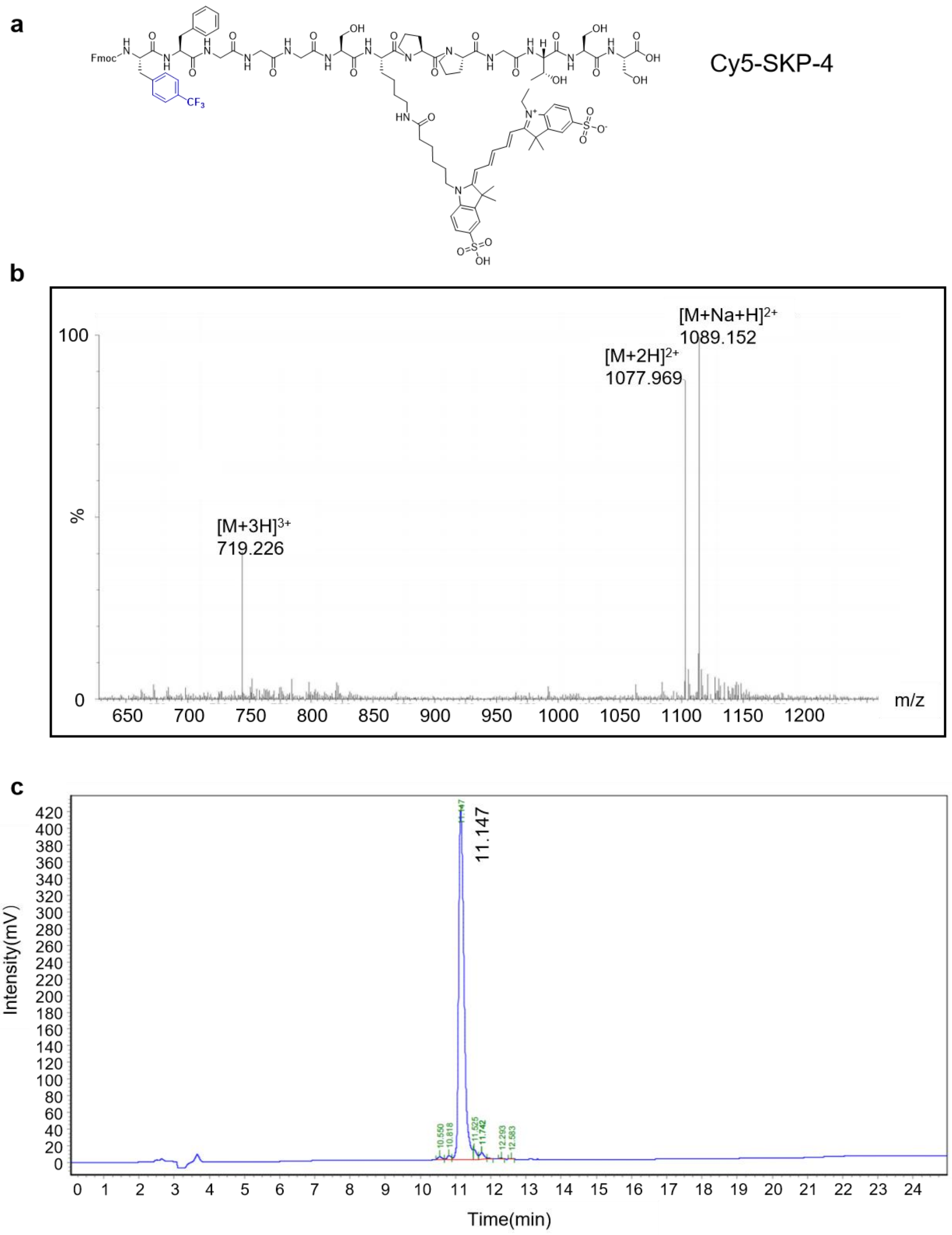

**Supplementary Fig. 8. HPLC and MS analysis of the Cy5-SKP-4.**

**a**, Structural formula of Cy5-SKP-4 peptide. **b**, LC/MS of Cy5-SKP-4 peptide. **c**, HPLC of Cy5-SKP-4 peptide. The purity of Cy5-SKP-4 was greater than 90%.

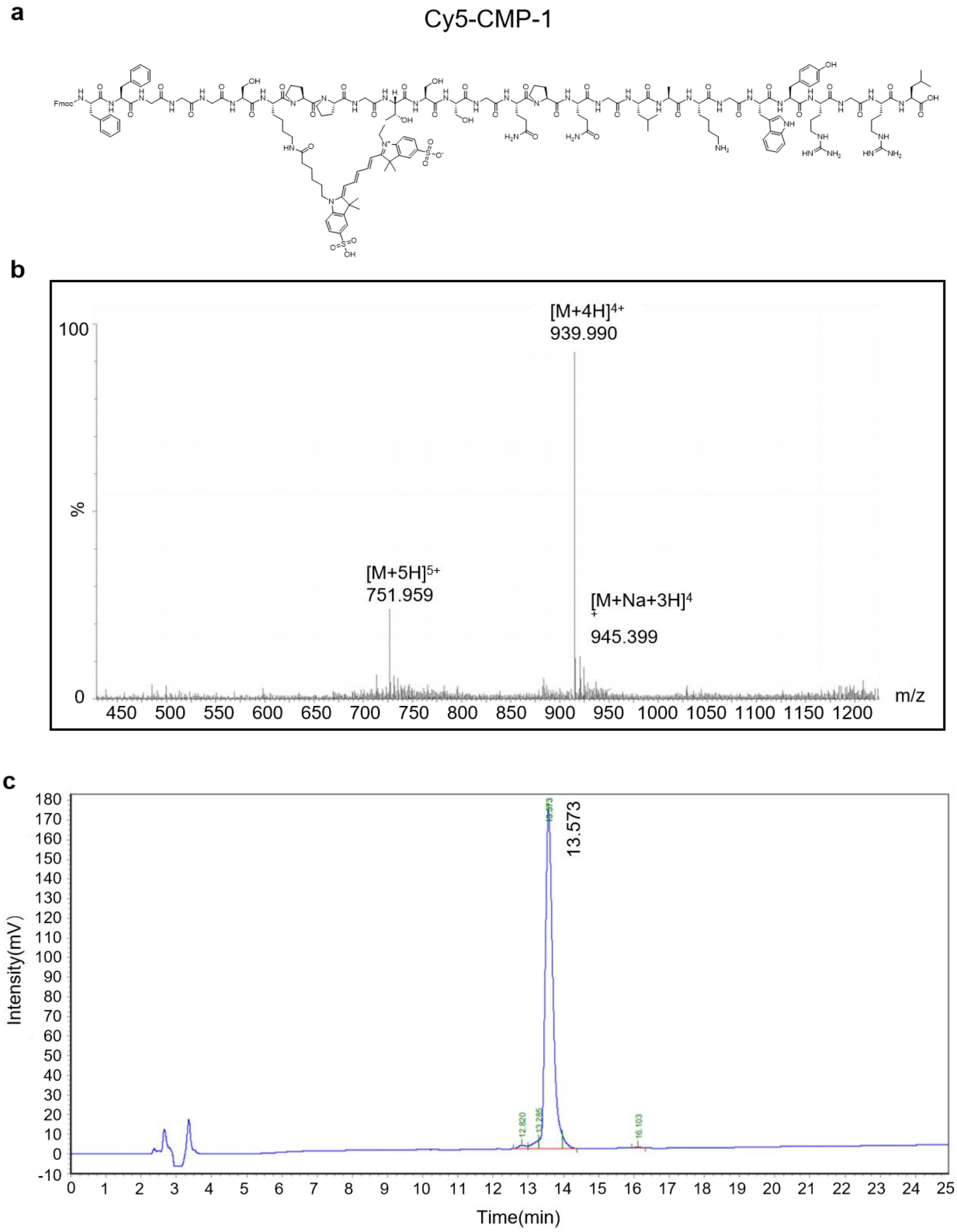

**Supplementary Fig. 9. HPLC and MS analysis of the Cy5-CMP-1.**

**a**, Structural formula of Cy5-CMP-1 peptide. **b**, LC/MS of Cy5-CMP-1 peptide. **c**, HPLC of Cy5-CMP-1 peptide. The purity of Cy5-CMP-1 was greater than 90%.

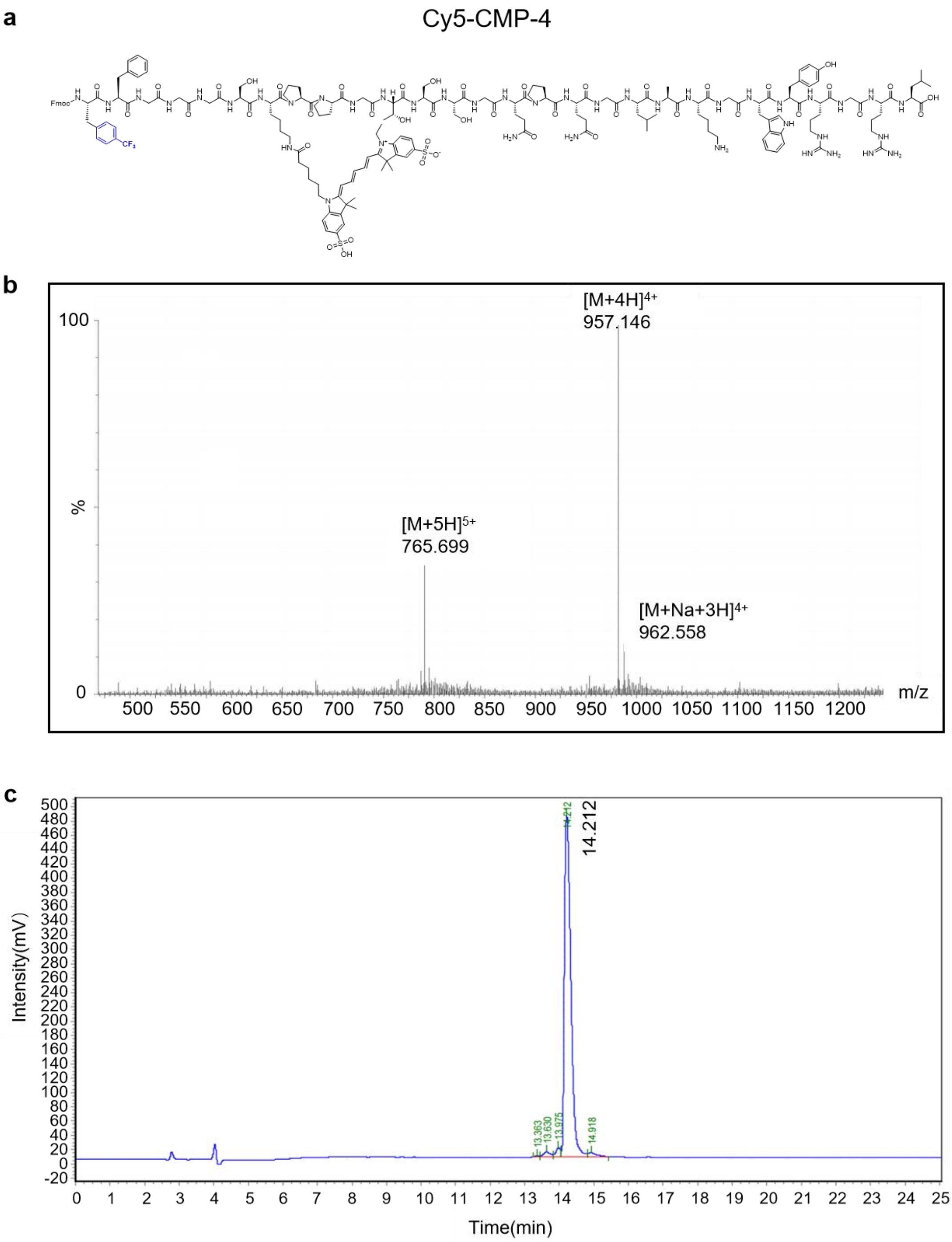

**Supplementary Fig. 10. HPLC and MS analysis of the Cy5-CMP-4.**

**a**, Structural formula of Cy5-CMP-4 peptide. **b**, LC/MS of Cy5-CMP-4 peptide. **c**, HPLC of Cy5-CMP-4 peptide. The purity of Cy5-CMP-4 was greater than 90%.

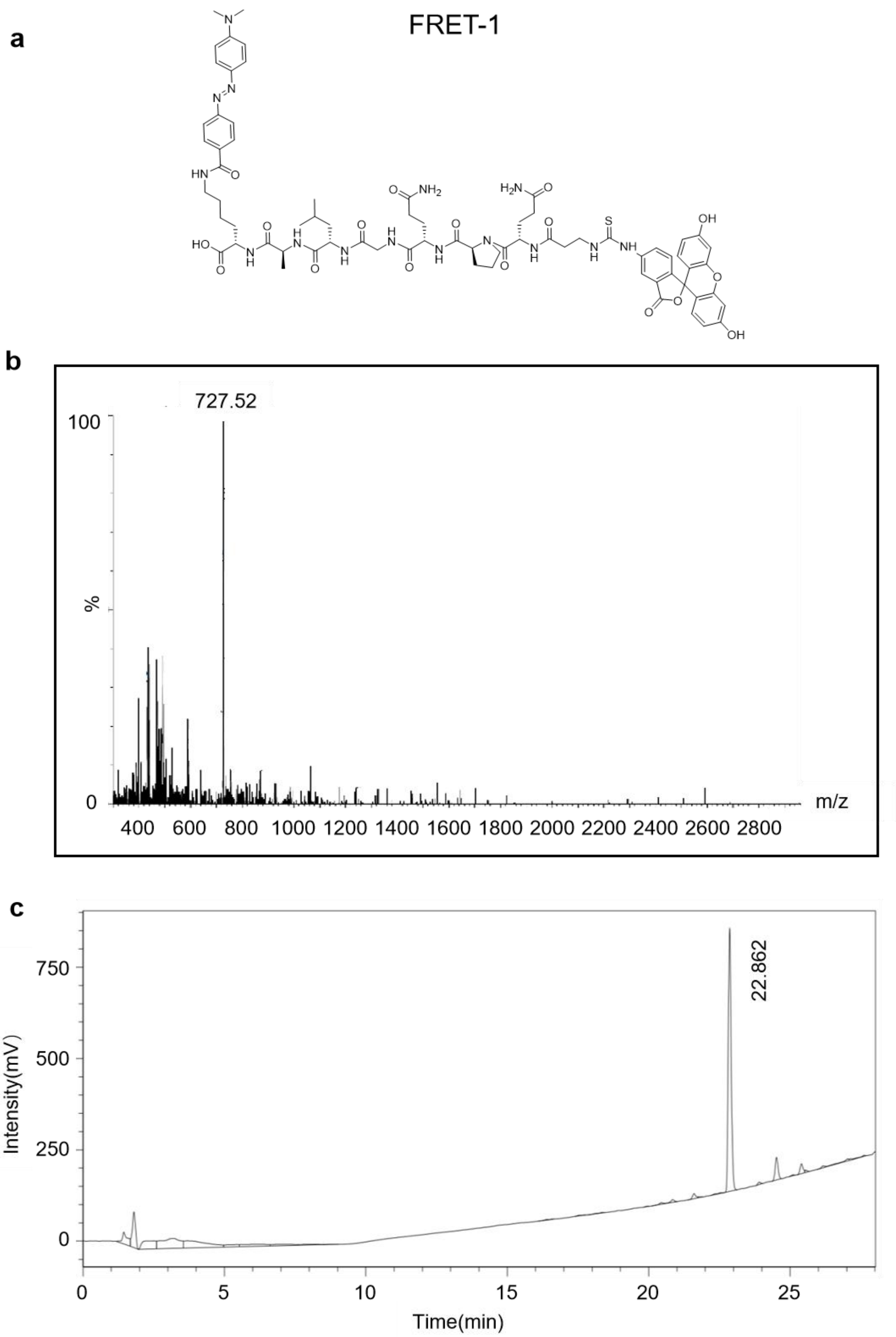

**Supplementary Fig. 11. HPLC and MS analysis of the FRET-1.**

**a**, Structural formula of FRET-1 peptide. **b**, LC/MS of FRET-1 peptide. **c**, HPLC of FRET-1 peptide. The purity of FRET-1 was greater than 90%.

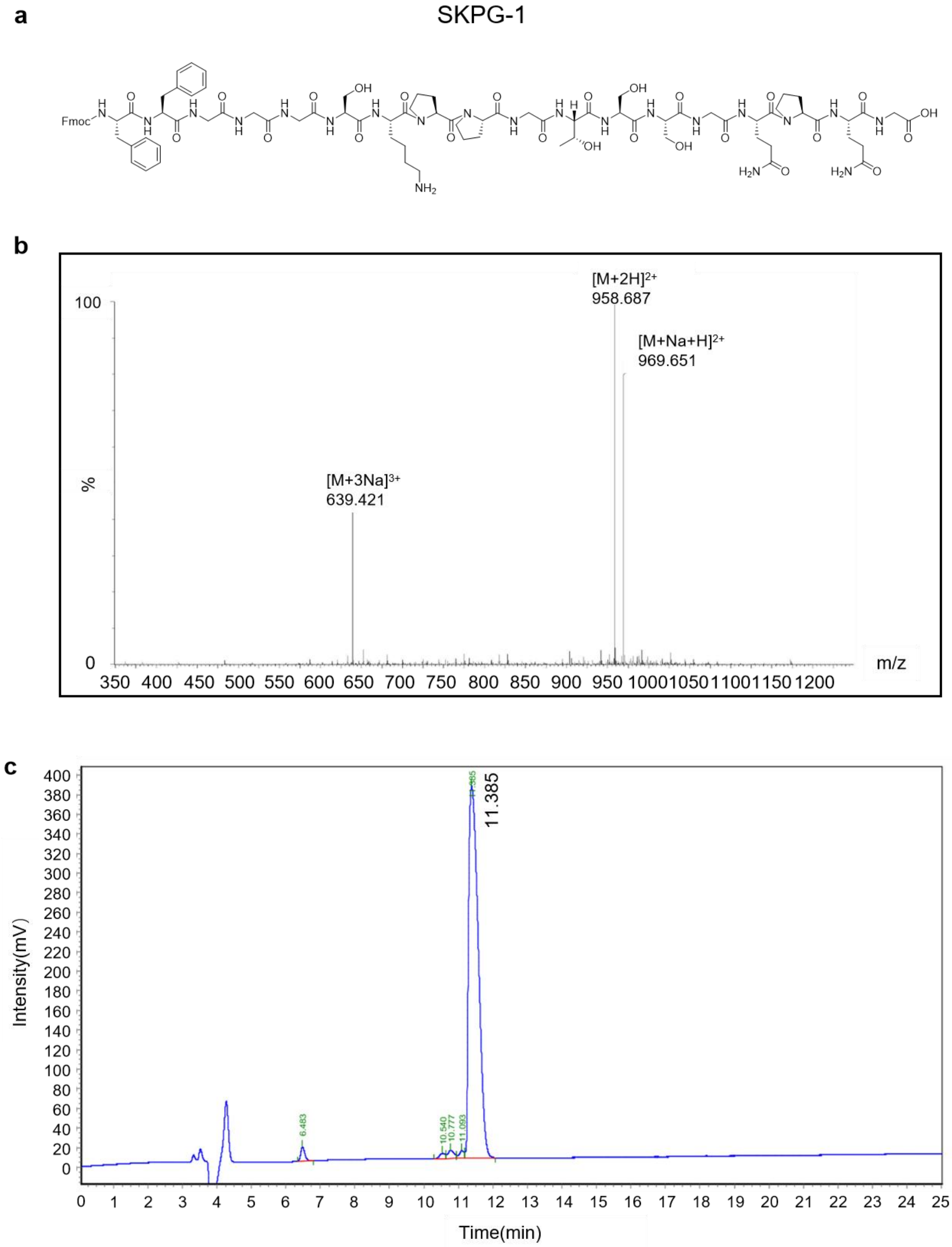

**Supplementary Fig. 12. HPLC and MS analysis of the SKPG-1.**

**a**, Structural formula of SKPG-1 peptide. **b**, LC/MS of SKPG-1 peptide. **c**, HPLC of SKPG-1 peptide. The purity of SKPG-1 was greater than 90%.

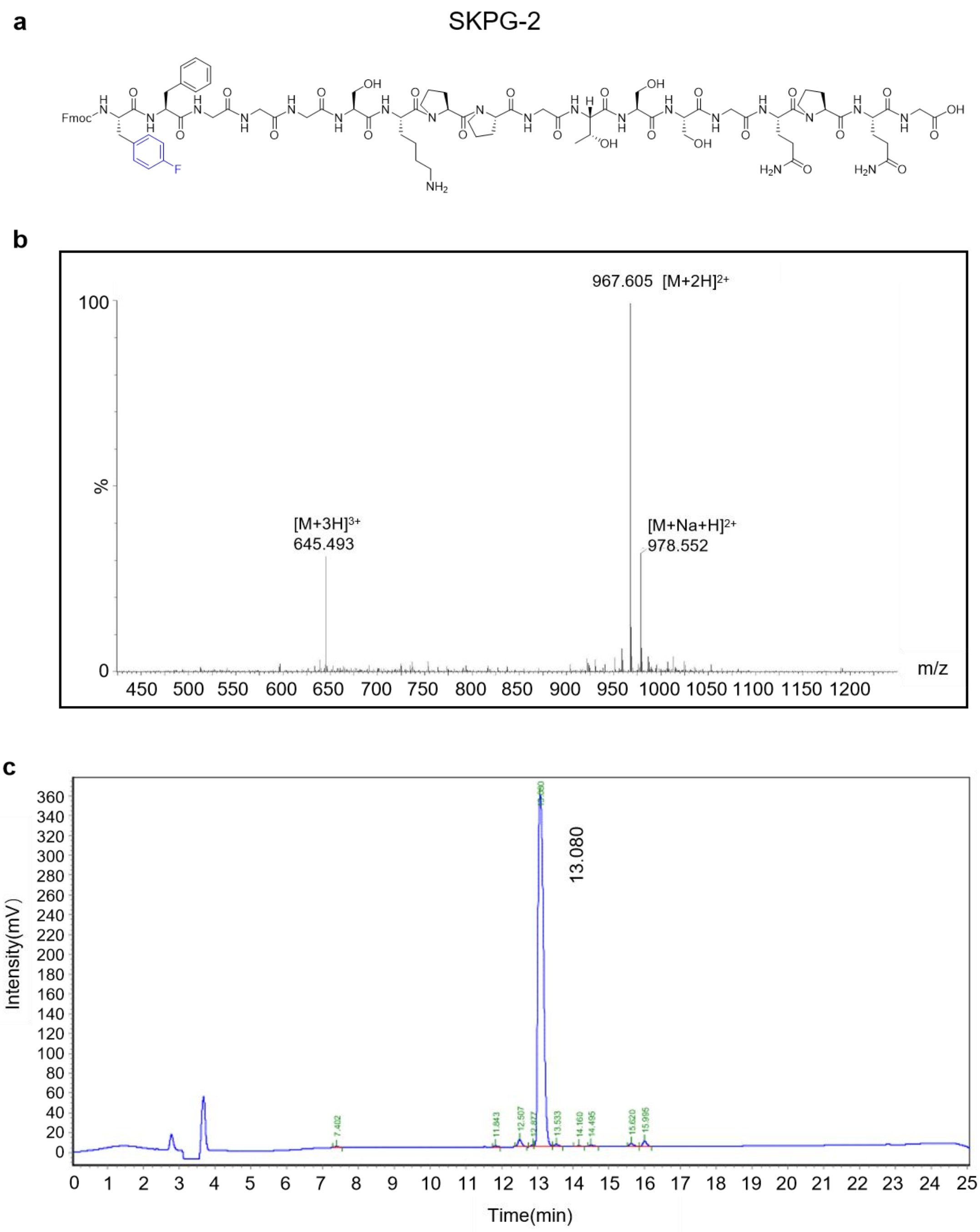

**Supplementary Fig. 13. HPLC and MS analysis of the SKPG-2.**

**a**, Structural formula of SKPG-2 peptide. **b**, LC/MS of SKPG-2 peptide. **c**, HPLC of SKPG-2 peptide. The purity of SKPG-2 was greater than 90%.

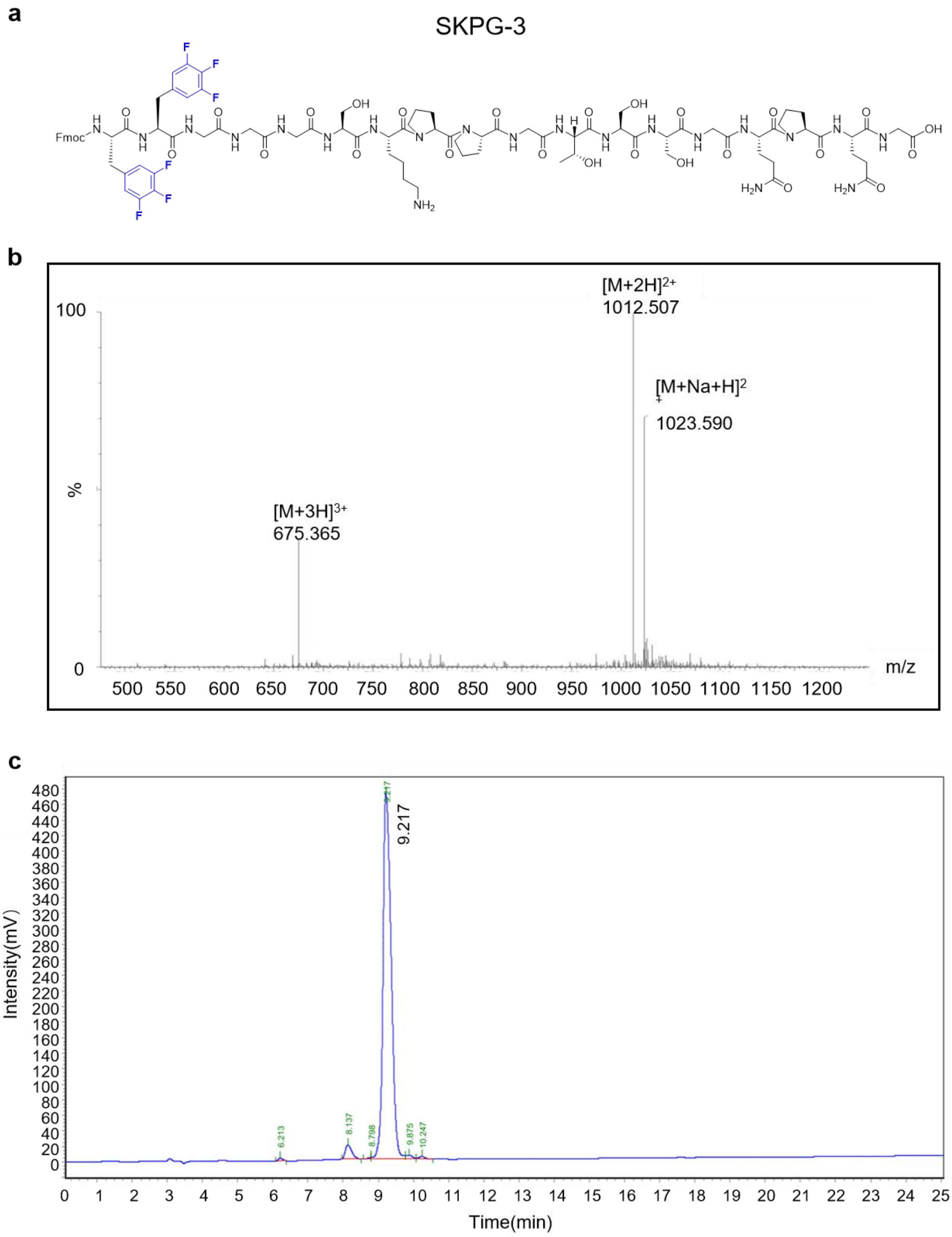

**Supplementary Fig. 14. HPLC and MS analysis of the SKPG-3.**

**a**, Structural formula of SKPG-3 peptide. **b**, LC/MS of SKPG-3 peptide. **c**, HPLC of SKPG-3 peptide. The purity of SKPG-3 was greater than 90%.

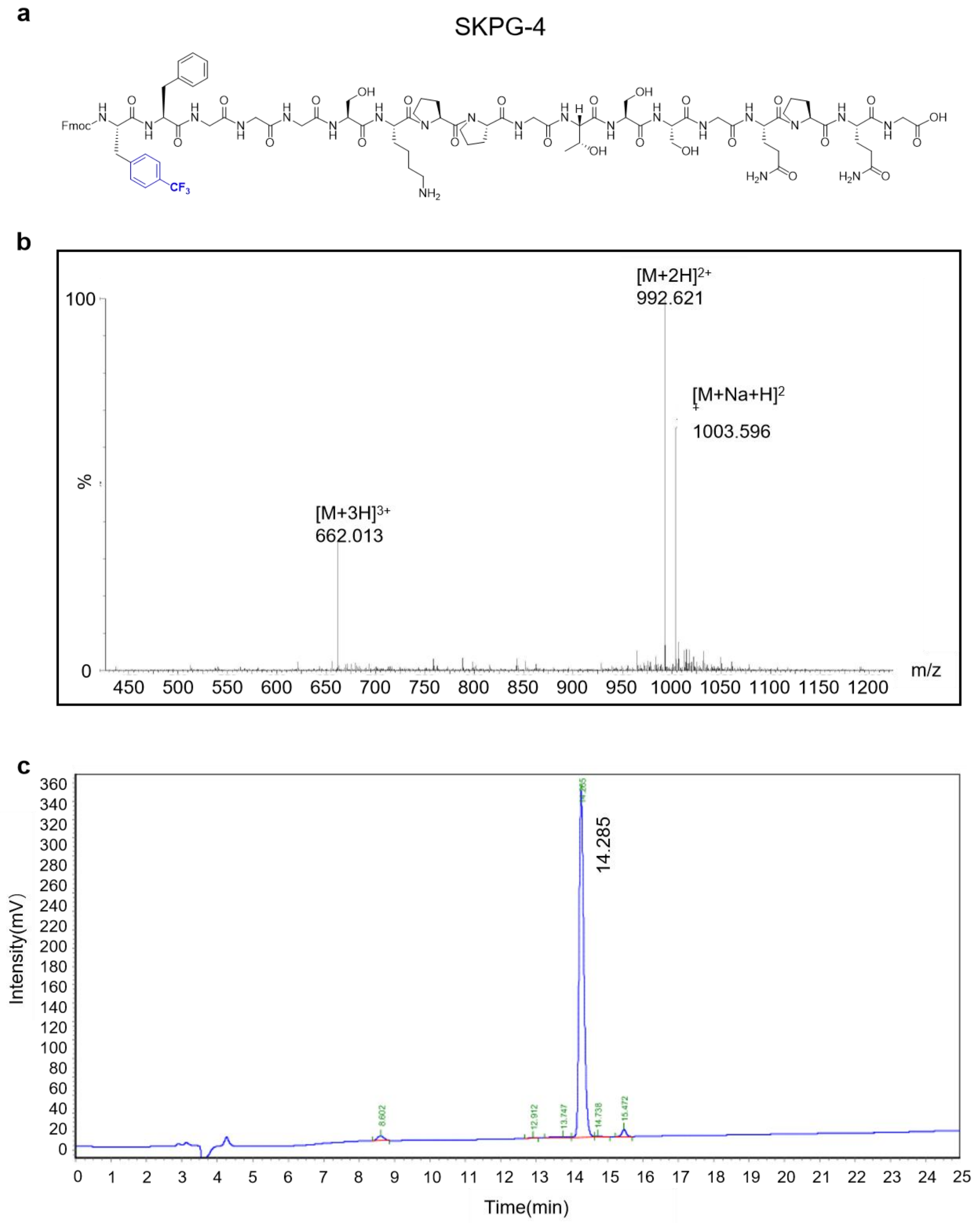

**Supplementary Fig. 15. HPLC and MS analysis of the SKPG-4.**

**a**, Structural formula of SKPG-4 peptide. **b**, LC/MS of SKPG-4 peptide. **c**, HPLC of SKPG-4 peptide. The purity of SKPG-4 was greater than 90%.

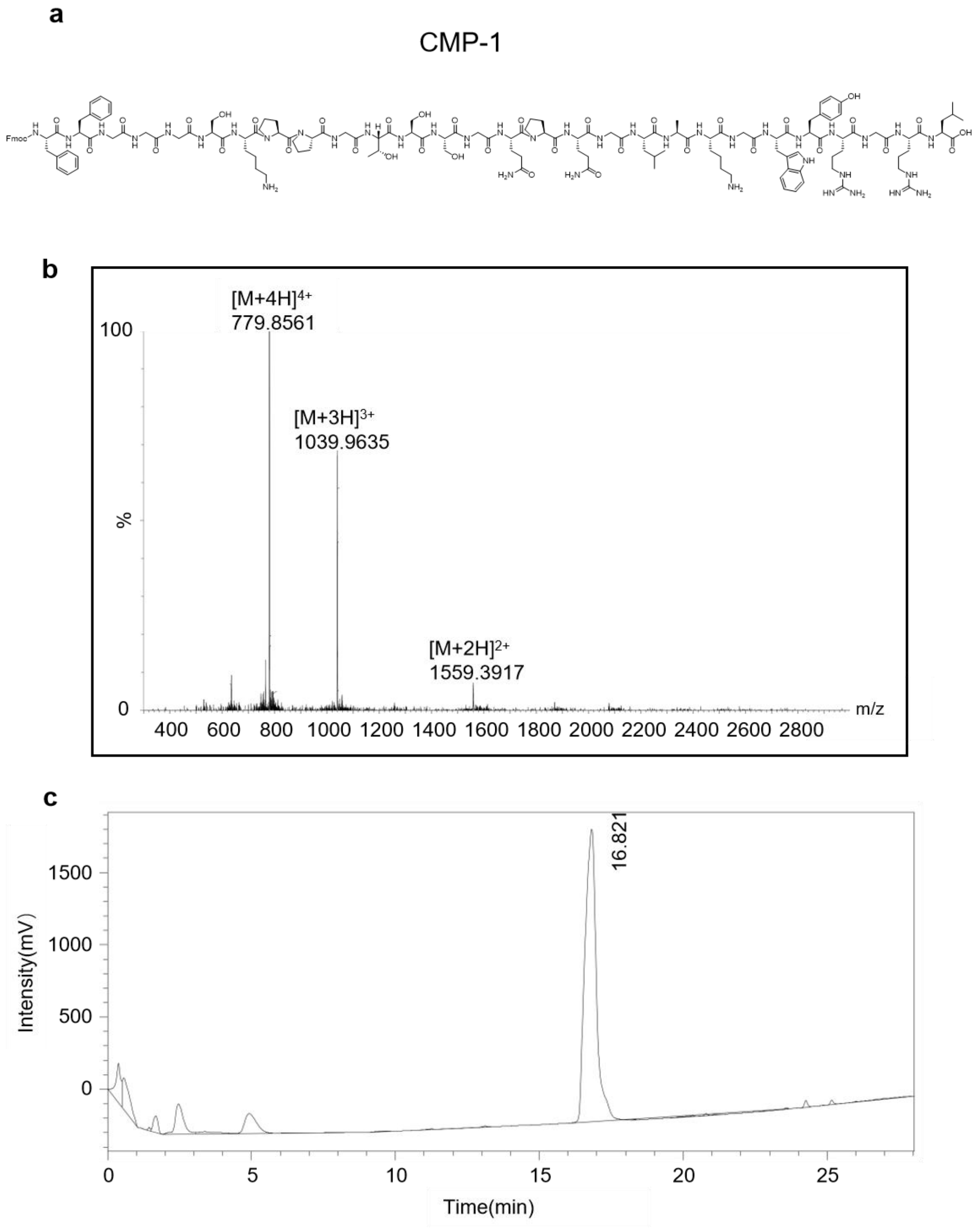

**Supplementary Fig. 16. HPLC and MS analysis of the CMP-1.**

**a**, Structural formula of CMP-1 peptide. **b**, LC/MS of CMP-1 peptide. **c**, HPLC of CMP-1 peptide. The purity of CMP-1 was greater than 90%.

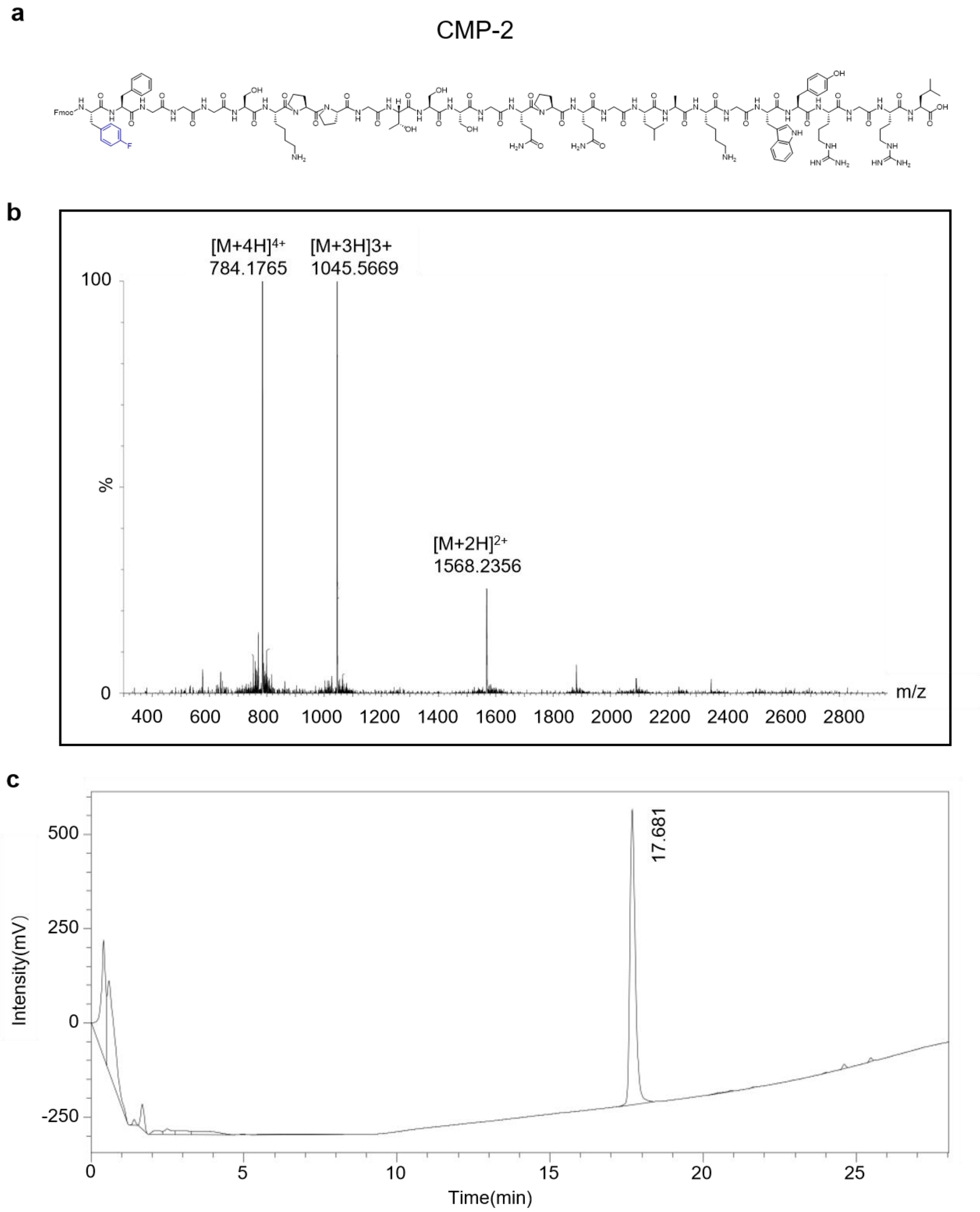

**Supplementary Fig. 17. HPLC and MS analysis of the CMP-2.**

**a**, Structural formula of CMP-2 peptide. **b**, LC/MS of CMP-2 peptide. **c**, HPLC of CMP-2 peptide. The purity of CMP-2 was greater than 90%.

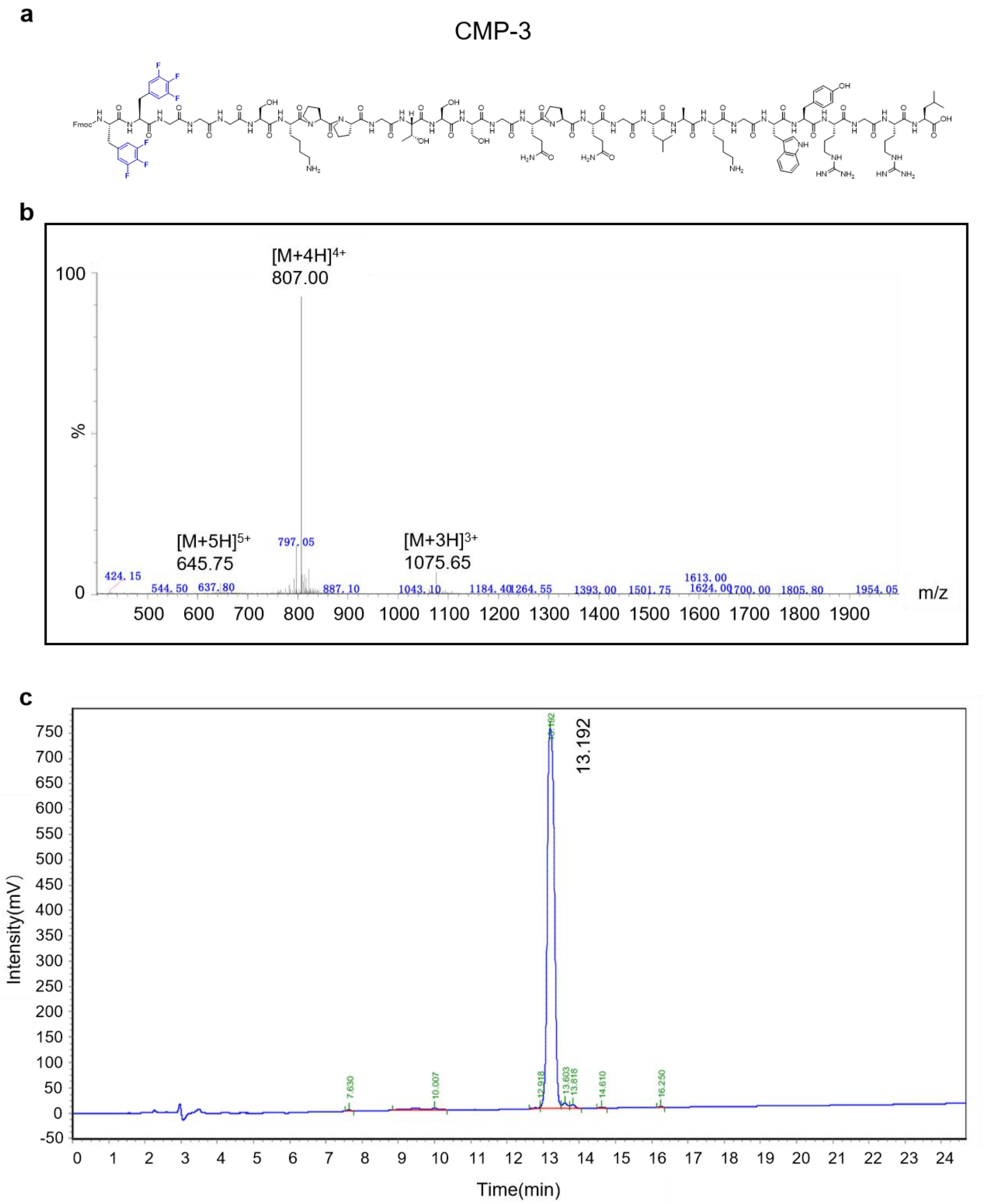

**Supplementary Fig. 18. HPLC and MS analysis of the CMP-3.**

**a**, Structural formula of CMP-3 peptide. **b**, LC/MS of CMP-3 peptide. **c**, HPLC of CMP-3 peptide. The purity of CMP-3 was greater than 90%.

c

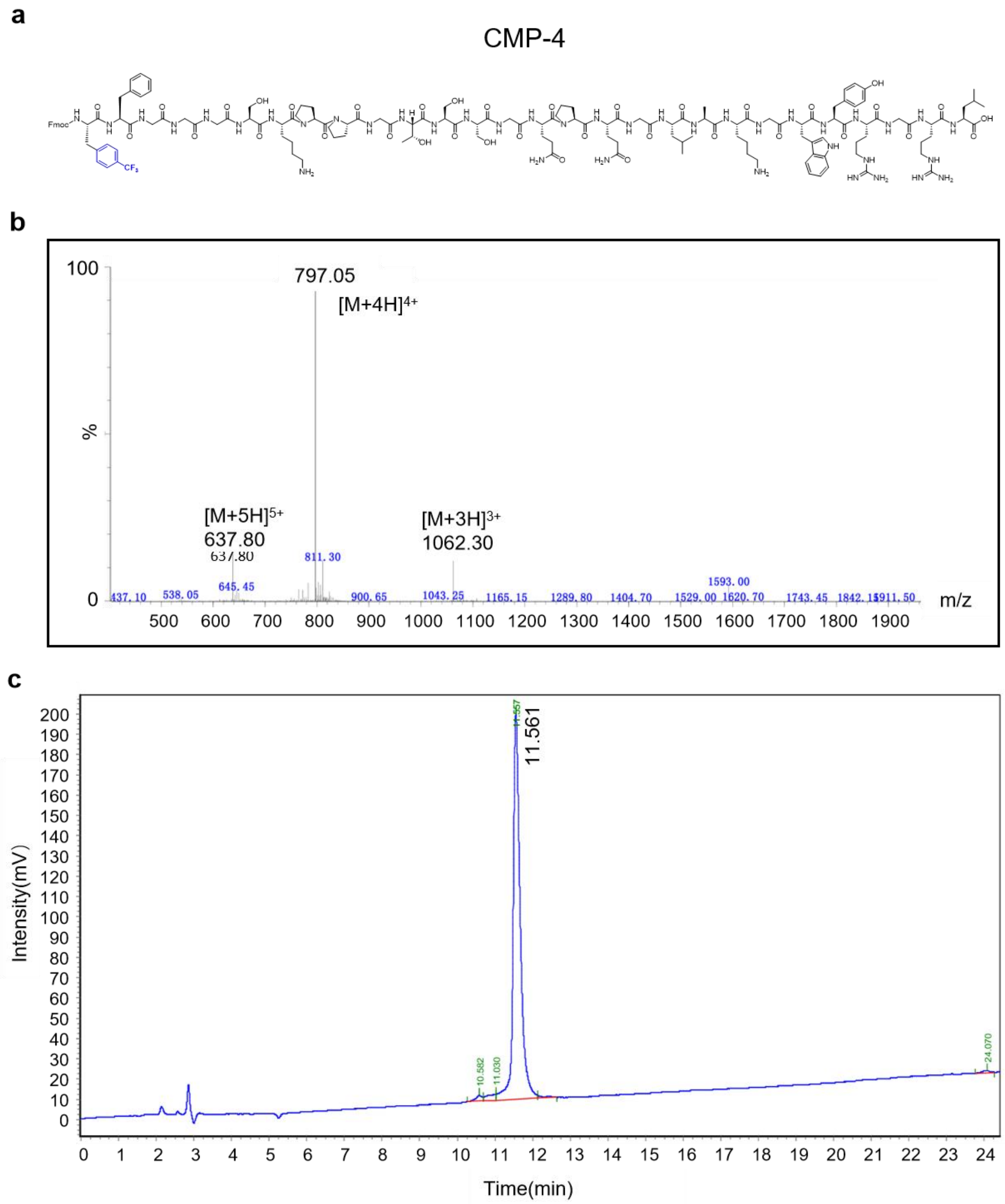

**Supplementary Fig. 19. HPLC and MS analysis of the CMP-4.**

**a**, Structural formula of CMP-4 peptide. **b**, LC/MS of CMP-4 peptide. **c**, HPLC of CMP-4 peptide. The purity of CMP-4 was greater than 90%.

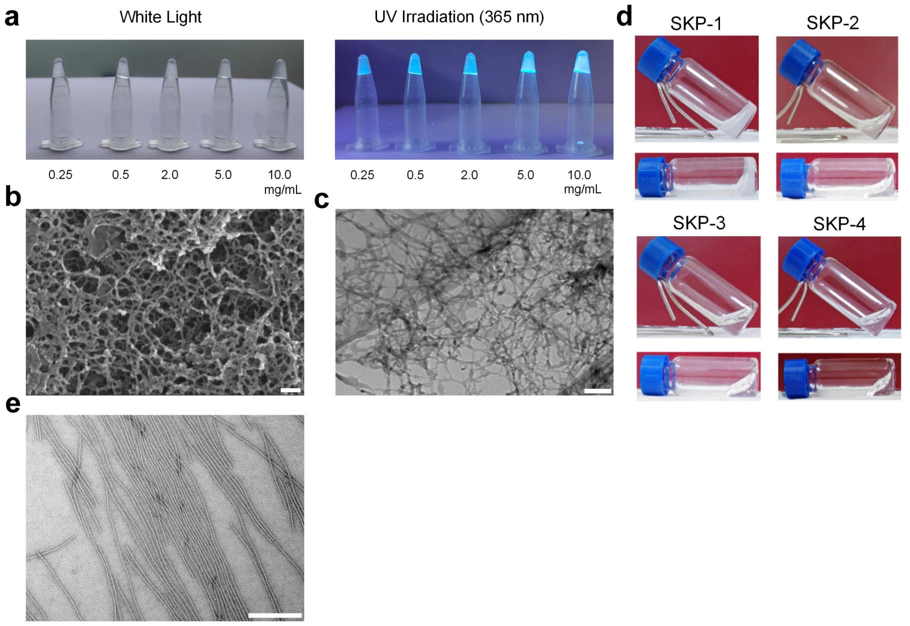

**Supplementary Fig. 20. Preparation and characterization of SKP peptide hydrogels with different fluorination modifications.**

**a**, Gelation results under white light (left) and aggregation-induced emission results under 365 nm UV irradiation (right) for TPE-SKP-3 peptides at concentrations of 0.25, 0.5, 2.0, 5.0, and 10.0 mg·mL$^{-1}$. **b**, SEM image of the TPE-SKP-3 hydrogel at 10.0 mg/mL. Scale bar: 200 nm. **c**, TEM image of the TPE-SKP-3 hydrogel at 10.0 mg·mL$^{-1}$, scale bar: 200 nm. **d**, Photographs of four different SKP hydrogels formed by 10.0 mg·mL$^{-1}$ of peptide in dd$H_2O$. **e**, TEM image of SKP-1 peptide. Scale bar: 200 nm.

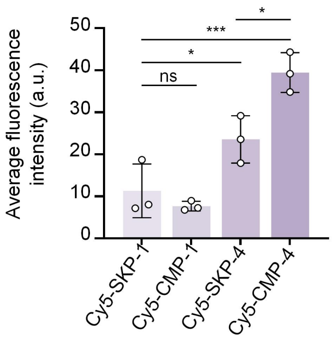

**Supplementary Fig. 21. Quantification of Cy5-labeled peptide (red) fluorescence in cartilage.** Statistical significances were calculated via one-way ANOVA. ns means no significance, *$p < 0.05$, ***$p < 0.001$.

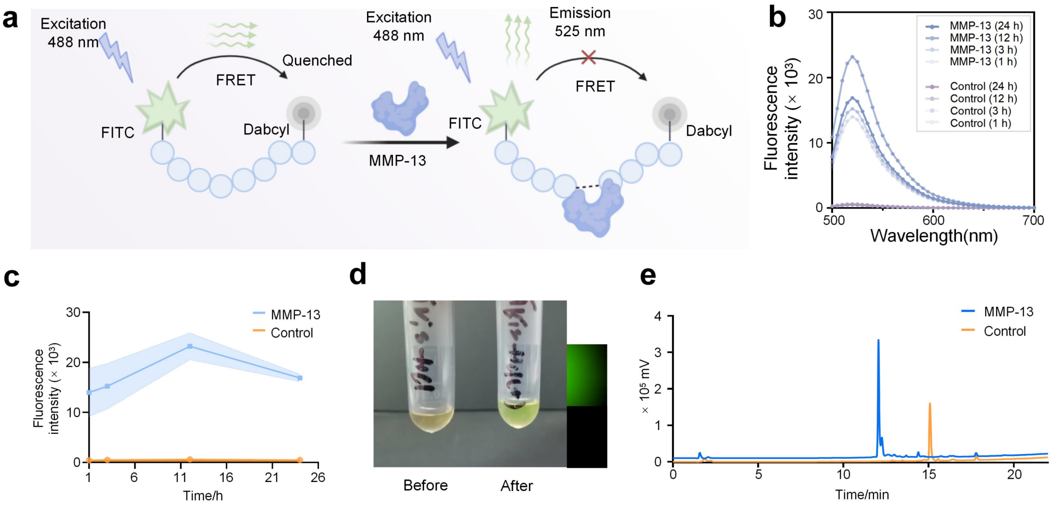

**Supplementary Fig. 22. Enzyme-Digestion Experiment Results for FRET of FRET-1 Peptide. a**, Working principle of FRET to detect MMP-13 enzymatic cleavage of peptides. This figure was created using BioRender. **b**, The fluorescence emission spectrum of the FRET-1 peptide at 1, 3, 12 and 24 h after the addition of MMP-13 enzyme or identical Tris-HCl buffer. **c**, The fluorescence intensity at 525 nm of the FRET-1 peptide at 1, 3, 12 and 24 h after the addition of MMP-13 enzyme or identical Tris-HCl buffer (Control). **d**, Color/fluorescence changes of FRET-1 under visible light and 488 nm excitation light before and after enzymatic digestion. **e**, HPLC chromatograms of FRET-1 peptide before and after enzymatic digestion.

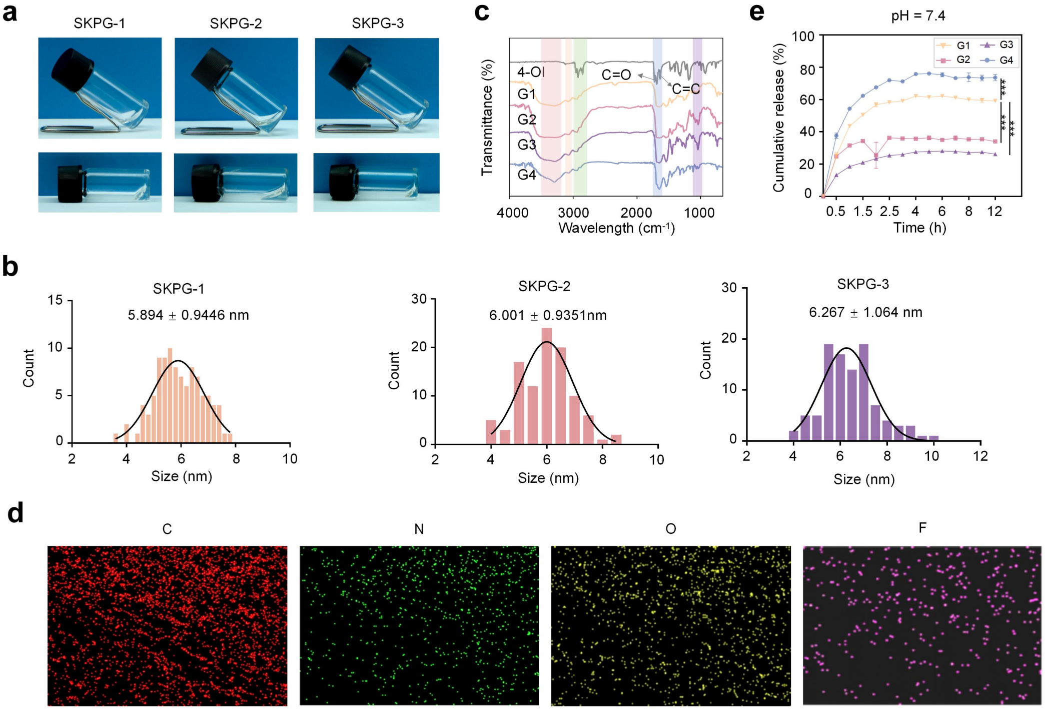

**Supplementary Fig. 23. Characterization of the enzymatically cleaved and drug-loaded SKPG-series**

**self-assembling peptide hydrogels.**

**a**, Photographs of different fluorinated SKPG hydrogels encapsulation of 4-OI formed by 10 mg·mL$^{-1}$ of peptide in ddH$_2$O. **b**, Histogram showing the diameter distribution of SKPG-1, SKPG-2 and SKPG-3 peptide nanofibers. The solid line represents a Gaussian fit to the data, n = 100. **c**, FTIR spectra of the enzymatically cleaved self-assembling hydrogels. **d**, EDS elemental mappings of G4. **e**, 4-OI sustained release curve of G1/2/3/4 in PBS buffer (pH 7.4) solution was determined by UPLC/MS, n = 3. Statistical significances were calculated via one-way ANOVA. *p < 0.05, **p < 0.01, ***p < 0.001; ns means no significance.

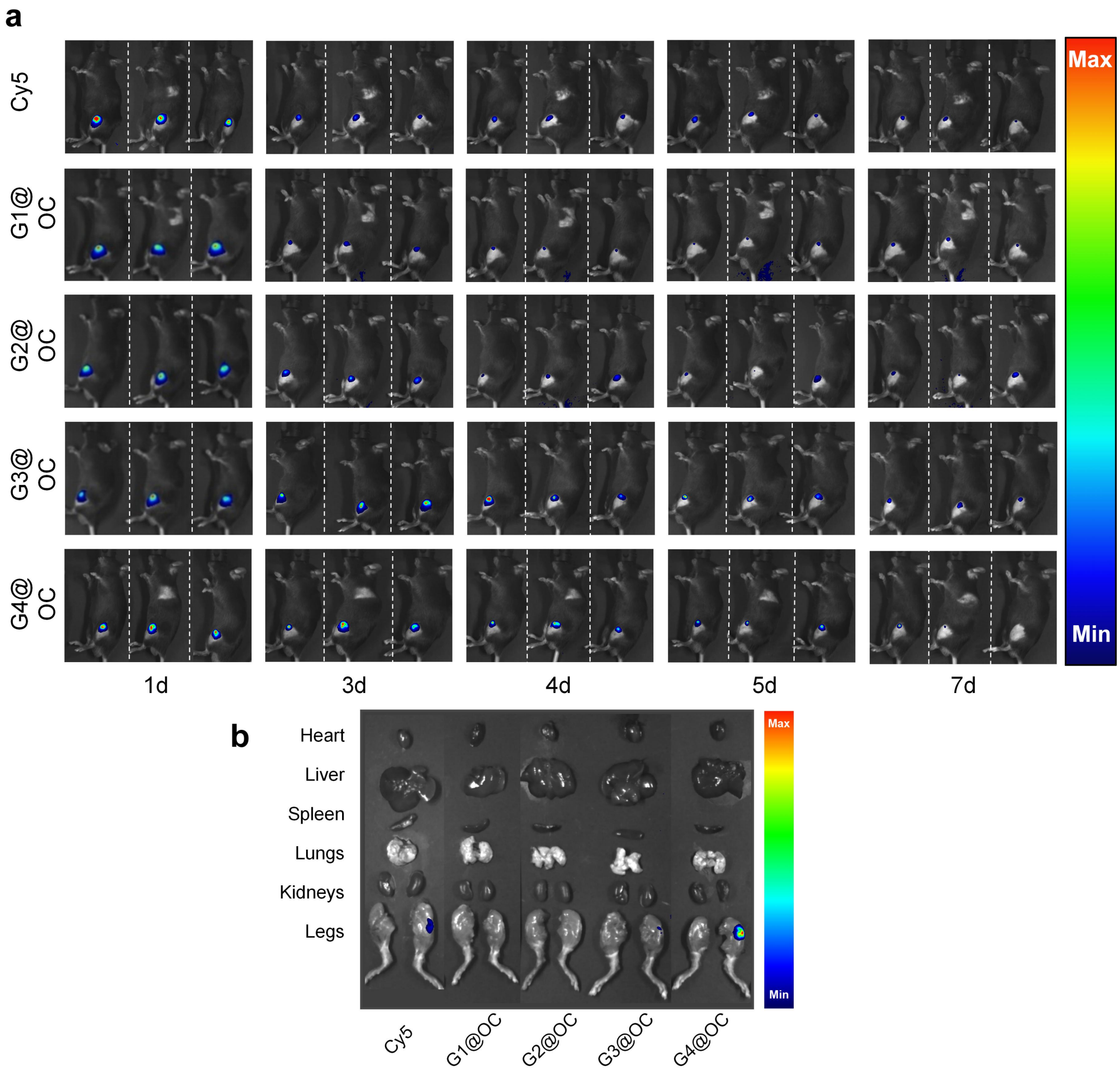

**Supplementary Fig. 24. *In vivo* retention of hydrogel in the knee joint.**

**a**, Joint cavity injection of Cy5 and SKPG hydrogels packaged Cy5 (G1@OC, G2@OC, G3@OC and G4@OC) followed by detection of biofluorescent signals at day 1, 3, 4, 5 and 7. **b**, *Ex vivo* organs fluorescence distribution results 3 days after *in situ* injection of Cy5 and SKPG hydrogels packaged Cy5 into the knee joint region.

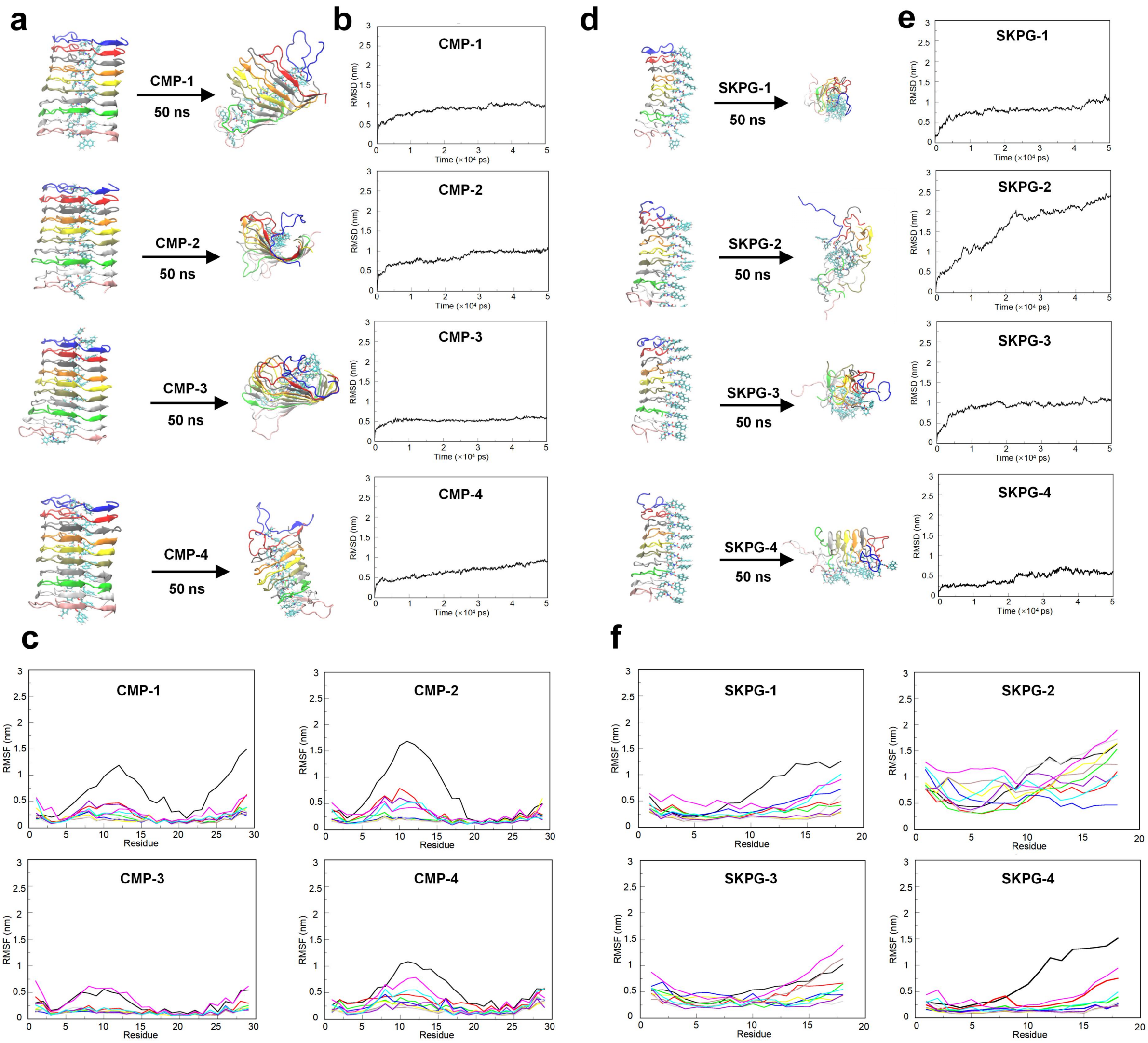

**Supplementary Fig. 25. AA-MD simulations of CMP and SKPG decamers over 50 ns.**

**a**, Structural representations of CMP-1/2/3/4 decamers after 50 ns of AA-MD simulation. **b**, RMSD trajectories of CMP-1/2/3/4 decamers over 50 ns. **c**, RMSF analysis of CMP-1/2/3/4 decamers over 50 ns. **d**, Structural representations of SKPG-1/2/3/4 decamers after 50 ns of AA-MD simulation. **e**, RMSD trajectories of SKPG-1/2/3/4 decamers over 50 ns. **f**, RMSF analysis of SKPG-1/2/3/4 decamers over 50 ns.

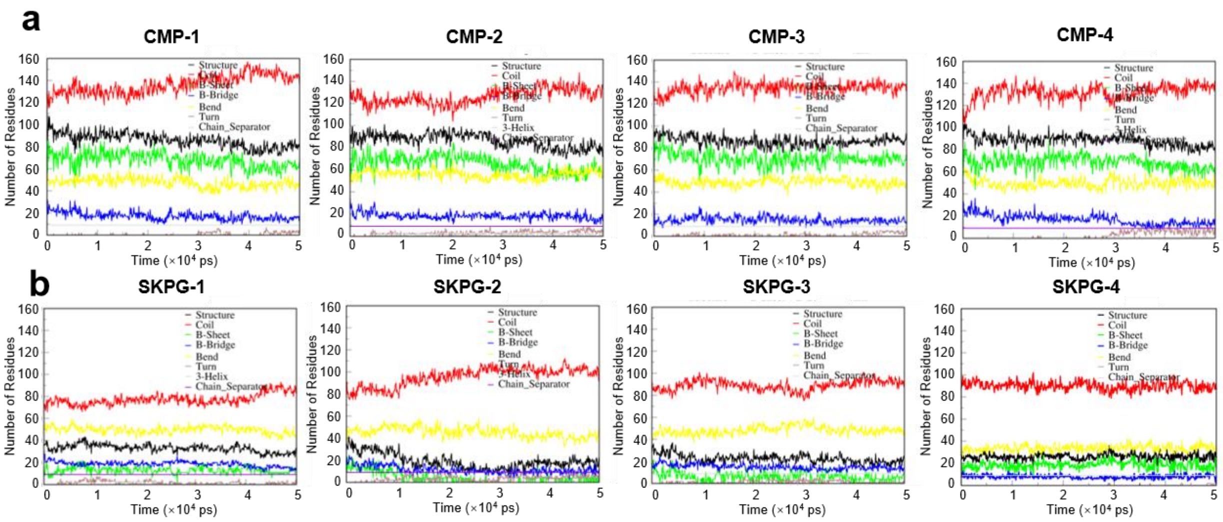

**Supplementary Fig. 26. Secondary structure analysis of CMP and SKPG peptides during AA-MD simulations.**

**a**, Secondary structure composition (coil, β-sheet, β-bridge, bend, turn, 3-helix and chain separator) and per-residue secondary structure dynamics of CMP-1/2/3/4 peptides over 50 ns. **b**, Secondary structure composition and per-residue secondary structure dynamics of SKPG-1/2/3/4 peptides over 50 ns.

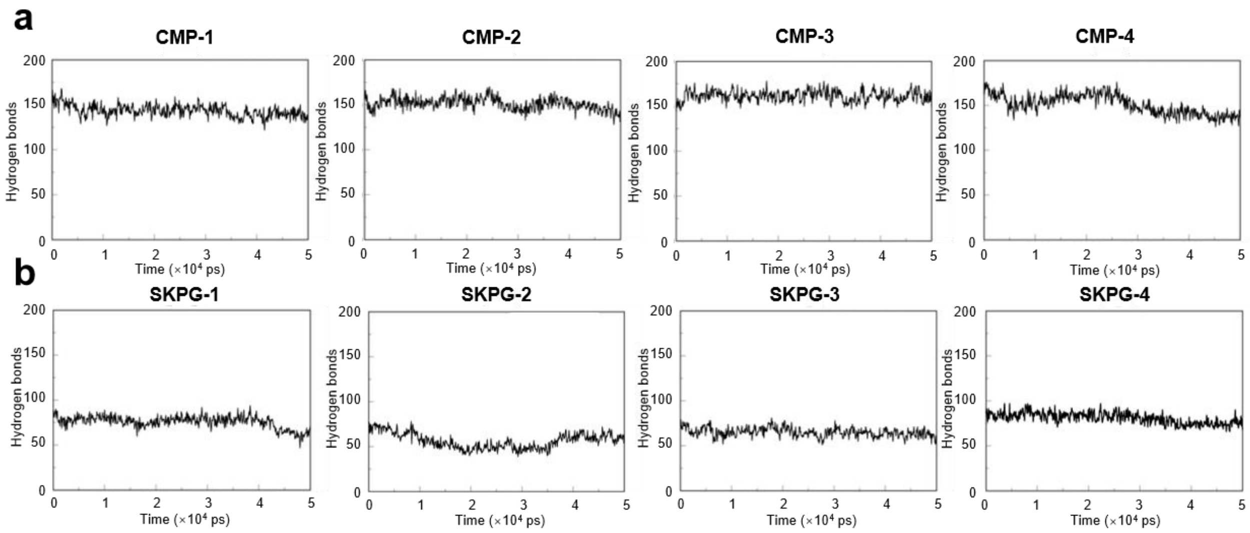

**Supplementary Fig. 27. Hydrogen bond dynamics of CMP and SKPG peptides over 50 ns AA-MD simulations.**

**a**, Time evolution of hydrogen bond counts within CMP- 1/2/3/4 assemblies over 50 ns. **b**, Time evolution of hydrogen bond counts within SKPG- 1/2/3/4 assemblies over 50 ns.

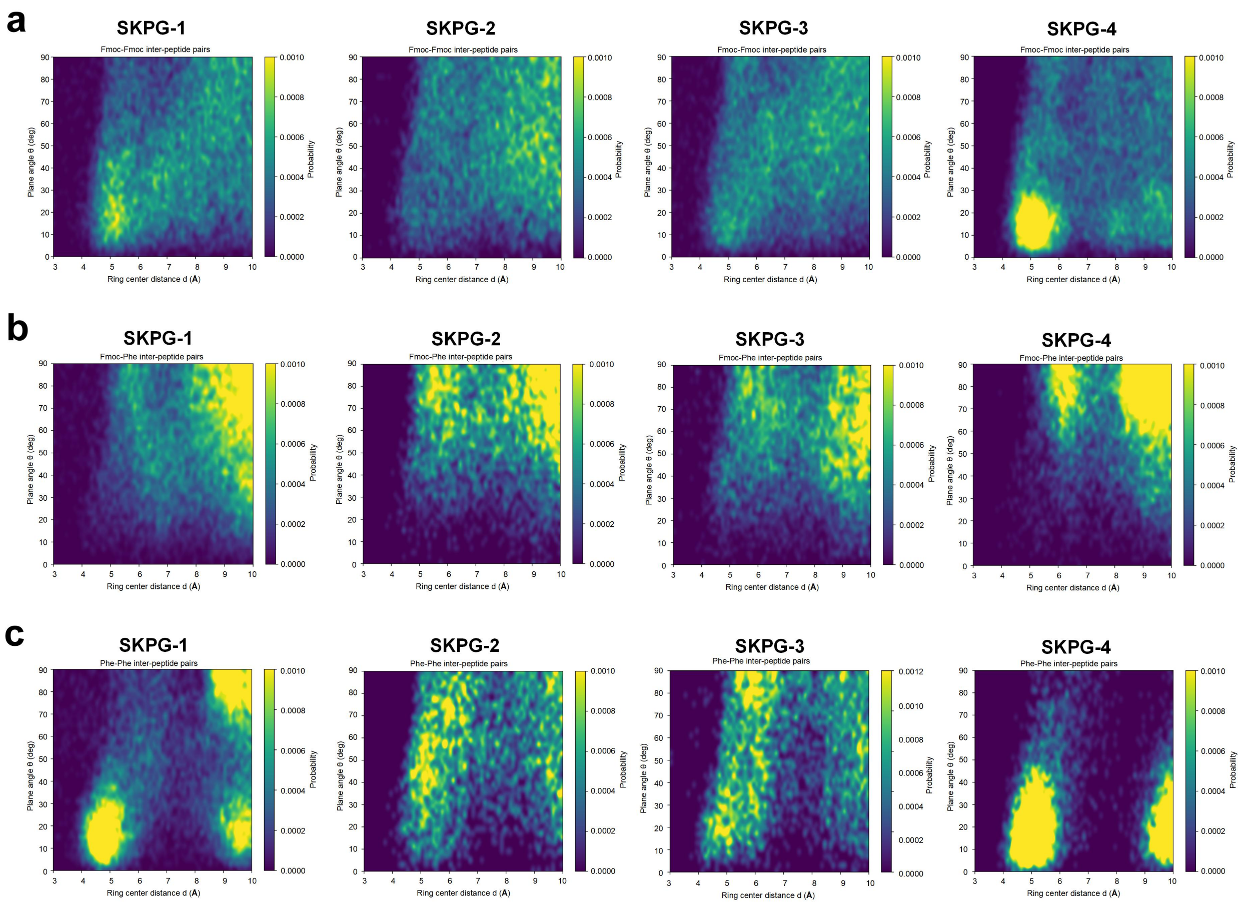

**Supplementary Fig. 28. Free-energy landscapes of SKPG peptide self-assembly as a function of centroid distance and dihedral angle between aromatic ring pairs from AA-MD simulations.**

**a**, Free-energy landscape of SKPG-1/2/3/4 peptides between Fmoc-Fmoc aromatic ring pairs. **b**, Free-energy landscape of SKPG-1/2/3/4 peptides between Fmoc-Phe aromatic ring pairs. **c**, The Free-energy landscape of SKPG-1/2/3/4 peptides between Phe-Phe aromatic ring pairs.

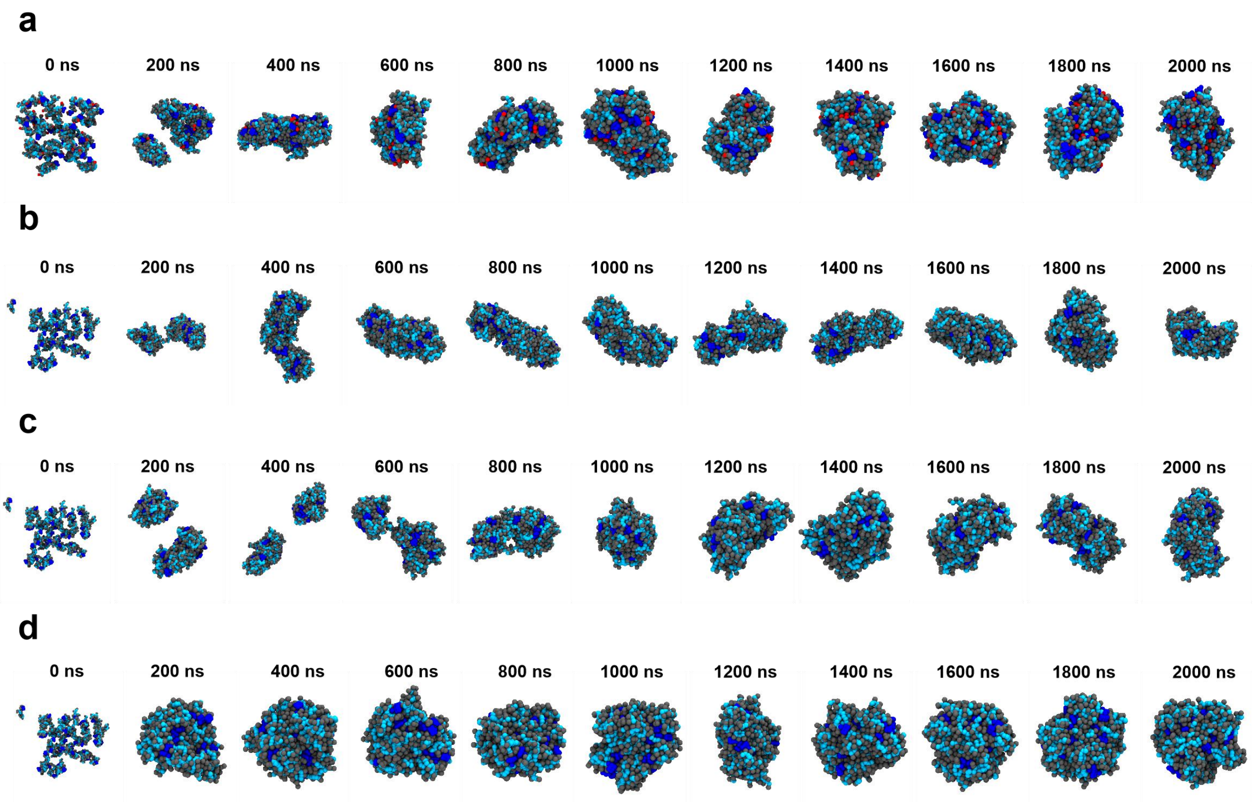

**Supplementary Fig. 29. CG-MD simulations of SKPG peptide self-assembly in aqueous solution.** All simulations were initiated from disordered states. **a**, Representative snapshots of SKPG-1 self-assembly over 2000 ns CG-MD. **b**, Representative snapshots of SKPG-2 self-assembly over 2000 ns CG-MD. **c**, Representative snapshots of SKPG-3 self-assembly over 2000 ns CG-MD. **d**, Representative snapshots of SKPG-4 self-assembly over 2000 ns CG-MD.

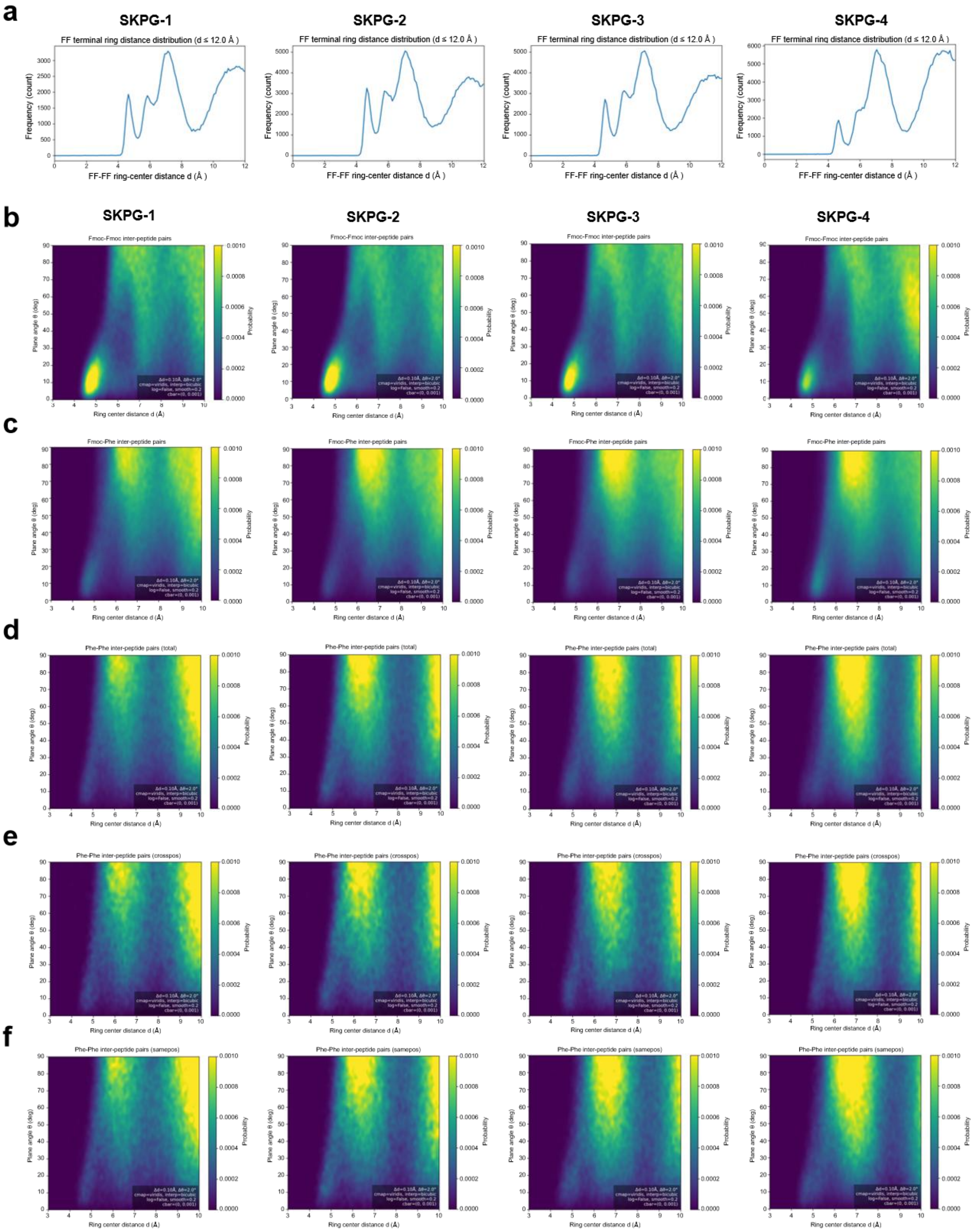

**Supplementary Fig. 30. Free-energy landscapes and end-to-end distance distributions of SKPG peptide self-assembly from CG-MD simulations.**

**a**, Probability density function of end-to-end distances for SKPG- 1/2/3/4 peptides. **b**, Free-energy landscape of SKPG-1/2/3/4 peptides between Fmoc-Fmoc aromatic ring pairs. **c**, Free-energy landscape of SKPG-1/2/3/4 peptides between Fmoc-Phe aromatic ring pairs. **d**, Free-energy landscape of SKPG-1/2/3/4 peptides between Phe-Phe aromatic ring pairs. **e**, Free-energy landscape of SKPG-1/2/3/4 peptides between cross- position Phe-Phe aromatic ring pairs. **f**, Free-energy landscape of SKPG-1/2/3/4 peptides between same-position Phe-Phe aromatic ring pairs.

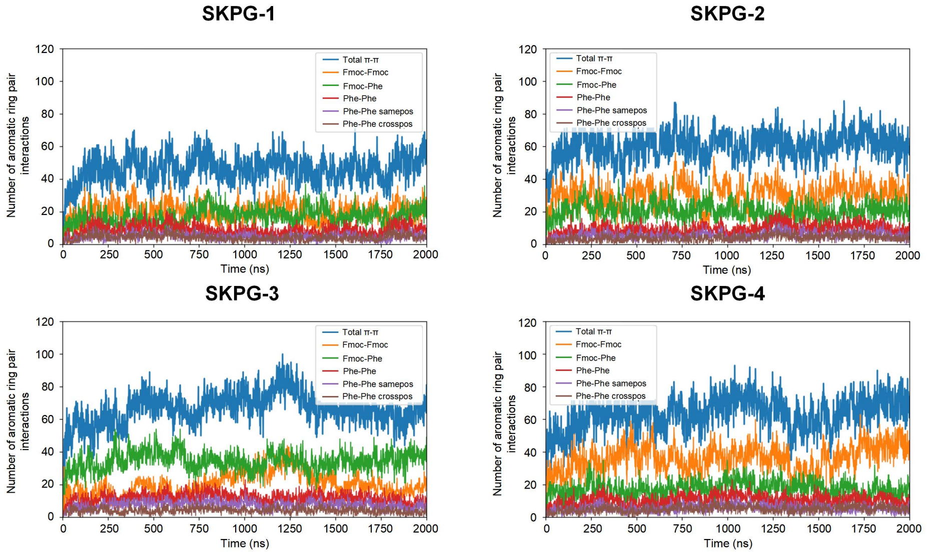

**Supplementary Fig. 31. Statistical counts of aromatic ring pair interactions of SKPG peptides over 2000 ns CG-MD simulations.**

Statistical counts of total, Fmoc-Fmoc, Phe-Phe, same-position Phe-Phe and cross-position Phe-Phe aromatic ring pairs in SKPG-1/2/3/4 peptides.

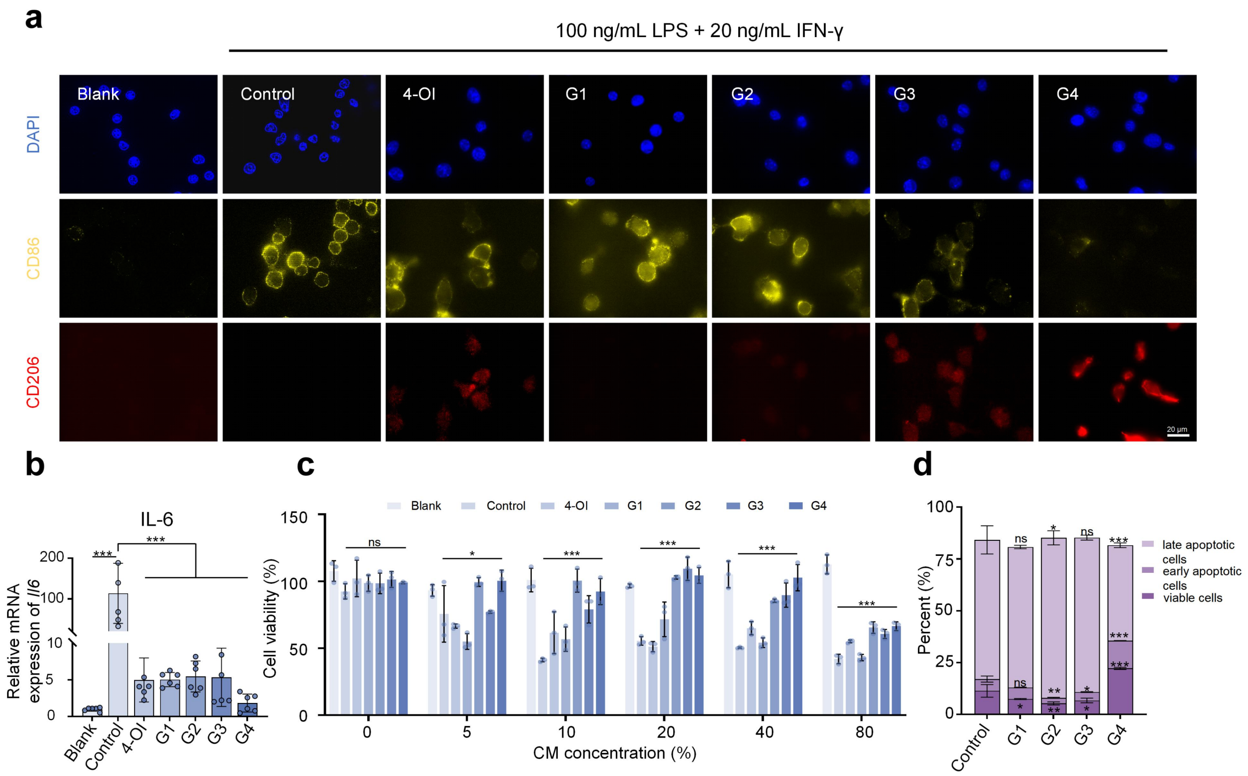

**Supplementary Fig. 32. Macrophage polarization, oxidative stress and chondrocyte protection by G4 (related to Fig. 6).**

**a**, CLSM images of CD86 and CD206 immunostaining in macrophages treated with different formulations (single-channel views). Scale bar: 20 μm. **b**, qRT- PCR analysis of IL-6 mRNA expression in macrophages treated with different formulations, n = 6. **c**, Cell viability of ATDC-5 chondrocytes cultured with conditioned media from macrophages treated with different hydrogel formulations, determined by CCK-8 assay, n = 3. **d**, Flow cytometry analysis and quantification of apoptosis in ATDC-5 chondrocytes after culture with conditioned media from different treatment groups, assessed by Annexin V-FITC/PI staining. Data are presented as mean ± SD. Statistical significance was determined by one-way ANOVA. ns, not significant; *$p < 0.05$, **$p < 0.01$, ***$p < 0.001$, ****$p < 0.0001$.

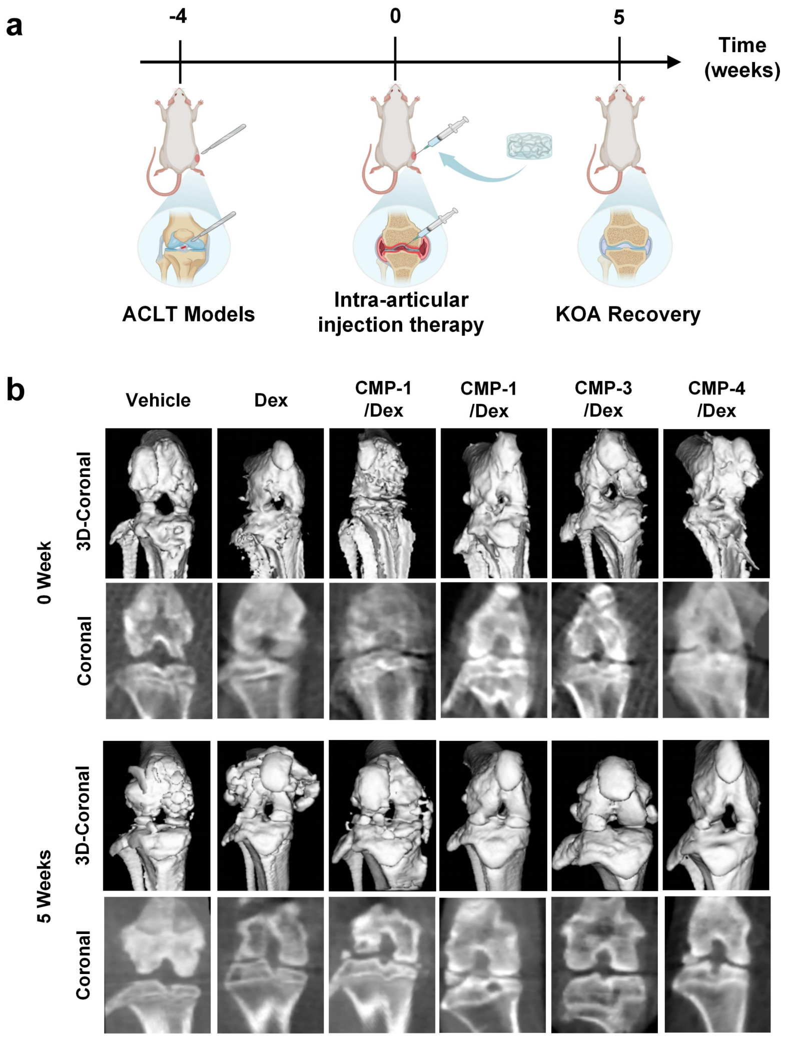

**Supplementary Fig. 33. CMP-based smart delivery system with dexamethasone for ACLT-induced OA treatment.**

**a**, Schematic diagram of the ACLT modeling and treatment timeline. **b**, Three-dimensional and two-dimensional CT reconstructions of rat knee joints before and after treatment (0 and 5 weeks).

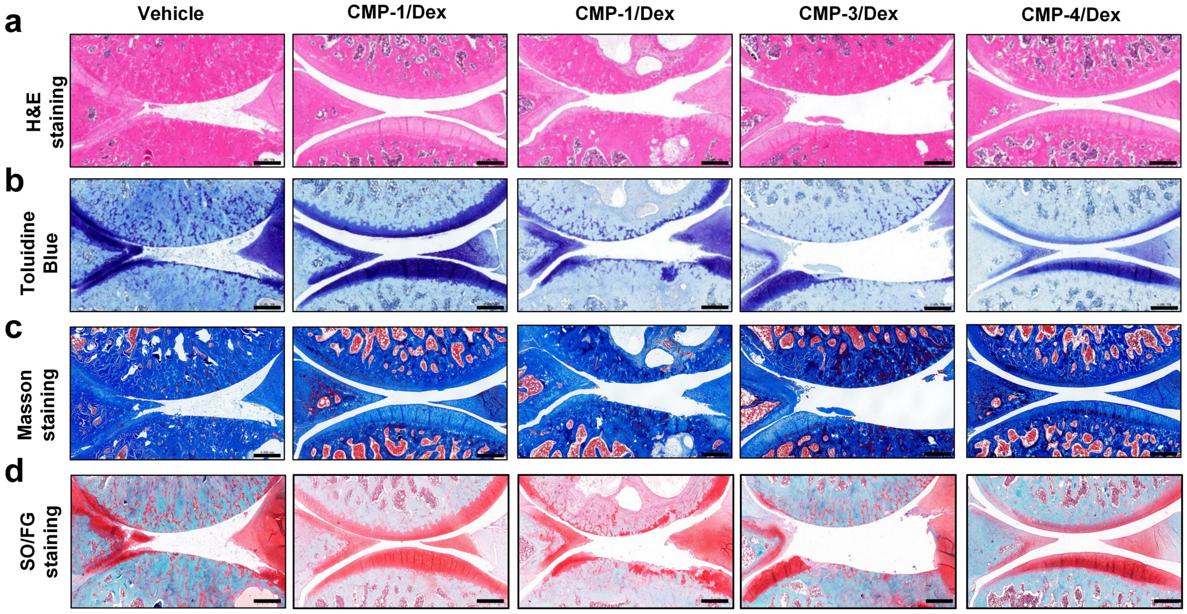

**Supplementary Fig. 34. Histological analysis of knee joint sections from ACLT-induced OA rats treated with CMP/Dex system.**

**a,** H&E staining. **b,** Toluidine blue staining. **c,** Masson staining. **d,** SO-FG staining. Scale bars: 0.5 mm.

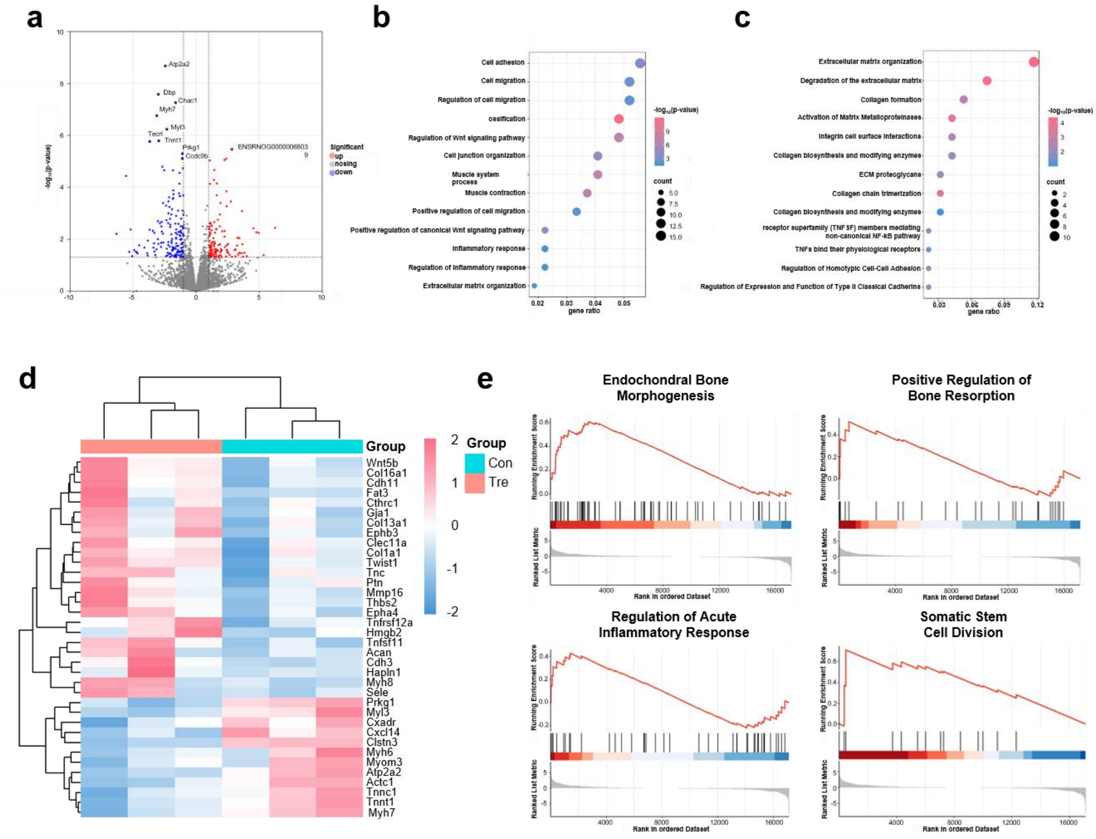

**Supplementary Fig. 35. Transcriptomic analysis of CMP-4/Dex-treated versus PBS-treated ACLT-induced**

**OA rat knees.**

**a**, Volcano plot of differentially expressed genes (DEGs) between the treatment (CMP-4/Dex) and control (PBS) groups. DEGs were defined with cutoffs of FDR < 0.05 and $|\log_2$ fold change$| > 1$. **b**, Gene Ontology (GO) pathway enrichment analysis of DEGs. **c**, Reactome pathway enrichment analysis of DEGs. **d**, Heatmap of the transcriptomic profiles of rat knee joints injected with PBS or CMP-4/Dex (n = 3). **e**, GSEA of endochondral bone morphogenesis, positive regulation of bone resorption, regulation of acute inflammatory response and somatic stem cell division between the treatment and control groups.

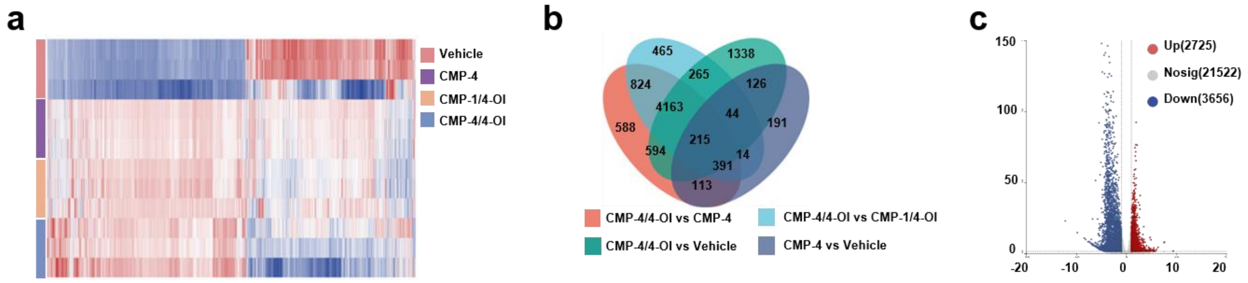

**Supplementary Fig. 36. Transcriptomic profiling of rat knee joints treated with Vehicle, CMP-4, CMP-1/4-OI and CMP-4/4-OI.**

**a**, Heatmap of gene expression profiles in rat knee joints injected with PBS (Vehicle), CMP-4, CMP-1/4-OI and CMP-4/4-OI (n = 3 per group). **b**, Venn diagram showing the overlap of differentially expressed genes among the four treatment groups. **c**, Volcano plot of DEGs between the CMP-1/4-OI and CMP-4/4-OI groups. DEGs were defined with cutoffs of FDR < 0.05 and $|\log_2$ fold change$| > 1$.

**Supplementary table S1. List of qPCR primers.**

| | | |
|---|---|---|
| Human-*COL2A1* | F | CCTGGCAAAGATGGTGAGACAG |
| | R | CCTGGTTTTCCACCTTCACCTG |
| Human-*GAPDH* | F | GTCTCCTCTGACTTCAACAGCG |
| | R | ACCACCCTGTTGCTGTAGCCAA |
| Human-*SOX9* | F | CGAGCACTCGGGGCAATC |
| | R | GAAGTCGATAGGGGGCTGTC |
| Human-*PIEZO1* | F | ATCGCCATCATCTGGTTCCC |
| | R | TGGTGAACAGCGGCTCATAG |

| | | |
|---|---|---|
| Human-*PIEZO2* | F | ACTGGACACCATTGACGAGC |
| | R | TTCAGTGTAGCAGCTGGAGAT |
| Human-*PIK3C3* | F | CCAGTGAGAACATCCTACAAAGC |
| | R | CCTTGGCGAAACATGCCGTA |
| Human-*AKT1* | F | CTCTTTCCAGACCCACGACC |
| | R | ACAGGTGGAAGAACAGCTCG |
| Mouse-*Il1b* | F | TGCCACCTTTTGACAGTGATG |
| | R | CATCTCGGAGCCTGTAGTGC |
| Mouse-*Il6* | F | GCCTTCTTGGGACTGATGCT |
| | R | TGTGACTCCAGCTTATCTCTTGG |
| Mouse-*Tnf* | F | ACCCTCACACTCACAAACCA |
| | R | CCGGACTCCGCAAAGTCTAA |
| Mouse-*Il10* | F | GGCCCAGAAATCAAGGAGCA |
| | R | AGACACCTTGGTCTTGGAGCTTAT |
| Mouse-*Il4* | F | GGTCTCAACCCCCAGCTAGT |
| | R | GCCGATGATCTCTCTCAAGTGAT |
| Rat-*Gapdh* | F | ACCATCTTCCAGGAGCGAGA |
| | R | CTCGTGGTTCACACCCATCA |
| Rat-*Timp1* | F | GAATGTGCATGACGGAGCTG |
| | R | GCGCCATCGTGGTATCTCTA |
| Rat-*Itgb2* | F | CACTCGGCACAGAAGACGTA |
| | R | AGGAGGGTTTCGAGGGTTCT |
| Rat-*Il6r* | F | CAACGAGTGTCCTCGCCCC |
| | R | CAGAGCTGTTATGGGACCCTG |
| Rat-*Vav3* | F | CAGGCAACAGCTTGCTAAGT |
| | R | AATGGCGTGCAGGTTTCTGT |
| Rat-*Ptk2b* | F | CTTGCCCAGCCCGGAG |
| | R | CGGACATCCTCTCAGACTGC |
| Rat-*Mmp8* | F | ACCAATGCTGGAGATACGACA |

| | | |
|---|---|---|
| | R | ACGCTTGCTATGCTAGTGGG |
| Rat-*Uqcrfs1* | F | AGTGGGCCTGAATGTTCCTG |
| | R | TATGGCGCACAAACAGAGGT |
| Rat-*Adipoq* | F | AGAAGGGAGACGCAGGTGTTC |
| | R | TACACTTGGAGCCAGACTTGG |
| Rat-*Irf4* | F | TTATGGCTCTCTGCCAACCC |
| | R | CCTGTCACCTGGCAACCATT |
| Rat-*Tgfb2* | F | TCCCCTCCGAAAATGCCATC |
| | R | GAGACATCGAAGCGGACGAT |
